# Microenvironmental arginine restriction sensitizes pancreatic cancers to polyunsaturated fatty acids by suppression of lipid synthesis

**DOI:** 10.1101/2025.03.10.642426

**Authors:** Patrick B. Jonker, Mumina Sadullozoda, Guillaume Cognet, Juan J. Apiz Saab, Kelly H. Sokol, Violet X. Wu, Deepa Kumari, Colin Sheehan, Mete E. Ozgurses, Darby Agovino, Grace Croley, Lindsey N. Dzierozynski, Chufan Cai, Leah M. Ziolkowski, Smit A. Patel, Althea Bock-Hughes, Kay F. Macleod, Hardik Shah, Jonathan L. Coloff, Evan C. Lien, Alexander Muir

## Abstract

Nutrient limitation is a characteristic feature of poorly perfused tumors. In contrast to well-perfused tissues, nutrient deficits in tumors impose metabolic constraints on cancer cells. The metabolic constraints created by the tumor microenvironment can lead to vulnerabilities in cancers. Identifying the metabolic constraints of the tumor microenvironment and the vulnerabilities that arise in cancers can provide new insight into tumor biology and identify promising antineoplastic targets. To identify how the microenvironment constrains the metabolism of pancreatic tumors, we challenged pancreatic cancer cells with microenvironmental nutrient levels and analyzed changes in cellular metabolism. We found that arginine limitation in pancreatic tumors perturbs saturated and monounsaturated fatty acid synthesis by suppressing the lipogenic transcription factor SREBP1, in part via activation of the amino acid sensor GCN2. Synthesis of these fatty acids is critical for maintaining a balance of saturated, monounsaturated, and polyunsaturated fatty acids in cellular membranes. Because of microenvironmental constraints on fatty acid synthesis, pancreatic cancer cells and tumors are unable to maintain lipid homeostasis when exposed to polyunsaturated fatty acids, leading to cell death by ferroptosis. In sum, arginine restriction in the tumor microenvironment constrains lipid metabolism in pancreatic cancers, which renders these tumors vulnerable to polyunsaturated-enriched fats.

## Introduction

Nutrients are delivered to tissues via perfusion from the circulatory system. In healthy tissues, perfusion delivers nutrients and removes waste at a rate that meets the metabolic demands of the tissue, resulting in tissue microenvironments that are in metabolic equilibrium with the circulatory system^1,2^. In contrast to healthy tissues, many tumors have abnormal and dysfunctional vasculature^3–7^. This is especially true for pancreatic ductal adenocarcinomas (PDAC), which are amongst the least vascularized tumors^8^. The limited blood vessels that do infiltrate PDAC tumors are largely collapsed and non-functional^9–11^. This dysfunctional vasculature, coupled with high metabolic activity of malignant and stromal cells in tumors, results in a tumor microenvironment (TME) that is not in metabolic equilibrium with the circulation and has substantially different nutrient levels than healthy tissues. Analysis of tumor interstitial fluid (TIF), the local perfusate of the TME, confirms that metabolite availability in the TME of many solid tumors is substantially different from metabolite levels in the circulatory system and healthy tissues^12–14^. Thus, abnormal vasculature limits perfusion in tumors, resulting in abnormal nutrient availability in the TME.

Limited perfusion of the TME alters cancer cell metabolism by subjecting cancer cells to nutrient limitation^14–16^. Cancer cells must adapt to these metabolic constraints to survive and proliferate in the TME^14–16^. Since healthy tissues do not face these constraints or require similar adaptive mechanisms, such adaptations could be novel anti-neoplastic drug targets that may be targetable with a wide therapeutic index^12,17–19^. As a result, there is a significant interest in understanding how cancers adapt to TME metabolic stresses. However, both the metabolic landscape of the TME and the consequent adaptations by tumors are poorly defined.

We sought to identify how abnormal nutrient levels in the TME constrain the metabolism of PDAC tumors. To do so, we measured nutrient concentrations in TIF from murine models of PDAC^20,21^. We then developed a cell culture medium termed Tumor Interstitial Fluid Medium (TIFM) that contains PDAC TIF levels of nutrients^22^. This medium allows ex vivo culture and analysis of cells challenged with TME nutrient constraints. Additionally, as the medium is chemically defined, TIFM enables identification of specific nutrient stresses that drive phenotypes in malignant PDAC^22^ or stromal cells^23^.

In this study, we performed transcriptomic and metabolic analysis of PDAC cells in TIFM and animal models of PDAC. We found that nutrient stress in the TME inhibits fatty acid synthesis so acutely that it causes PDAC cells to rely on exogenous fatty acids for growth. Mechanistically, we found arginine restriction inhibits fatty acid synthesis by suppression of the lipogenic transcription factor SREBP1, in part via activation of the amino acid sensor GCN2. Furthermore, we found these TME constraints on fatty acid synthesis challenge the ability of PDAC to maintain lipid homeostasis. Cells must tightly regulate the saturation of fatty acids in cellular lipids to ensure homeostasis^24^. Fatty acid synthesis generates saturated and monounsaturated fatty acids that allow cells to prevent toxic fatty acid imbalances^25–30^, which can be brought upon by exposure to exogenous fatty acid sources enriched in either highly saturated^28,29^ or polyunsaturated fats^25,30,31^. We found that inhibition of fatty acid synthesis, due to arginine deprivation, renders both cultured PDAC cells and tumors incapable of buffering their lipidome from exposure to exogenous polyunsaturated fatty acids. Collectively, this work delineates how arginine limitation, a metabolic limitation of the TME, constrains PDAC fatty acid synthesis, rendering these tumors vulnerable to dietary interventions that alter lipid saturation in tumors.

### Physiological nutrient availability perturbs *de novo* lipogenesis in PDAC cancer cells by reducing SREBP1 protein expression

To study how nutrient availability in the TME impacts PDAC biology, we previously developed Tumor Interstitial Fluid Medium (TIFM) that enables culture of cells in nutrient levels representative of the murine PDAC TME^22^. TIFM contains the same base electrolyte mix, bicarbonate buffer and dialyzed fetal bovine serum (FBS) supplementation as the standard cell culture media RPMI-1640. However, TIFM replaces the metabolites present in RPMI-1640 with 115 metabolites at the concentrations measured in PDAC interstitial fluid^20^ (Supplementary file 1). Thus, TIFM recapitulates the metabolic content of the PDAC TME, while containing the same base components as the standard cell culture media RPMI-1640. We then derived paired cell lines from murine *LSL-Kras^G12D/+^; Trp53^flox/flox^; Pdx-1-Cre*^32^ tumors in standard cell culture medium (RPMI-1640; herein RPMI) or TIFM, termed mPDAC-RPMI or mPDAC-TIFM cell lines^22^ (Fig. 1A).

**Figure 1:**
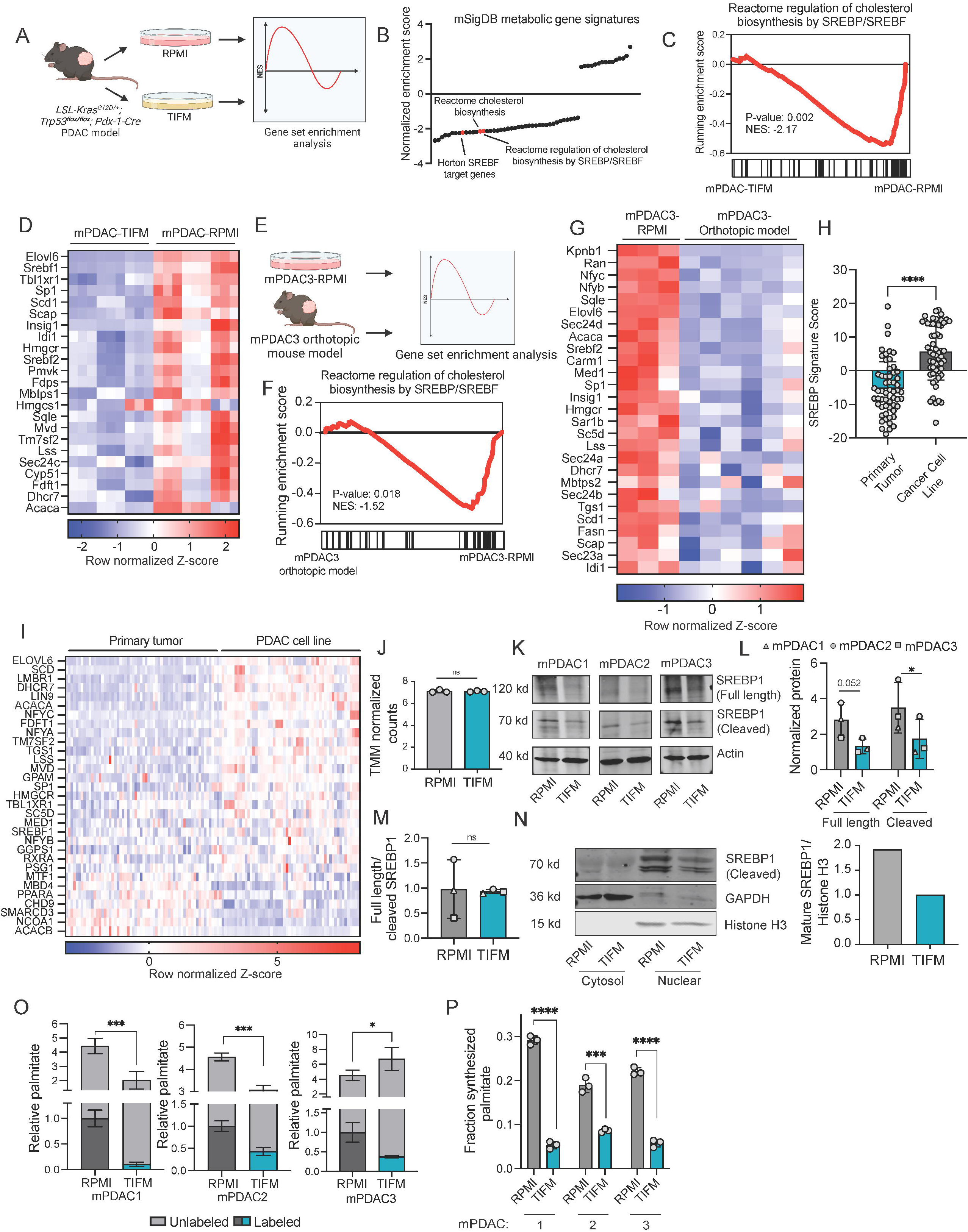
TME nutrient stress impair SREBP1 and fatty acid synthesis in PDAC. **(A)** Diagram of workflow for the transcriptomic comparison of mPDAC cells grown in TIFM or RPMI. Three pairs of cell lines (mPDAC1, mPDAC2 and mPDAC3) derived into TIFM and RPMI were analyzed with a sample size of n = 3 for each cell line. **(B)** Plotted normalized enrichment scores of metabolic gene sets from the C2 mSigDB database set. SREBP-related gene sets are highlighted in red. **(C)** Gene Set Enrichment Analysis (GSEA) of the *Reactome regulation of cholesterol biosynthesis by SREBP/SREBF* gene set in transcriptomic data of mPDAC cell lines cultured in TIFM or RPMI. **(D)** Row-scaled heatmap of leading edge genes from the GSEA analysis in **(C)**. Z-scores calculated from trimmed mean of *M* values (TMM) gene counts for Reactome regulation of cholesterol biosynthesis by SREBP/SREBF genes in mPDAC cells cultured in TIFM versus RPMI. **(E)** Diagram of workflow for the transcriptomic comparison of mPDAC3 cells grown in RPMI (*n* = 3) or as orthotopic allograft murine tumors (*n* = 6). mPDAC3 cells from each condition were isolated by fluorescence-activated cell sorting (FACS) and RNA was isolated for transcriptomic analysis by next generation sequencing. **(F)** GSEA of the *Reactome regulation of cholesterol biosynthesis by SREBP/SREBF* gene set using transcriptomic data of mPDAC3 cell lines cultured in RPMI or orthotopically implanted in C57BL/6 mice. **(G)** Row-scaled heatmap of leading edge genes from the GSEA analysis in **(F)**. Z-scores calculated from trimmed mean of *M* values (TMM) gene counts for *Reactome regulation of cholesterol biosynthesis by SREBP/SREBF* genes in mPDAC3 cells cultured in RPMI or isolated from orthotopic tumors. **(H)** SREBP signatures for human PDAC tumors and PDAC cancer cell lines generated using publicly available transcriptomic data using MERAV. Each point indicates a tumor specimen or cell line. **(I)** Row normalized heatmap z-scores of human transcriptomic data of genes from the *Reactome regulation of cholesterol biosynthesis by SREBP/SREBF* gene set. Genes shown are significantly differentially expressed between patient samples and PDAC cell lines with p-value cutoff of p<0.05. **(J)** TMM normalized counts from transcriptomic analysis of mPDAC3 cells grown in TIFM or in RPMI (n=3). **(K)** Immunoblot analysis of full length and cleaved SREBP1 expression in mPDAC1 and mPDAC3 cell lines grown in TIFM or RPMI treated with 10 µM MG132. **(L)** Quantification of immunoblot signal of full length and proteolytically processed SREBP1 normalized to actin. **(M)** Quantification of immunoblot signal of proteolytically processed SREBP1 divided by immunoblot signal of full length SREBP1. This analysis was based on triplicate analysis of each cell line in each condition. **(N)** Nuclear extraction of mPDAC1 cells cultured in TIFM of RPMI. Blot stained for SREBP1, with GAPDH as a control for cytosol and Histone H3 as a control for nuclear protein (left). Quantification of mature SREBP1 normalized to Histone H3 (right). **(O)** Levels of unlabeled (grey) and deuterium labeled (blue, dark grey) palmitate (16:0) in mPDAC cell lines grown in TIFM or RPMI. Labeled palmitate indicates de novo synthesized lipid (n=3). **(P)** Fraction of synthesized palmitate to total palmitate in mPDAC1, 2, and 3 cells grown in TIFM or in RPMI (n=3). Statistical significance was assessed using two-tailed Welch’s t-test in H, J, M, O, and P. Statistical significance was assessed using a paired t test in L. Statistical significance was assessed using the weighted Kolmogorov–Smirnov statistic in C, F.

We performed transcriptomic analysis of murine PDAC cell lines cultured in TIFM or RPMI cells to identify pathways altered by nutrient availability in the TME in PDAC cells. We used two pathway analyses to discern which metabolic pathways were affected by culture in TIFM: gene set enrichment analysis (GSEA^33^) and shiny GATOM^34^, which identifies enrichment of differentially expressed genes in KEGG annotated metabolic pathways. Gene set enrichment analysis (GSEA) identified significant changes in a number of metabolic pathways in mPDAC-TIFM cells (Fig. 1B, Fig. 1 – Figure supplement 1A). Multiple gene sets related to the transcription factor SREBP, which regulates lipid and sterol metabolism^35^, were among the most downregulated metabolic gene sets in TIFM cultured cells (Fig. 1B-D). Similarly, GATOM analysis revealed that lipid metabolic pathways were the most significantly downregulated among KEGG metabolic pathways between mPDAC1-TIFM and mPDAC1-RPMI cells (Fig. 1 – Figure supplement 1B). Analysis of transcriptomic data from mPDAC3-RPMI cells either cultured in standard media or sorted from orthotopic allograft tumors (Fig. 1E)^22^ showed that SREBP target gene expression is similarly downregulated in PDAC cancer cells in vivo (Fig. 1F,G). Finally, we performed reverse-phase protein array analysis (RPPA)^36^ to identify changes in protein level and/or modification that occur in mPDAC cells upon culture in TIFM. We found that several SREBP1 target genes were downregulated at the protein level in TIFM cultured mPDAC cells (Fig. 1 – Figure supplement 1C). Together, these data indicate that nutrient availability in the TME, modeled in TIFM, suppresses SREBP target gene expression. To see if the TME suppresses SREBP target gene expression in human tumors, we used MERAV^37^ to compare transcriptomic data from primary PDAC specimens and human PDAC cell lines. We found that PDAC tumors exhibit markedly lower SREBP target gene expression than that of PDAC cell lines grown in vitro (Fig. 1H,I). SREBP targets and lipid synthesis genes are also downregulated in murine PDAC compared to healthy pancreas^38^ and analysis of Human Protein Atlas data indicates that protein expression of SREBP target genes is decreased in PDAC compared to normal pancreases (Fig. 1 – Figure supplement 1D). Altogether, these data suggest that the TME suppresses SREBP target gene expression in murine and human PDAC, and that TIF nutrient availability is sufficient to downregulate expression of SREBP target genes.

SREBP transcription factors act as master regulators of lipid synthesis by facilitating the transcription of genes important for fatty acid and sterol biosynthesis^35^. There are two major isoforms of SREBP: SREBP1, which primarily controls fatty acid synthesis by enhancing the transcription of fatty acid synthesis proteins, and SREBP2, which mainly controls expression proteins involved in sterol synthesis and uptake. Analysis of SREBP1-specific and SREBP2-specific transcripts in mPDAC cells indicated that SREBP1 targets were specifically downregulated by TIFM culture (Fig. 1 – Figure supplement 2A,B). Thus, we focused our studies on SREBP1 regulation in the PDAC TME.

SREBP activity is regulated both by modulation of SREBP1 expression, proteolytic processing of SREBP to the mature transcription factor and translocation of mature SREBP to the nucleus^35^. To determine how TIFM inhibits SREBP1, we performed transcriptomic and immunoblotting analysis of SREBP1 in mPDAC cells cultured in TIFM and RPMI. We did not find differences in expression of SREBP1 transcript (Fig. 1J) between TIFM and RPMI cultured mPDAC cells, indicating that transcriptional regulation does not account for differences in SREBP1 activity between culture conditions. At the protein level, both full length and mature SREBP1 protein levels were significantly lower in TIFM-cultured mPDAC cells and human PDAC cell lines in TIFM (Fig. 1K,L, Fig.1 – Figure supplement 3A). In contrast, the ratio of mature to full length SREBP1 (a proxy of proteolytic activation of SREBP1) was not significantly altered between TIFM and RPMI culture conditions (Fig. 1M). Lastly, to assess whether nuclear translocation was altered by nutrient availability, we extracted nuclear and cytosolic protein from mPDAC1-TIFM and mPDAC1-RPMI cells and compared levels of nuclear SREBP1. We found that nuclear levels of SREBP1 in TIFM-cultured cells were reduced, but to a similar extent as occurred at the whole cell level (Fig. 1N), indicating that nuclear translocation of SREBP1 is unaffected by TIFM culture. Thus, the major mechanism by which nutrient stress in the TME suppresses SREBP1 in murine and human PDAC is by regulating SREBP1 protein expression.

Lastly, given the suppression of SREBP1, we assessed whether TIFM culture suppresses fatty acid biosynthesis. To do this, we performed deuterium stable isotope labeling of fatty acids in cells cultured in TIFM or in standard cell culture media^39^. We found that mPDAC cells cultured in TIFM synthesize fewer fatty acids than cells cultured in standard media (Fig. 1O), and that synthesized fatty acids represent a smaller fraction of the cellular fatty acid pool (Fig.1P). Collectively, these data indicate that physiological nutrient stress in the PDAC TME reduces SREBP1 activity and fatty acid biosynthesis in PDAC cells.

Lastly, our studies above focused on a murine model of PDAC in which loss of p53 drives tumor development^32^. PDAC is frequently driven by p53 mutations, which drive distinct microenvironmental^40,41^ and metabolic programs^42,43^. Therefore, we assessed if TIFM culture affects SREBP1 levels and fatty acid synthesis in human PDAC cells with p53 mutations. We found that SREBP1 levels and fatty acid synthesis are lowered in the human PDAC lines Aspc-1 (*KRAS^G12D^*;*SMAD4^R100T^;TP53^C135A^*; *MAP2K4*^-/-^)^44^ and Panc1 (*KRAS^G12D^*;*TP53^R273H^*;*CDKN2A*^-/-^)^44^ upon culture in TIFM (Fig. 1 – Figure supplement 3B-D). Thus, TME metabolic stress suppresses lipid synthesis in both murine and human PDAC with both loss and mutation of p53.

### Arginine restriction reduces SREBP1-mediated fatty acid synthesis in PDAC

TIFM and standard media, such as RPMI, differ both in the concentration of shared nutrients and the presence of nutrients unique to TIFM (Fig. 2 – Figure supplement 1A). We sought to identify the specific changes in nutrient availability responsible for decreased SREBP1-mediated fatty acid synthesis in PDAC. To develop a high throughput assay to assess SREBP1 expression and fatty acid synthesis in culture, we reasoned that mPDAC cells cultured in TIFM would be unable to grow without exogenous lipids, while mPDAC cells in RPMI could grow normally in the absence of exogenous lipids. To test this hypothesis, we chemically stripped lipids from FBS to make TIFM and RPMI media with depleted levels of exogenous lipids^45^. We then compared cell growth of mPDAC1-TIFM and mPDAC1-RPMI cells cultured in each medium with or without depletion of exogenous lipids. The cell line mPDAC1 was chosen because this cell line had the strongest changes in fatty acid synthesis due to TIFM culture. As predicted, we found that mPDAC1 cells deprived of exogenous lipids in TIFM grew significantly slower compared with cells cultured in RPMI (Fig. 2 – Figure supplement 1B).

We then used this assay to identify which nutrients in TIFM are responsible for perturbing fatty acid synthesis in PDAC. TIFM was constructed using separate pools of nutrient that are readily swappable with their RPMI counterparts (Fig. 2 – Figure supplement 1C). This allows us to systematically grow mPDAC cells with different pools of TME nutrients and identify the subset of nutrients responsible for a phenotype^46–49^, such as growth rate without exogenous lipids. Once a subset of phenotype-driving nutrients is identified, further subsets or individual metabolites from the pool can be assessed (Fig. 2 – Figure supplement 1D). Using this approach we found that arginine deprivation in TIFM, which is the most differentially abundant amino acid between TIFM and RPMI, limits mPDAC1 growth without exogenous lipids (Fig. 2 – Figure supplement 1E). Arginine is present in TIFM at the pathophysiologically low TME concentration of 2.27 µM while it is at the supraphysiological concentration of 1149 µM in RPMI. This led us to hypothesize that arginine deprivation in TIFM was responsible for decreased SREBP1 activity and fatty acid synthesis in PDAC cells under nutrient stress in the TME. Consistent with this hypothesis, we found that supplementing TIFM with arginine increased SREBP1 protein abundance to levels comparable to what is observed in RPMI cultures, in both mPDAC (Fig. 2A) and human PDAC cells (Fig. 2 – Figure supplement 2A). This increase in SREBP1 occurred at the protein but not mRNA level (Fig. 2 – Figure supplement 2B) and was accompanied by an increase in fatty acid synthesis (Fig. 2B, Fig. 2 – Figure supplement 2C-F). Arginine restriction also lowers SREBP1 levels (Fig. 2C,D) and fatty acid synthesis in RPMI cultured mPDAC cells (Fig. 2B). Altogether, this data suggests that arginine restriction suppresses SREBP1 and fatty acid synthesis in PDAC cells.

**Figure 2:**
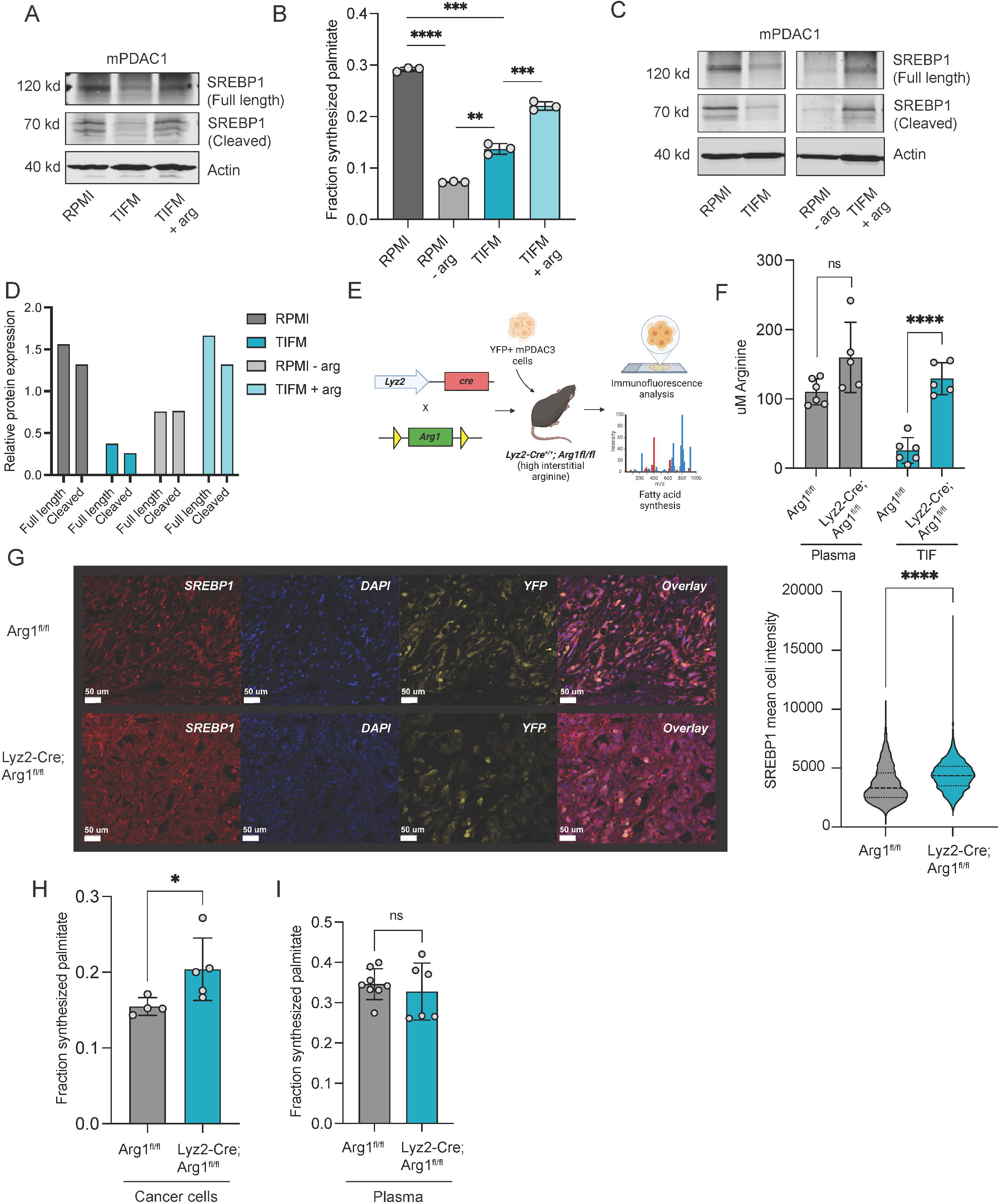
TME arginine restriction limits SREBP1 expression and fatty acid synthesis in PDAC cells and tumors. **(A)** Immunoblot analysis of full length and cleaved SREBP1 expression in mPDAC1 cells grown in TIFM, RPMI, or TIFM with supplemented arginine (TIFM + arg; 1.19 mM) treated with 10 µM MG132. **(B)** The fraction of cellular palmitate (16:0) that is deuterium labeled in mPDAC1 cell lines grown in TIFM, RPMI, TIFM + arg or RPMI with TIFM (2.27 µM) levels of arginine (n=3). **(C)** Immunoblot analysis of full length and mature SREBP1 in the same conditions as **(B)** and treated with 10 µM MG132 with quantification in **(D)**. **(E)** Schematic for generation of orthotopic tumors in *Lyz2-Cre* and *Arg1^fl/fl^* host animals and analysis of tumor fatty acid synthesis and SREBP1 protein levels. **(F)** Concentrations of arginine in TIF and plasma isolated from animals bearing orthotopic mPDAC3-TIFM tumors. n=6 for *Lyz2-Cre^+/+^; Arg1^fl/fl^* animals and n=5 for *Arg1^fl/fl^* littermate controls. **(G)** *(left)* Representative immunofluorescent images of mPDAC3-TIFM tumors from *Lyz2-Cre^+/+^; Arg1^fl/fl^* (*n* =9) and *Arg1^fl/fl^* mice (n=9). Cancer cells are YFP labeled. *(right)* Quantification of mean SREBP1 intensity for YFP positive cancer cells in *Lyz2-Cre^+/+^; Arg1^fl/fl^* animals and *Arg1^fl/fl^* littermate controls. (n=3249 cells for each genotype). **(H)** The fraction of cellular palmitate (16:0) that is deuterium labeled from mPDAC3 cells isolated from tumors from *Lyz2-Cre^+/+^; Arg1^fl/fl^* (*n* =4) and *Arg1^fl/fl^* (n=5) mice. **(I)** The fraction of plasma palmitate (16:0) that is deuterium labeled from *Lyz2-Cre^+/+^; Arg1^fl/fl^* (*n* =6) and *Arg1^fl/fl^* (n=7) mice. Significance was assessed using the Mann-Whitney U-test for H and I. Two tailed Welch’s t-test correction was used to assess significance in B, F, and G.

We next sought to determine whether arginine deprivation suppresses SREBP1 and fatty acid biosynthesis in vivo. We previously found that arginine is depleted in PDAC tumors due to arginase-1 activity from myeloid cells in the PDAC TME^22^. We established orthotopic allograft tumors by implanting YFP labeled mPDAC cells in *Lyz2-Cre^+/+^; Arg1^fl/fl^* mice, which harbor myeloid cell specific arginase-1 knockout, or control *Arg1^fl/fl^* animals (Fig. 2E). We confirmed that loss of myeloid arginase-1 increases arginine in the interstitial fluid of murine PDAC tumors to plasma levels (Fig. 2F)^47,48^. Thus, this animal model allows us to compare PDAC tumors with or without arginine restriction in the TME. We examined SREBP1 levels by immunofluorescence (IF) as previously described^50,51^ in sections from PDAC tumors from *Lyz2-Cre^+/+^; Arg1^fl/fl^* and *Arg1^fl/fl^* hosts. We found increased levels of SREBP1 in PDAC cells from arginase-1 knockout hosts (Fig. 2G). IF analysis of tissues with high and low expression of SREBP1 confirmed that the antibody and IF protocol we used yields signals proportional to tissue SREBP1 expression (Fig. 2 – Figure supplement 3A,B). To determine if arginine restriction in the TME affects fatty acid synthesis in PDAC, we administered deuterated water to PDAC bearing mice as previously described^49^. We then sorted PDAC cells from the tumors and assessed incorporation of deuterium into fatty acids, as a measure of fatty acid synthesis. We found that mPDAC cells from *Lyz2-Cre^+/+^; Arg1^fl/fl^* hosts had increased fatty acid synthesis compared to control hosts (Fig. 2H). Importantly, label incorporation in circulating fatty acids was not different between *Lyz2-Cre^+/+^; Arg1^fl/fl^* and control animals (Fig. 2I). This suggests the difference in fatty acid labeling we see in mPDAC cells from *Lyz2-Cre^+/+^; Arg1^fl/fl^* and control mice is not due to uptake of differently labeled fatty acids from the circulation. Altogether, these data show that arginine deprivation reduces the expression of SREBP1 in PDAC cancer cells both in TIFM and in vivo, leading to the suppression fatty acid synthesis.

### SREBP1 inactivation by arginine deprivation in the TME leads to changes in cellular lipidome

We observed that arginine restriction in the TME reduces fatty acid synthetic capacity in PDAC cells. As a result, we expected arginine restriction would lead to changes in the cellular lipidome. To determine if arginine affected the PDAC cellular lipidome, we performed mass spectrometry-based lipidomics analysis of mPDAC cells in TIFM, RPMI and TIFM supplemented with arginine (TIFM + arg). De novo lipogenesis generates saturated (SFA) and monounsaturated fatty acids (MUFA) in mammalian cells, while essential polyunsaturated fatty acids (PUFA) and their derivatives are primarily taken up from the cellular environment. Therefore, we reasoned that arginine deprivation, by inhibiting SREBP1 and de novo lipogenesis, would lead to an increase in the ratio of PUFAs to MUFAs (PUFA/MUFA) in PDAC cellular lipids. Consistent with this hypothesis, analysis of PUFA, MUFA, SFA abundance^52^ in the PDAC lipidome showed that TIFM cultured cells had a higher PUFA/MUFA ratio than cells cultured in standard media or TIFM + arg (Fig. 3A). Interestingly, this increase in PUFA/MUFA ratio is not apparent across all lipid species. Arginine starved PDAC cells have increased PUFA/MUFA ratio in certain lipid species (triglycerides [TG], phosphatidylethanolamines [PE], diacylglycerols [DG], ceramides [Cer], and sphingomyelin [SM]) but not others (phosphatidylcholines [PC], cholesterol esters [ChE], Hexosylceramides [Hex1-Cer], lysophosphatidylcholines [LPC] and lysophosphatidylethanolamines [LPE]) (Fig. 3B). Arginine starved PDAC cells also had other lipidomic features of impaired fatty acid synthesis. Cells with impaired fatty acid synthesis accumulate TG in lipid droplets^53^. TIFM cultured cells also had higher levels of most TG species than mPDAC cells cultured in RPMI or TIFM + arg (Fig. 3C). Consistent with the increase in TGs, TIFM cultured cells also contained more lipid droplets (Fig. 3 – Figure supplement 1). While most TG species followed this trend, ether-linked TG species were suppressed by culture in TIFM and levels of these lipids were not rescued by arginine supplementation (Fig. 3C). Thus, this class of ether-linked neutral lipids may be regulated differently by SREBP1 than other TGs. Altogether, these data indicate that arginine restriction leads to significant changes in the PDAC lipidome, which are consistent with suppression of SREBP1 and fatty acid synthesis.

**Figure 3:**
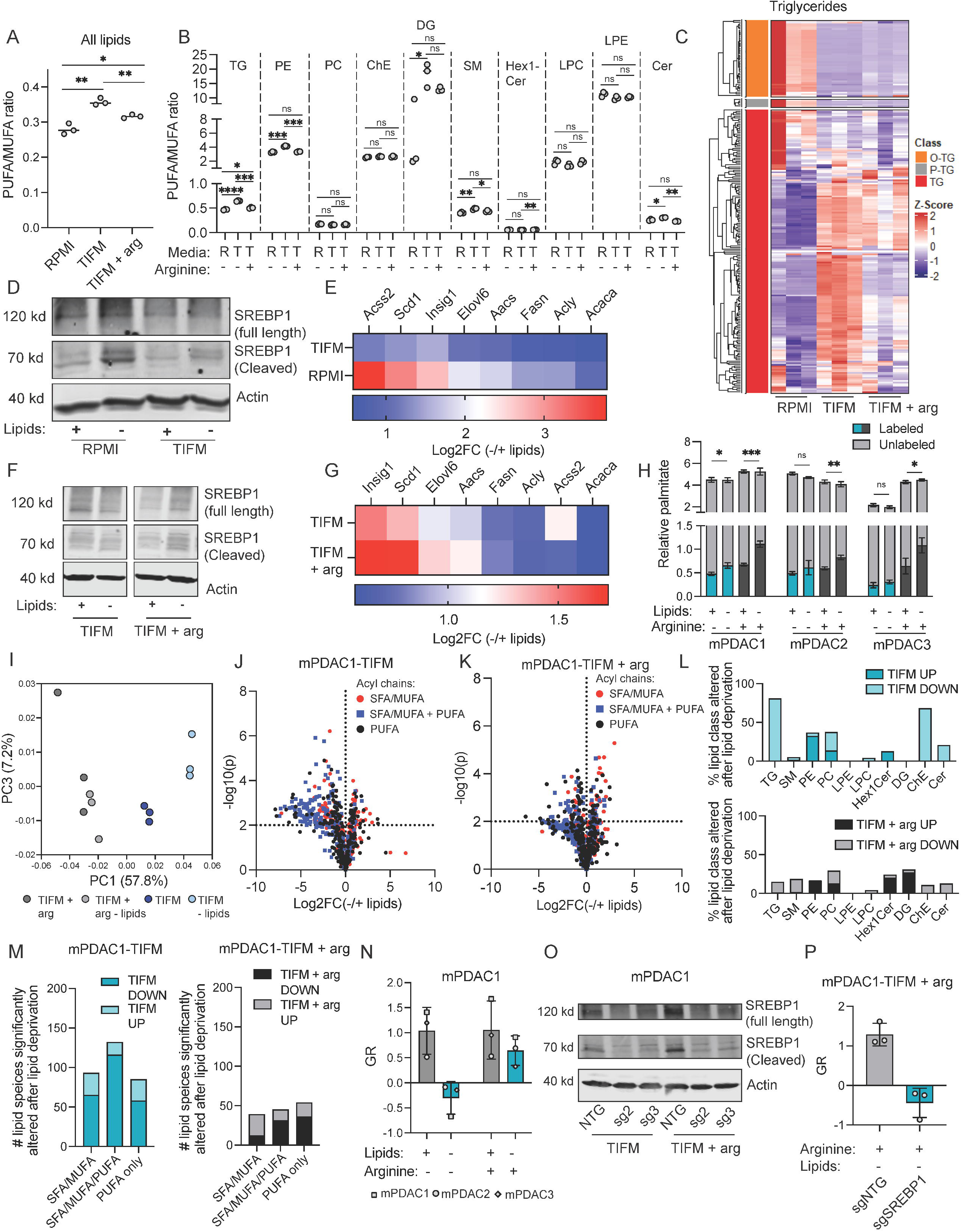
Arginine restriction prevents PDAC cells from maintaining lipid homeostasis when starved of exogenous lipids. **(A)** PUFA/MUFA ratios of all cellular lipids and **(B)** major classes of lipids species in mPDAC1 cells cultured in TIFM, RPMI or TIFM + arg. **(C)** Triglyceride (TG) levels in mPDAC1 cells cultured in TIFM, RPMI or TIFM + arg. Plotted data in **(C)** is row normalized z-scores of total-signal normalized peak areas (n=3). **(D)** Immunoblot analysis of full length and cleaved SREBP1 expression in mPDAC1 cells grown in TIFM or RPMI with and without starvation of exogenous lipid. **(E)** Transcriptomic analysis of expression of SREBP1 target genes involved in fatty acid synthesis in mPDAC1 cells cultured with exogenous lipids (+ lipids) or without exogenous lipids (-lipids) in either TIFM or RPMI. Data plotted are log2 fold change of mRNA expression between cultures (n=3). **(F)** Immunoblot analysis of mPDAC1 full length and cleaved SREBP1 expression in mPDAC1 cells grown in TIFM, RPMI or TIFM + arg with and without exogenous lipid. **(G)** Transcriptomic analysis of expression of SREBP1 target genes involved in fatty acid synthesis in mPDAC1 cells cultured with exogenous lipids (+ lipids) or without exogenous lipids (-lipids) in either TIFM or TIFM + arg. Data plotted are log2 fold change of mRNA expression between cultures (n=3).**(H)** Levels of unlabeled and deuterium labeled palmitate (16:0) in mPDAC cell lines grown in TIFM or TIFM + arg (n=3). **(I)** Principal component analysis of LC-MS measurements of lipid levels of mPDAC1 cells cultured in TIFM or TIFM + arg with or without exogenous lipids (n=3). **(J,K)** Volcano plots depicting the log2 fold change in levels of lipids in mPDAC1 cells upon starvation of exogenous lipids when cultured in TIFM **(J)** or TIFM + arg **(K)**. Lipids containing only SFA/MUFA acyl groups are highlighted in red. Lipids containing both an SFA or MUFA and PUFA acyl group are highlighted in blue. Lipids containing only PUFA acyl chains are shown in black. The cut off for statistical significance in this analysis is p<0.01. **(L)** Quantification of the percent of an indicate lipid class that is altered upon lipid deprivation in *(upper)* TIFM or *(lower)* TIFM + arg cultured mPDAC1 cells. (TG: triglycerides; SM: sphingomyelin; PE: phosphatidylethanolamine; PC: phosphatidylcholine; LPE: lysophosphatidylethanolamine; LPC: lysophosphatidylcholine; Hex1Cer: hex-1-ceramides; DG: diacyglycerol; ChE: cholesterol esters; Cer: ceramides) **(M)** Number of lipid species with indicated composition of acyl groups whose cellular abundance is significantly altered upon lipid withdrawal in *(left)* TIFM or *(right)* TIFM + arg cultured mPDAC1 cells. **(N)** Cellular growth rate inhibition (GR) of mPDAC1, 2 and 3 cell lines grown in TIFM or TIFM + arg upon lipid deprivation. Each point is plotted average of technical triplicates of the assay for each cell line (n=3). **(O)** Immunoblot analysis of SREBP1 expression in mPDAC1 cells transduced with control non-targeting or SREBP1 targeting sgRNAs. **(P)** GR of mPDAC1 cells analyzed in cultured in TIFM + arg with and without exogenous lipids (n=3). Statistical significance was assessed in all panels by two-tailed Welch’s t-tests.

### Arginine deprivation prevents PDAC cancer cells from adapting to changes in exogenous lipid availability

An important function of SREBP1 is driving transcription of lipid synthesis genes upon depletion of lipids cellular which occurs upon starvation of exogenous lipids^51^. Given that arginine restriction suppresses SREBP1 in PDAC, we asked if arginine restriction in the TME would impair the ability of mPDAC cells to cope with lipid starvation. We found mPDAC cells cultured in RPMI robustly increased SREBP1 protein levels and expression of SREBP1 target genes in response to lipid depletion (Fig. 3D,E, Fig. 3 – Figure supplement 2A,B). In contrast, mPDAC cells in TIFM had decreased activation of SREBP1 and less induction of SREBP1 target genes in lipid depleted conditions (Fig. 3D,E). Similarly, human PDAC cells in TIFM were less able to induce SREBP1 target genes upon lipid starvation compared to human PDAC cells cultured in RPMI (Fig. 3 – Figure supplement 2C). Addition of arginine to TIFM increased the induction of SREBP1 and SREBP1 target gene expression upon lipid starvation in mPDAC cells. While arginine-supplemented cells (Fig. 3G) and RPMI cultured cells (Fig. 3E) had differences in exactly which SREBP1-target genes were induced compared to TIFM cultures, key genes associated with fatty acid synthesis (FASN), desaturation (SCD1) and elongation (ELOVL6) were upregulated by both arginine supplementation and RPMI culture, suggesting arginine activate fatty acid synthesis in PDAC (Fig. 3F,G). Consistent with arginine restriction impairing SREBP1 induction upon lipid starvation, we found that induction of fatty acid synthesis upon lipid starvation was blunted in TIFM cultured cells and rescued by the addition of arginine to culture media (Fig. 3H). These data reveal that arginine restriction prevents PDAC cells from responding to changes in lipid availability via SREBP1 signaling and induction of de novo lipid synthesis.

We next asked if impaired SREBP1 signaling and de novo lipogenesis would prevent arginine-restricted PDAC cells from maintaining lipid homeostasis under lipid starvation. To ask this, we deprived PDAC cells of lipids for 24 hours and performed lipidomic analysis. The lipidome of TIFM cultured cells was perturbed by lipid starvation, as lipid replete and lipid starved samples clustered separately in principal component analysis (Fig. 3I). In contrast, lipid starvation did not lead to separate clustering in PDAC cells cultured with supplemental arginine (Fig. 3I). Analysis of individual lipid species also indicated that more lipid species were significantly depleted upon lipid starvation in arginine-restricted cultures compared to arginine replete (Fig. 3J-L) and standard media cultures (Fig. 3 – Figure supplement 3A,B). Neutral lipids such as TG and cholesterol esters (ChE) were particularly depleted upon lipid deprivation in TIFM (Fig. 3L). These effects were ameliorated by arginine addition (Fig. 3L) or culture in standard conditions (Fig. 3 – Figure supplement 3B). TGs and ChEs function as lipid storage molecules that reside in lipid droplets, indicating that TIFM-cultured PDAC cells rapidly deplete these stores when deprived of lipids, whereas arginine supplemented cells are able to maintain lipid stores during starvation. Analysis of changes in the fatty acyl content of the cellular lipidome indicated that MUFA and SFA containing species were particularly depleted relative to PUFA containing species in TIFM-cultured cells (Fig. 3M and Figure 3 – Figure supplement 3C). This finding is consistent with arginine restriction preventing SREBP1-driven synthesis of MUFA and SFA species that enable PDAC adaptation to exogenous lipid starvation Lastly, we asked if inhibition of SREBP1 signaling would prevent arginine restricted cells from being able to adapt to lipid starvation. To determine the impact of lipid starvation on cell growth kinetics, we computed the growth rate of cells with or without lipids and calculated growth rate inhibition values (GR)^54^ to account for the differential growth kinetics of cells in TIFM or TIFM with additional arginine. We found that mPDAC cells cultured in TIFM cannot grow without exogenous lipids (Fig. 3N). In contrast, mPDAC cells cultured in TIFM with arginine supplementation can grow in the absence of lipids (Fig. 3N). SREBP1 is required for arginine supplementation to enable mPDAC cells to grow without exogenous lipids, as CRISPR-mediated knockdown of SREBP1 prevents arginine supplemented mPDAC cells from growing upon lipid starvation (Fig. 3O,P). We additionally asked if expression of mature SREBP1 would rescue lipid auxotrophy caused by arginine deprivation. However, we found that we were unable to express mature SREBP1 in TIFM-cultured mPDAC cells, despite successful overexpression in RPMI-cultured mPDAC cells (Fig. 3 – Figure Supplement 4A). Intriguingly, this suggests that nutrient stress in the TME may select against SREBP1 expression. Altogether, this data suggests that arginine restriction alters the lipidome of PDAC, increasing PUFA and TG content, while preventing PDAC cells from adapting to changes in exogenous lipid availability.

### Arginine restriction suppresses SREBP1 levels by activation of GCN2

We next sought to understand how arginine restriction decreases SREBP1 protein levels in PDAC cells. Activation of the amino acid sensor GCN2 and the integrated stress response (ISR) by amino acid deprivation is known to regulate SREBP1 specifically at the protein level^55,56^ (Fig. 4A). Therefore, we hypothesized that GCN2 activation upon arginine deprivation could suppress SREBP1 and lipid synthesis in PDAC. To determine if GCN2 was regulated by TIFM, we assessed levels of the ISR transcription factor ATF4 in TIFM-cultured PDAC cells. TIFM-culture increased expression of ATF4 (Fig. 4B), which was dependent on GCN2 (Fig 4C). Furthermore, transcriptomic (Fig. 4D) and immunoblot analysis (Fig. 4E) of PDAC cells cultured in TIFM with or without supplemental arginine indicated that arginine deprivation activated the ISR in TIFM. Thus, arginine deprivation in TIFM activates the ISR via GCN2 and could suppress SREBP1 via this nutrient sensing pathways.

**Figure 4:**
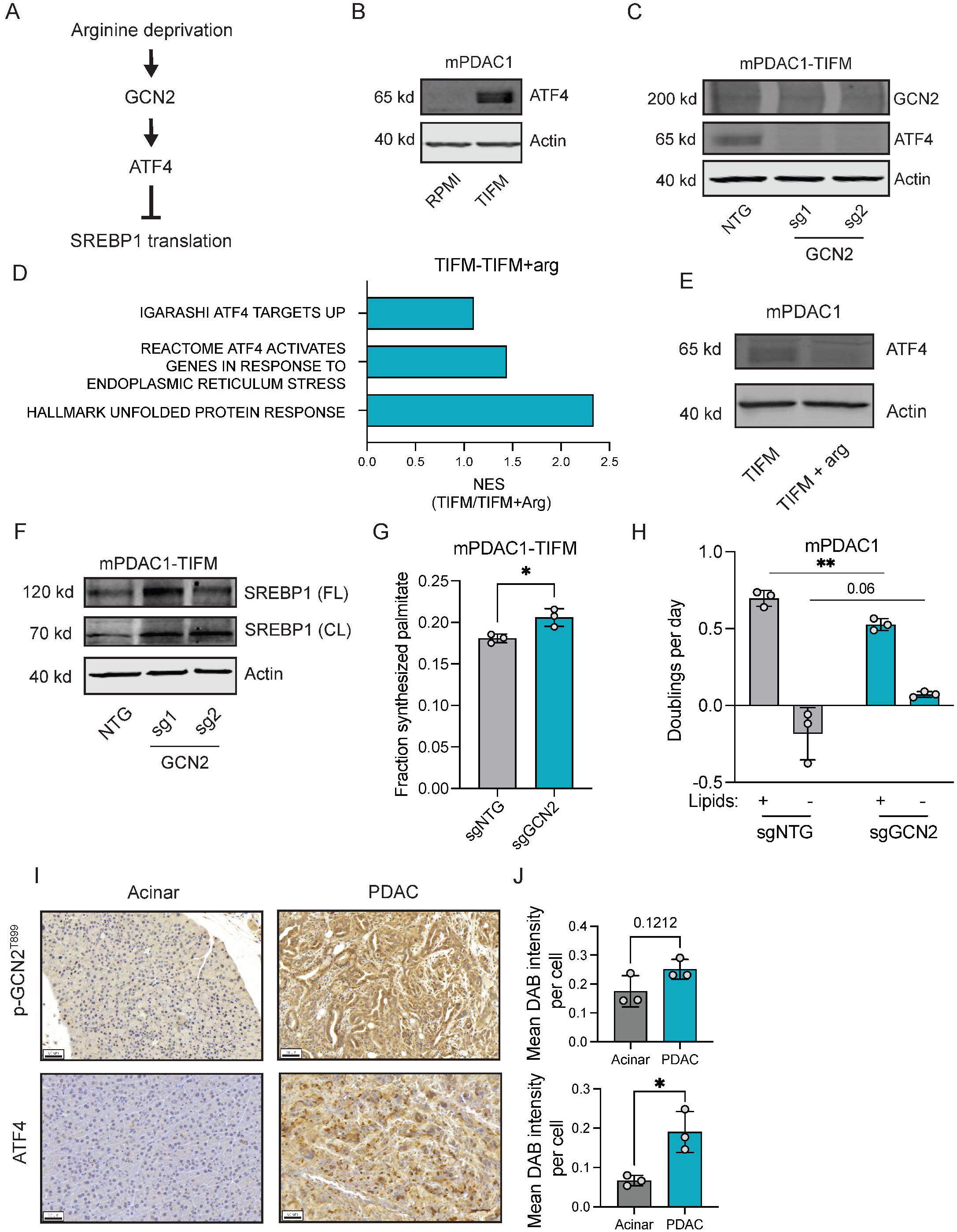
Arginine restriction suppresses SREBP1 levels by activation of GCN2. **(A)** Illustration of how GCN2 senses arginine and regulates SREBP1. **(B))** Immunoblot of ATF4 and GCN2 levels in mPDAC1-RPMI and mPDAC1-TIFM cells. **(C)** Immunoblot of ATF4 and GCN2 levels in mPDAC1-TIFM cells with or without CRISPR-mediated knockdown of GCN2. **(D)** GSEA NES scores of ISR related gene sets using transcriptomic data of mPDAC1 cells grown in TIFM or TIFM with supplemented arginine. **(E)** Immunoblot of ATF4 levels in mPDAC1 cells grown in TIFM or TIFM with supplemented arginine. **(F)** Immunoblot of SREBP1 levels in mPDAC1-TIFM cultures with GCN2 knockdown and non-targeting control. **(G)** The fraction of cellular palmitate (16:0) that is deuterium labeled in mPDAC1-TIFM cultures with CRISPR-mediated knockdown of GCN2 or non-targeting controls (n=3). **(H)** Growth rates of mPDAC1-TIFM cell lines with CRISPR-mediated knockdown of GCN2 or non-targeting controls in lipid depleted or lipid replete media (n=3). **(I,J)** Immunohistochemical analysis of ATF4 and activated GCN2 (p-T899) expression in *LSL-Kras^G12D/+^; TP53^-+^; Pdx-1-Cre* murine tumors (n=3) and adjacent acinar tissue from *LSL-Kras^G12D/+^; Pdx-1-Cre (n=2) or LSL-Kras^G12D/+^; TP53^-/+^; Pdx-1-Cre (n=1)*. Significance assessed in all panels using two-tailed Welch’s t-test.

We next asked if GCN2 activation in TIFM-cultured PDAC cells suppresses SREBP1 and lipid synthesis. GCN2 knockdown increased SREBP1 protein levels (Fig. 4F) and lipid synthesis (Fig. 4G). We further found that loss of GCN2 partially rescued the ability of these mPDAC-TIFM cultures to grow in the absence of exogenous lipids (Fig. 4H). These results indicate that arginine restriction suppresses SREBP1 and lipid synthesis in TIFM, in part by activation of GCN2. Lastly, we found that ATF4 levels are strongly increased and there is a trend towards increased levels of phosphorylated and active GCN2 in murine PDAC tumors compared with normal pancreas (Fig. 4I,J). Thus, this pathway is active and could contribute to suppression of SREBP1 and lipogenesis in vivo. Altogether, these data indicate that GCN2 activation, in part mediates suppression of SREBP1 and lipogenesis by arginine deprivation. However, as GCN2 deletion does not completely rescue SREBP1 levels in arginine starved cells, other mechanisms likely contribute.

### Arginine deprivation prevents PDAC cells from maintaining lipid homeostasis upon exposure to exogenous PUFAs

We next explored the implications of SREBP1 suppression in PDAC cells exposed to nutrient stress in the TME. PUFA-enriched cells are more sensitive to ferroptosis-inducing treatments because PUFAs have increased oxidative susceptibility in the cellular lipidome^26,31,52,61–64^. In addition to finding increased PUFA/MUFA in arginine-restricted cells (Fig. 3A,B), we also calculated the Cellular Peroxidation Index (CPI)^62^, a metric for the oxidative potential of the lipidome, of mPDAC cells cultured in TIFM with or without supplemented arginine. We found that TIFM cells without exogenous arginine had a significantly higher CPI than those with added arginine (Fig. 5A). Interestingly, as with PUFA/MUFA ratio (Fig. 3B), the increase in CPI in arginine restricted cells was not apparent in every lipid class. Specifically, TG, DG, SM and Cer species had increased CPI in arginine-restricted cultures, while PE, PC, ChE, LPC LPE and Hex1-Cer species did not have increased CPI (Fig. 5B). Thus, given the increase in relative PUFA content and CPI, we hypothesized that arginine-restricted PDAC cells would be more sensitive to induction of ferroptosis, a form of cell death involving lipid oxidation that cells enriched in polyunsaturated lipids are prone to undergo^65,66^. Surprisingly, arginine-restricted mPDAC cells did not consistently show decreased growth or viability when treated with xCT or GPX4 inhibitors that trigger ferroptosis (Fig. 5 – Figure supplement 1A-L). Thus, SREBP1-suppression by arginine restriction in the TME does not sensitize PDAC cells to ferroptosis inducers that target cysteine metabolism or GPX4.

**Figure 5:**
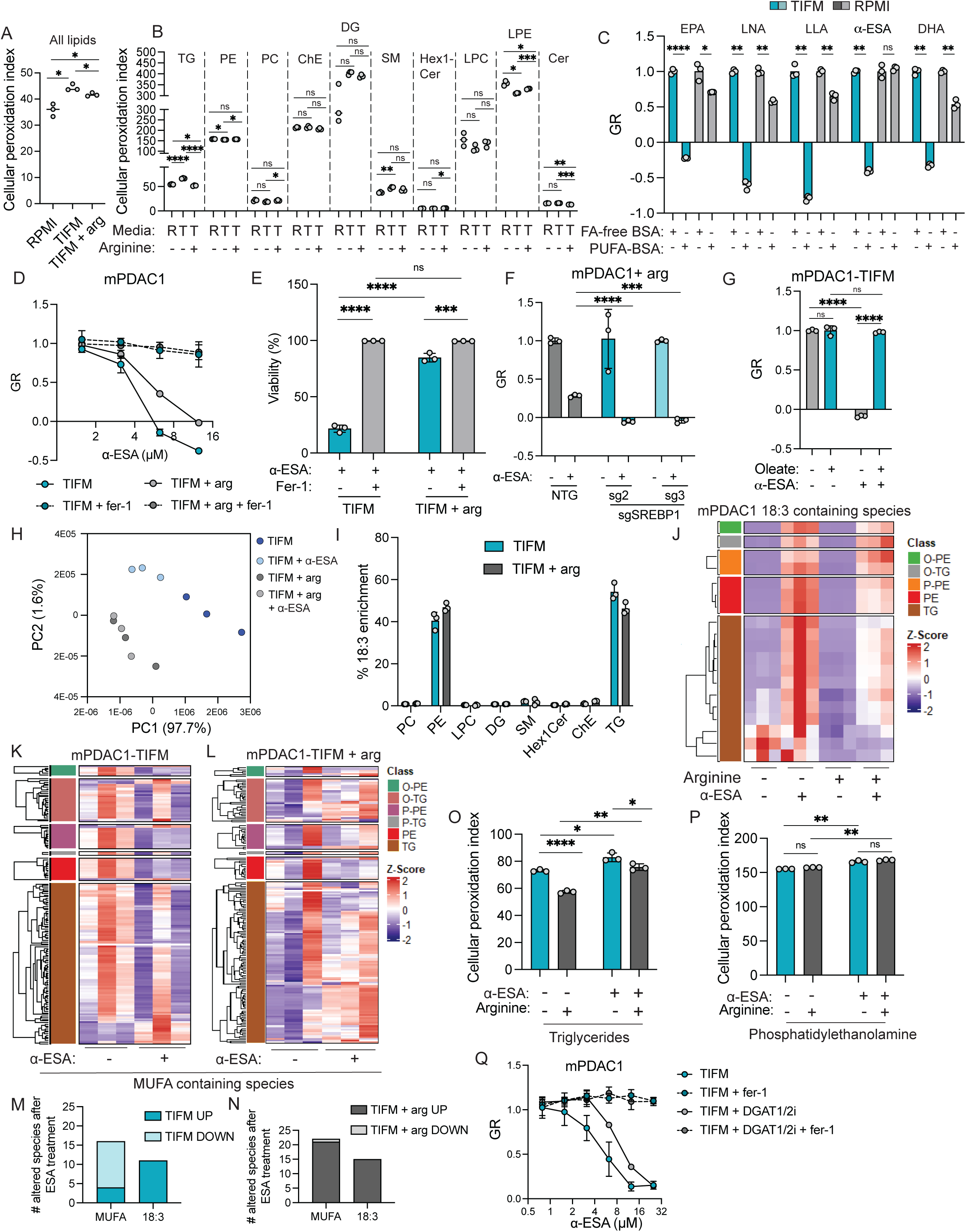
Arginine restriction sensitizes PDAC cells to exogenous PUFAs. **(A)** Cellular peroxidation index of all cellular lipids and **(B)** major classes of lipids species in mPDAC1 cells cultured in TIFM, RPMI or TIFM + arg (n=3). **(C)** Cellular GR value of mPDAC1-TIFM or mPDAC1-RPMI cells upon treatment with various PUFAs (n=3). (EPA: eicosapentanoic acid (250 µM); LNA: linolenic acid (250 µM); LLA: linoleic acid (250 µM); α-ESA: alpha-eleostearic acid (6.25 µM); DHA: docosahexaenoic acid (50 µM). **(D)** Cellular GR values of mPDAC1 cells cultured in TIFM or TIFM + arg treated with increasing doses of α-ESA with or without ferrostatin-1 (fer-1, 2 µM) (n=3 per condition). **(E)** Cell viability of mPDAC1 cells treated with 3.125 µM α-ESA after 48 hours of growth (n=3). **(F)** Cellular GR values of mPDAC1 cells cultured in TIFM + arg when transduced with control non-targeting or SREBP1 targeting sgRNAs and treated with α-ESA (6.25 µM) (n=3). **(G)** GR values of mPDAC1 cells cultured in TIFM when treated 6.25 µM α-ESA at indicated concentrations with or without supplementation of oleate (MUFA) at 100 µM (n=3). **(H)** Principal component analysis of LC-MS measurements of lipid levels of mPDAC1 cells cultured in TIFM or TIFM + arg with α-ESA (50 µM) or vehicle (fatty acid free BSA) (n=3). **(I)** Distribution of 18:3 fatty acyl groups in different complex lipid species. **(J)** Heatmap of TG and PE species with 18:3 acyl chains in mPDAC1 cells cultured as indicated upon treatment with α-ESA (50 µM). Plotted values are Z-scores for each lipid species (n=3). **(K,L)** Heatmaps of MUFA-containing PE and TG species in mPDAC1 cells cultured as indicated upon treatment with α-ESA (50 µM). Plotted values are Z-scores for each lipid species (n=3). **(M,N)** Analysis of the number of significantly altered lipid species containing MUFA or 18:3 acyl groups in mPDAC1 cells cultured as indicated upon treatment with α-ESA (50 µM). Significance cutoff p<0.05. **(O,P)** Calculated cellular phospholipid peroxidation index (CPI) for TGs **(O)** and PEs **(P)** in mPDAC1-TIFM and mPDAC1-TIFM + arg cells treated with α-ESA (50 µM) (n=3). **(Q)** Cellular GR values of mPDAC1 cells cultured in TIFM treated with indicated α-ESA concentrations and with or without 2.5 µM DGAT1 inhibitor (PF-04620110) and 2.5 µM DGAT1 inhibitor (PF-06424439) (n=3). Significance assessed in all panels using two-tailed Welch’s t-test.

Fatty acid synthesis maintains cellular lipid homeostasis by moderating the relative abundance PUFAs, MUFAs and SFAs in cellular lipids, primarily by producing MUFAs and SFAs^25,64^ and lowering PUFA uptake^25,67^. Imbalanced fatty acyl species in the cellular lipidome can result in impaired membrane function and cell death by apoptosis, due to SFA toxicity^62,63^, or ferroptosis, due to PUFA enrichment^65,66^. Because arginine restriction impairs fatty acid synthesis in PDAC cells, leading them to rely on uptake of exogenous fatty acids, we reasoned that arginine restriction would render PDAC cell sensitive to exogenous lipids that are enriched in one class of fatty acids, such as PUFAs^68^. To test this, we treated PDAC cell lines grown in TIFM with various PUFAs and measured their growth. PDAC cell lines grown in TIFM were potently sensitive to the addition of PUFAs, whereas cells grown in standard media were less affected by PUFA treatment (Fig. 5C). Thus, while arginine-restriction does not sensitize PDAC cells to cysteine/GPX4-targeting ferroptosis inducers, arginine-restriction does sensitize PDAC to exogenous PUFAs.

We next sought to understand why arginine-restricted PDAC cells are sensitive to exogenous PUFAs. We chose to focus our studies on the PUFA α-eleostearic acid (α-ESA) because TIFM cultured mPDAC cells are sensitive to this PUFA at low concentrations (Fig. 5C). Additionally, conjugated linolenic (18:3) acids like α-ESA are being investigated for their anti-neoplastic activity in culture and animal models of cancer^31,69–75^. We treated PDAC cells grown in TIFM or in TIFM with supplemental arginine with α-ESA. We found that TIFM-cultured PDAC cells were highly sensitive to α-ESA, as we observed previously (Fig. 5D,E and Fig. 5 Figure supplement 2A,B). Addition of arginine significantly decreased sensitivity to α-ESA (Fig. 5D,E and Fig. 5 Figure supplement 2A,B). Importantly, treatment with ferrostatin rescued growth α-ESA treated cells (Fig. 5D,E and Fig. 5 Figure supplement 2A,B), indicating that α-ESA treatment was suppressing the growth of arginine-restricted mPDAC cells by causing ferroptosis. Similarly, arginine restriction sensitized human PDAC cells to α-ESA induced ferroptosis (Fig. 5 – Figure supplement 2C). We next found that genetic (Fig. 5F) or pharmacological inhibition of SREBP1 (Fig. 5 – Figure supplement 2D) sensitized mPDAC cells to α-ESA in arginine replete conditions. We also found MUFA supplementation ameliorated α-ESA toxicity in arginine-restricted mPDAC cells (Fig. 5G). Altogether, these data indicate that arginine-restriction, by limiting SREBP1 activity, sensitizes PDAC to increases in exogenous PUFAs.

We hypothesized that arginine-restricted cells are sensitive to exogenous PUFAs because impaired SREBP1 activity and fatty acid synthesis could lead to PUFA overloading, or an increase in PUFA content that is incompatible with cellular viability. To test this, we performed lipidomic analysis on α-ESA-treated PDAC cells with or without supplemental arginine. In principal component analysis, α-ESA treated cells cluster separately from untreated samples in TIFM cultured mPDAC cells (Fig. 5H). In contrast, mPDAC cells cultured in TIFM + arg (Fig. 5H) or cultured in RPMI (Fig. 5 – Figure supplement 3A) do not cluster separately in this analysis. This indicates that α-ESA treatment has a larger impact on the cellular lipidome in arginine-restricted PDAC cells compared to PDAC cells not starved of arginine. We next investigated into which classes of lipids α-ESA was incorporated in mPDAC cells. We found that α-ESA treatment primarily increased levels of 18:3 fatty acid containing TG and PE lipids (Fig. 5I). This prompted us to analyze how arginine availability affected the impact of α-ESA supplementation on these lipid species. We found that arginine supplementation significantly reduced the incorporation of 18:3 fatty acids into PEs and TGs in TIFM cultured mPDAC cells (Fig. 5J). Culture in RPMI similarly led to reduced 18:3 fatty acids in PE and TG species (Fig. 5 – Figure supplement 3B). Along with the increase of 18:3 containing PE and TG, α-ESA treatment reduced levels of MUFA and SFA containing TG and PE species in TIFM cultured cells (Fig. 5K,M). This effect was also blunted by arginine supplementation (Fig. 5L,N) or culture in RPMI (Fig. 5 – Figure supplement 3C,D). Consistent with the finding that arginine supplementation blunts both the loss of MUFA/SFA and the incorporation of α-ESA into lipids, arginine supplemented cells had a lower TG CPI index (Fig. 5O). Notably, the CPI of PE species was largely unaffected by both arginine and α-ESA supplementation (Fig. 5P).

While polyunsaturated membrane phospholipids are thought to be primary drivers of ferroptotic cell death, recent reports indicate that neutral lipids can contribute to ferroptosis^31,76^. The above results indicate that α-ESA incorporation into TGs may contribute to ferroptotic cell death in arginine starved PDAC cells. To test this hypothesis, we inhibited diacylglyceride transferase 1 and 2 (DGAT1/2) to abrogate new synthesis of TGs and assessed the impact on α-ESA induction of ferroptosis in TIFM cultured cells. We found DGAT1/2 inhibition reduced the induction of ferroptosis in α-ESA treated cells (Fig. 5Q), which indicates that enhanced incorporation of α-ESA into neutral lipids contributes to ferroptosis induction by this PUFA in arginine-restricted PDAC cells. Altogether, these data show that arginine-restriction prevents cells from buffering their lipidome from exogenous PUFA supplementation, which results in PUFA overload in TGs that causes ferroptosis.

### Arginine-restricted PDAC cells and tumors are sensitive to PUFA enriched oils

Many lipids in the circulation are derived from dietary sources, which makes it possible to manipulate systemic lipid availability through dietary interventions^77–79^. We sought to determine if arginine-restriction in the PDAC TME would sensitize PDAC tumors to increased systemic PUFA availability brought on by dietary changes. We used tung oil to test this which is rich in α-ESA (Fig. 6A) and has previously been tested in animals as an anti-neoplastic dietary supplement^77^. We found, as with purified α-ESA, TIFM-cultured mPDAC cells are more sensitive to ferroptosis when treated with tung oil than arginine-replete cultures (Fig. 6B). Tung oil treatment also increases 18:3 PUFA content and CPI of TGs in TIFM cultured cells, an effect which is blunted by arginine supplementation (Fig. 6 – Figure Supplement 1A-D). To determine if tung oil treatment raises PUFA levels of PDAC tumors, we established orthotopic allograft tumors in mice and orally administered tung oil or safflower oil as a control (Fig. 6C), which contains balanced contents of SFAs/MUFAs/PUFAs (Fig. 6A). To determine if tung oil administration altered circulating fatty acid levels when orally administered, we measured the fatty acid content of the plasma in safflower and tung oil treated mice. Importantly, the only fatty acid modulated in the plasma by tung oil administration was α-ESA, which was enriched in tung oil treated mice (Fig. 6D). Tung oil treatment in tumor-bearing mice led to a significant decrease in tumor mass, which correlated with plasma α-ESA levels (Fig. 6E,F). Notably, tung oil administration increased the PUFA/MUFA ratio of PDAC tumors but not of healthy tissues (Fig. 6G). Thus, tung oil administration can be used to modulate plasma fatty acid PUFA content, specifically increase PUFA/MUFA ratio of tumors, and suppress the growth of established PDAC tumors.

**Figure 6:**
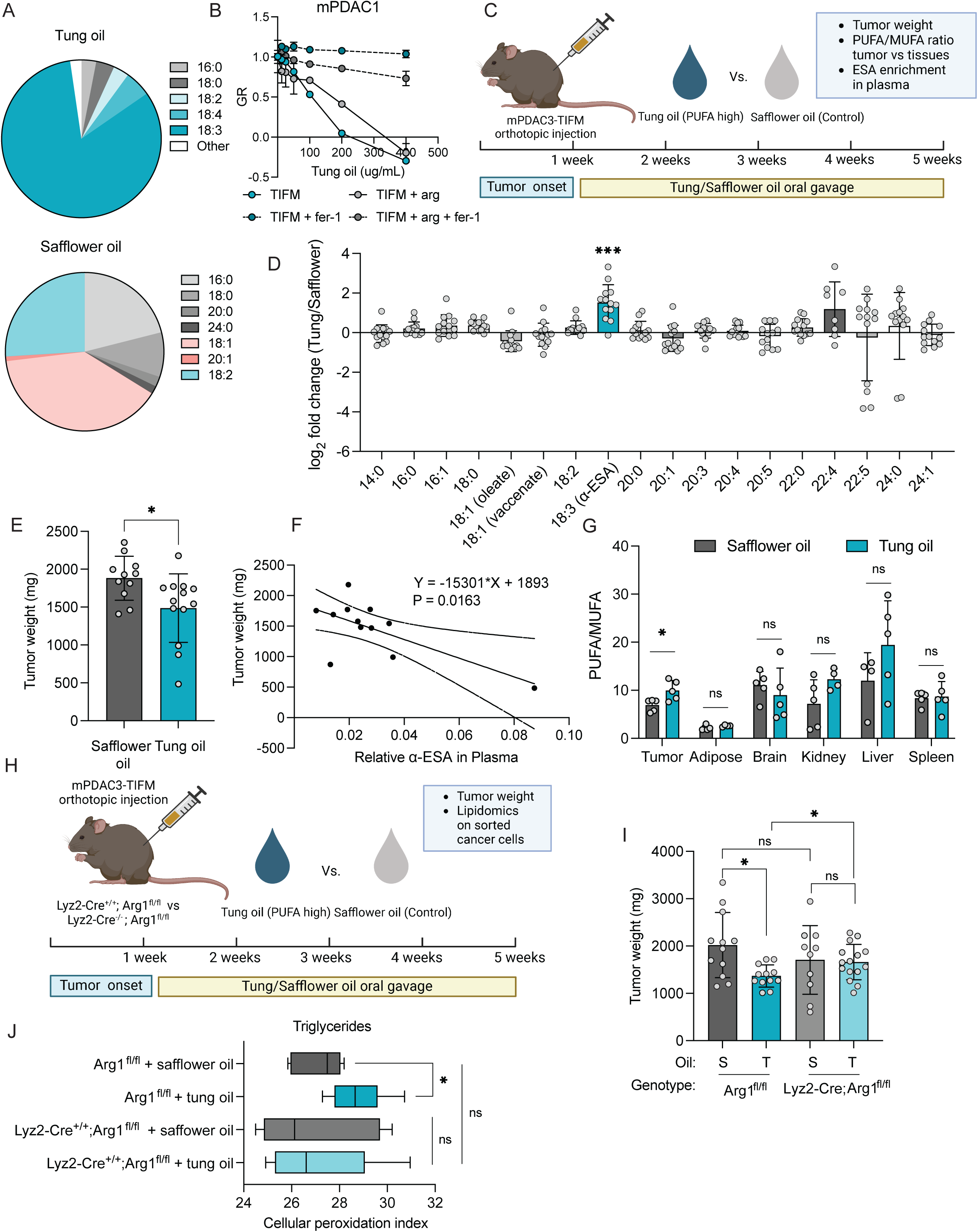
Arginine deprivation sensitizes PDAC cells and tumors to PUFA enriched oils. **(A)** Fatty acid content of tung and safflower oil measured by GC-MS fatty acid methyl ester (FAME) analysis (n=3 technical replicates). **(B)** GR values of mPDAC1 cells cultured in TIFM and TIFM + arg cell lines when treated with BSA-tung oil or fatty acid free BSA with or without ferrostatin-1 (fer-1, 2 µM) (n=3). **(C)** Diagram of workflow for treatment of mPDAC3-bearing mice with tung or safflower oil orally. **(D)** Log_2_ fold change of fatty acid methyl esters (FAMEs) in plasma of mice treated with tung or safflower oil (n=13 tung oil treated mice, n=12 safflower oil treated mice). **(E)** Tumor weights of mice treated with tung or safflower oil (n=11 safflower oil, n=13 tung oil). **(F)** Spearman’s correlation analysis of plasma α-ESA levels and PDAC tumor weights of mice treated with tung oil (n=13). **(G)** PUFA/MUFA ratios of PDAC tumors and other SREBP1 expressing (n=5 tung oil treated mice, n=5 safflower oil treated mice) **(H)** Diagram of treatment of mPDAC3-tumor bearing *Lyz2-Cre^+/+^; Arg1^fl/fl^* and *Lyz2-Cre ^-/-^; Arg1^fl/fl^* littermate control mice with tung or safflower oil. **(I)** mPDAC3 orthotopic tumor weights from *Lyz2-Cre^+/+^; Arg1^fl/fl^* and *Lyz2-Cr^-/-^; Arg1^fl/fl^* mice post tung or safflower oil treatment (*Arg1^fl/fl^*/Safflower n = 12; *Arg1^fl/fl^* /Tung n = 12; *Lyz2-Cre^+/+^; Arg1^fl/fl^*/Safflower n = 10; *Lyz2-Cre^+/+^; Arg1^fl/fl^*/Tung n = 15). **(J)** Cellular peroxidation index (CPI) for TGs in mPDAC3 cells sorted from tumor bearing *Lyz2-Cre^+/+^; Arg1^fl/fl^* and *Arg1^fl/fl^* mice given tung or safflower oil (*Arg1^fl/fl^* /Safflower n = 5; *Arg1^fl/fl^* /Tung n = 6; *Lyz2-Cre^+/+^; Arg1^fl/fl^* /Safflower n = 7; *Lyz2-Cre^+/+^; Arg1^fl/fl^* /Tung n = 6). Statistical significance in E and I was assessed using the Mann-Whitney U test. Statistical significance was assessed using a two-tailed Welch’s t-test correction in D, G, and J.

We next asked if arginine restriction in the TME and PUFA overloading contribute to suppression of PDAC growth by dietary PUFA supplementation with tung oil. We established orthotopic allograft tumors with YFP-labeled mPDAC cells in *Lyz2-Cre^+/+^; Arg1^fl/fl^* or control *Arg1^fl/fl^* animals that have high and low microenvironmental arginine levels, respectively (Fig. 2E). We then treated mice with tung or safflower oil (Fig. 6H). After treatment, tumor mass was measured and PDAC cells were sorted for lipidomic analysis. We observed that tung oil supplementation significantly impaired PDAC growth in control *Arg1^fl/fl^* host animals (Fig. 6I). In contrast, tung oil treatment did not significantly hinder the growth of arginine-replete tumors in *Lyz2-Cre^+/+^; Arg1^fl/fl^* animals (Fig. 6I). We asked if these differences in tumor growth could be driven by differences in PUFA levels between arginine-rich and arginine-starved PDAC tumors. Given our findings that α-ESA accumulates in TG species, we assessed how arginine levels in the TME affected PUFA levels in TGs, and in turn TG oxidation potential. We found that tung oil administration significantly increased the CPI of TGs in control *Arg1^fl/fl^* host animals (Fig. 6J). In contrast, tung oil treatment did not alter CPI of TGs in PDAC cells from arginine replete *Lyz2-Cre^+/+^; Arg1^fl/fl^* tumors (Fig. 6J). Collectively, these data show that arginine starvation in the PDAC TME sensitizes these tumors to dietary supplementation of PUFA-rich oils by increasing the amount of PUFA that accumulates in PDAC cells.

### Combining PUFA treatment with ferroptosis inducers may effectively target arginine-deprived PDAC tumors

Our findings indicate that arginine starvation increases sensitivity to PUFA-induced ferroptosis in PDAC cells, but not ferroptosis induction by inhibitors of cysteine uptake and GPX4 (Fig. 5 – Figure supplement 1A-L). Previous studies have found that Ras mutant tumors are sensitive to ferroptosis inducers that target these pathways^80^. PDAC tumors respond to targeting cysteine uptake^81,82^, although responses are incomplete^81,83–86^ and there is interest in identifying resistance mechanisms^30,86^ and methods to improve PDAC response to such treatments^83,84^. Therefore, we asked if PUFA supplementation could synergize with targeting GPX4 and cysteine uptake in arginine starved PDAC cells. To test this, we analyzed growth and viability of mPDAC cells in TIFM or RPMI treated with sublethal doses of α-ESA in conjunction with RSL3, to inhibit GPX4, and cystine withdrawal or erastin, to target cysteine uptake. Interestingly, we first noted that mPDAC cells in TIFM are more resistant to RSL3, erastin and cystine withdrawal (Fig. 7A-C). This suggests that nutrient availability in the TME may contribute to resistance to targeting ferroptosis via GPX4 inhibition or cysteine uptake in PDAC. We next found ferroptosis inducers killed cancer cells more effectively upon co-treatment with low doses of α-ESA that otherwise do not affect cell viability or proliferation (Fig. 7A-C). Critically, we found that sensitivity to ferroptosis inducers increased more in TIFM cultured cells compared to RPMI cultures, indicating that arginine restriction potentiates the effect of combining PUFA treatment with ferroptosis inducing drugs (Fig. 7A-C). We also examined if FSP1 inhibition would interact with α-ESA supplementation as FSP1 has recently been shown to reduce lipid peroxides in TGs in lipid droplets^76^. However, we observed no growth perturbations in mPDAC cell lines upon FSP1 inhibition (Fig. 7 – Figure Supplement 1). Together, these data suggest that PUFA supplementation can synergize with ferroptosis induction by targeting GPX4 and cysteine metabolism. Such combinations may prove useful in overcoming resistance to ferroptosis inducers in PDAC, which our data further suggest may be driven by TME nutrient availability.

**Figure 7:**
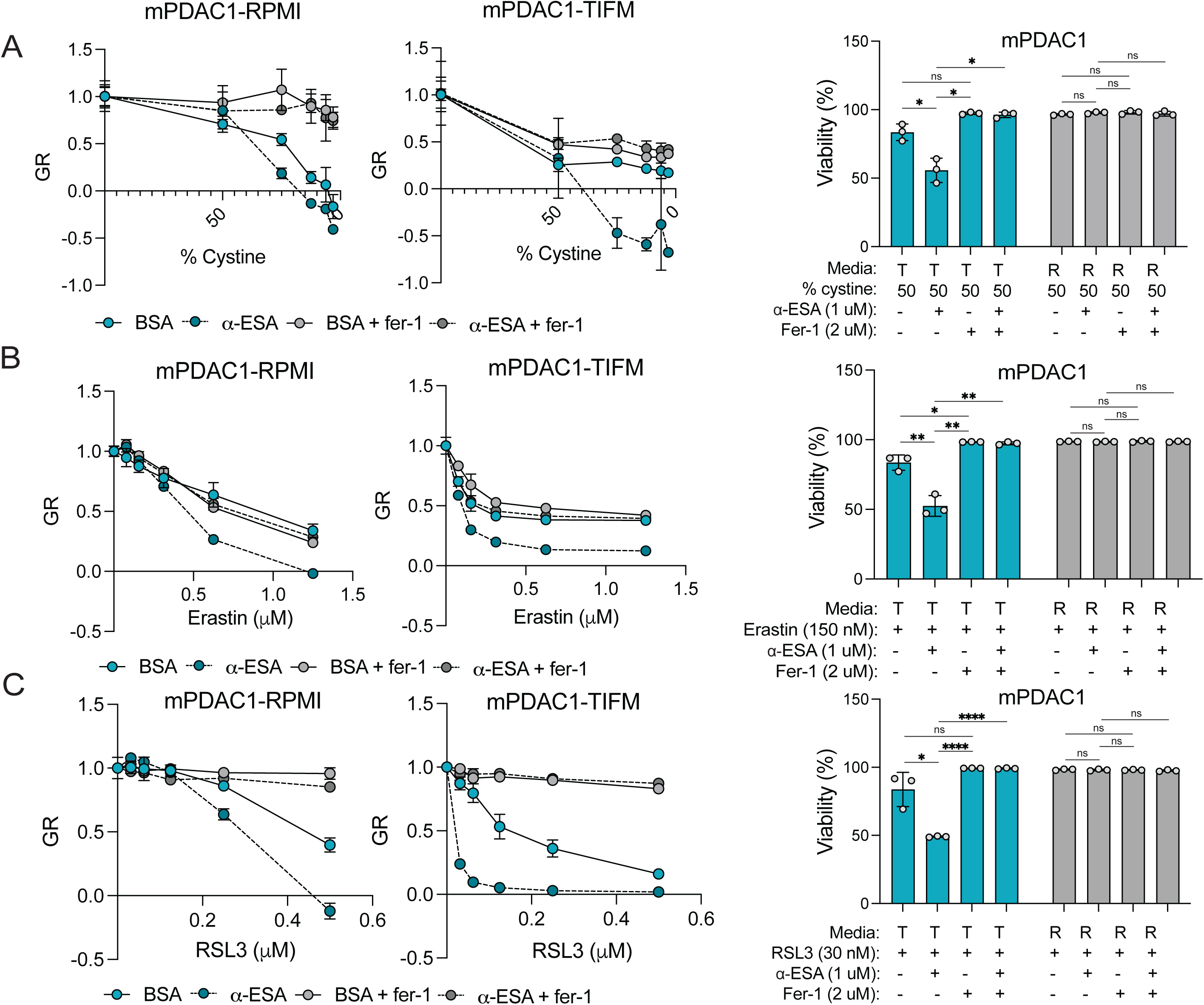
PUFA treatment synergizes with other ferroptosis inducers in arginine-restricted PDAC cells. GR value (left) and viability (right) of mPDAC1 cells cultured in RPMI or TIFM when treated with α-ESA (2 µM) of vehicle (fatty acid free BSA) in tandem with **(A)** deprivation of cystine to fraction present in base media as indicated, **(B)** Erastin at indicated concentrations, or **(C)** RSL3 at indicated concentrations. Ferrostatin-1 (fer-1, 2 µM) or vehicle (DMSO) was additionally added. Statistical significance was assessed using a two-tailed Welch’s t-test correction in all panels.

## Discussion

### Lipid metabolism in pancreatic cancer responds to arginine deprivation in the tumor microenvironment

Lipid metabolism in tumors is tightly linked to oncogenic mutations that cause cellular transformation^87–91^. Transformation activates lipid synthesis in cultured cells which promotes unrestricted cell growth^92^. As a result, targeting lipid synthesis in cancer cells has garnered attention as a potential cancer therapy^93,94^. However, cancer metabolism is not solely determined by cell-intrinsic features of cancer cells. Cancer cells also respond to cell-extrinsic stimuli in the tumor microenvironment^16,95–98^. Importantly, many microenvironmental features of tumors suppress cellular lipid synthesis^92,99^. How do these conflicting cues from cell-intrinsic oncogenic signals and the TME interact and influence the lipid metabolism of cancers?

Oncogenic Ras signaling leads to increased fatty acid synthesis in cultured cancer cells^100–102^. Despite activation of Ras in our PDAC models, transcriptomic analysis of PDAC cells in TIFM and in vivo shows that nutrient stress from the PDAC TME suppress SREBP1 signaling and fatty acid synthesis. These data suggest the cell-extrinsic cues are dominant over cell-intrinsic oncogenic signaling in dictating whether or not PDAC cells engage in fatty acid synthesis. This has several implications. First, cell culture studies that do not incorporate stimuli from the TME may overestimate the contribution of SREBP1 signaling and de novo synthesis to PDAC lipid metabolism and growth. Instead, given the constraints of the TME on *de novo* lipid synthesis, PDAC tumors likely require lipid uptake. Studies across tumor types support this, suggesting that uptake is the primary mechanism by which cancer cells acquire fatty acids^103^. Thus, future studies delineating the molecular mechanisms by which PDAC takes up lipids will be essential to understanding how lipid homeostasis is maintained in the metabolically constrained TME, given the lack of fatty acid synthesis in cancer cells.

The TME has many features that suppress de novo lipid synthesis in cancer cells^92,99^. We found that in vitro culture in TIFM, which models nutrient stress in the TME, recapitulates the suppression of SREBP1 signaling and lipid synthesis seen in vivo. This demonstrated that nutrient levels in the TME are important regulators of lipid metabolism. It also provided an opportunity to systematically identify specific metabolic cues in the TME that regulate SREBP1. Using TIFM, we showed specifically that low arginine levels found in PDAC tumors can suppress SREBP1 signaling and lipid synthesis. Alleviating arginine restriction in the TME increased SREBP1 levels and lipid synthesis, suggesting arginine starvation is a key constraint on lipid metabolism in PDAC tumors. Thus, in addition to known suppressors of lipid synthesis in the TME like hypoxia, amino acid stress also constrains lipid synthesis in PDAC. Arginine restriction regulates lipid metabolism in PDAC cells via downregulation of SREBP1 protein levels. SREBP1 is known to be regulated at the protein level by the arginine responsive nutrient sensor GCN2^55,56^. In this study, we identified GCN2 activation upon arginine starvation as partly responsible for repressing of SREBP1 expression and fatty acid synthesis in PDAC. Thus, we conclude that GCN2 plays a role in suppressing SREBP1 in arginine restricted PDAC cells. However, GCN2 deletion in TIFM cultured PDAC cells does not fully restore SREBP1 levels or fatty acid synthesis capacity to the same extent as arginine supplementation. Thus, while GCN2 activation is one key mechanism by which arginine starvation suppresses SREBP1-driven lipogenesis, future work will be required to fully elucidate how arginine restriction suppresses SREBP1.

### Stromal cells constrain the metabolism of pancreatic cancers

Solid tumors contain both malignant cells and a diverse population of stromal cells^105^. There is a growing appreciation that malignant and stromal cells interact metabolically in the TME^106–108^. Cancer-stromal metabolic interactions have largely been studied through the lens of how stromal cells facilitate the metabolism of cancer cells despite metabolic constraints in the TME. For example, lipid exchange from the stroma to PDAC cells contributes to PDAC metabolism and progression^109–112^. Thus, the metabolism of stromal cells and subsequent nutrient exchange with cancer cells can promote the ability of cancer cells to overcome the metabolic challenges of the TME. However, the interactions between cancer cells and the stroma do not necessarily increase the metabolic capacity of tumor cells. There is increasing evidence that some stromal-cancer metabolic interactions constrain the metabolic capabilities of cancer cells. For example, myeloid cells have been found to outcompete malignant cells for glucose in the TME^113^. We find that myeloid cells, by severely restricting arginine levels in the PDAC TME, suppress SREBP1 and lipid synthesis in PDAC tumors. Thus, stromal metabolic activity can impose metabolic limitations in the TME that constrain the metabolic capacity of tumors.

### Microenvironmental constraints on lipid synthesis render pancreatic cancers sensitive to polyunsaturated fatty acids: implications for therapeutically targeting ferroptosis

Precision dietary interventions targeting the metabolic liabilities of tumors are promising avenues in cancer therapy^114–117^. We have found that suppression of SREBP1 by arginine deprivation in the TME sensitizes PDAC cells to increased levels of exogenous PUFAs. Furthermore, arginine starvation sensitizes established murine PDAC tumors to dietary PUFA consumption. Interestingly, dietary PUFA consumption correlates with lower incidence of PDAC^118,119^ and recently dietary PUFA interventions were found to suppress PDAC initiation in animals^118^. Intriguingly, dietary PUFA supplementation also suppresses colorectal cancer (CRC) progression in animals^120,121^ and humans^122,123^. CRC tumors have recently been found to experience profound arginine depletion^124^. These findings suggest diets that manipulate PUFA levels could effectively suppress progression of tumors that are arginine starved, like PDAC and CRC. PUFA rich diets may also have an impact in tumors for which arginine deprivation therapy is being investigated clinically^125^. More broadly, our study suggests that the nutrient limitations of the TME, and the resulting constraints on cellular metabolism, are critical determinants of how diet intersects with tumor metabolism and progression.

PDAC tumors are prone to ferroptosis and can be targeted with ferroptosis-inducing therapies such as systemic cysteine depletion^81^. The underlying biochemistry that sensitizes PDAC to ferroptosis is incompletely understood. Oncogenic Ras signaling is known to increase the sensitivity of cells to ferroptotic cell death, especially cell death brought on by restriction of cysteine^81,126,127^. Thus, Ras signaling in PDAC likely contributes to ferroptotic sensitivity in these tumors. We have found that constraints on PDAC lipid synthesis arising from arginine starvation in the TME cause increased PUFA levels in PDAC cells. Further, arginine starvation sensitizes PDAC cells and tumors to PUFA overload and ferroptotic cell death upon PUFA supplementation. Thus, the TME, by restricting tumor lipid synthesis, also contributes to PDAC ferroptosis sensitivity. Our work suggests future studies are needed to understand how the TME regulates ferroptosis and response to therapies targeting this cell death mechanism^128,129^.

Lastly, in contrast to oncogenic Ras signaling, arginine starvation in the TME and suppression of lipid synthesis does not sensitize cancer cells to ferroptosis inducers that target proteins that quench lipid peroxides or uptake cysteine. Instead, arginine starved PDAC cells are sensitive to ferroptotic induction by PUFA supplementation. While future studies will be require to provide a mechanistic understanding of why arginine starvation only sensitizes PDAC cells to ferroptosis by PUFAs but not other ferroptosis inducers, this observation suggests that different methods of inducing ferroptosis can have different effects depending on the microenvironmental context. Importantly, as PDAC exhibits incomplete responses to ferroptosis induction by targeting cysteine^81,83–86^, our findings suggest that targeting ferroptosis via PUFA supplementation could leverage the microenvironmental context that renders these tumors especially sensitive to this form of ferroptosis induction. Furthermore, we show even a sublethal dose of PUFAs potently sensitizes PDAC cell lines to canonical ferroptosis inducers, particularly when they are exposed to levels of arginine present in the TME. This suggests combining PUFA-rich diets may increase the efficacy of other ferroptosis inducing therapies. Importantly, dietary PUFA supplementation specifically raises the PUFA content of PDAC, but not healthy tissues. Thus, future experiments to assess whether dietary PUFA supplementation increases the therapeutic window for targeting ferroptosis in tumors with limited lipid synthetic capacity are critical.

## Material and methods

### Cell lines and culture

Use of cancer cell lines was approved by the Institutional Biosafety Committee (IBC no. 1560). All cancer cell lines were tested quarterly for mycoplasma contamination using the Mycoalert Mycoplasma Detection Kit (Lonza, LT07-318). All cells were cultured in Heracell Vios 160i incubators (Thermo Fisher) at 37°C and 5% CO2. Cell lines were routinely maintained in RPMI-1640 or TIFM supplemented with 10% dialyzed FBS (Gibco, #26400-044, Lot#2244935P).

Cell culture was performed in static culture conditions. Because TIFM contains fewer nutrients than standard culture, media was changed on TIFM-cultured cell lines every 24 hours. Glucose was routinely measured using a GlucCell glucometer^130^ to ensure TIFM-cultured cells never experienced more than a 30% drop in glucose availability.

### Overexpression of mature SREBP1 in PDAC cells

Mature SREBP1 (amino acids 1-460) was cloned from pcDNA3.1-2xFlag-SREBP1a (a gift from Timothy Osborne) (Addgene plasmid #26801) and into pLV-EF1a-IRES-Blast (a gift from Tobias Meyer) (Addgene plasmid #85133) by Gibon assembly. pLV-EF1a-IRES-Blast was EcoRI digested and SREBP1(1-460) was PCR amplified Gibson assembly adapters in the primer sequences. These fragments were then assembled using HiFi Gibson assembly master mix (NEB E5510S). The sequence of the resulting vector was confirmed sequencing (Plasmidsaurus). HEK293T cells (Dharmacon) were transfected with the SREBP1(1-460) vector and the lentiviral packing plasmids psPAX2 (Addgene plasmid #12260) and pMD2.G (Addgene plasmid #12259). The medium was replaced after 24 hr, and lentivirus was harvested after 48 hr. Subconfluent mPDAC1 cells were infected with lentivirus using 8 μg/ml polybrene and infected cells were selected in 5 μg/ml blasticidin.

### CRISPR-mediated knockdown of SREBP1 and GCN2

sgRNAs targeting SREBP1 and GCN2 (Supplementary file 2) were taken from previously published optimized sgRNA designs for murine genes^131^. Oligonucleotide pairs were manufactured by Integrated DNA Technologies (IDT) and cloned into lentiCRISPRv2 (Addgene plasmid #52961) as previously described^132^. Lentivirus was produced as described above.

### Preparation of lipid stripped FBS

Lipid stripping of FBS was performed with a small modification from previously published methods^45^. Non-dialyzed FBS (Corning 35-010-CV) was split in two batches, one for chemical delipidation and one to use as a lipid replete control. For delipidation, di-isopropyl ether and n-butanol were mixed with FBS at a ratio of 0.8:0.2:1 for 30 minutes at room temperature. This mixture was then centrifuged at 4500xg for 15 minutes at 4C and the non-polar layer was removed. The aqueous layer was mixed 1:1 with di-isopropyl ether and mixed for 30 minutes at room temperature, then spun at 4500xg for 15 minutes. This process was repeated to ensure remove residual lipids remaining after the first delipidation. After lipid stripping, the aqueous layer was mixed under a gentle stream of nitrogen gas at room temperature for 2 hours to evaporate excess ether. This layer, as well as the non-delipidated fraction of FBS, was dialyzed against 9 g/L NaCl for 5 days at 4C using 10 kd SnakeSkin Dialysis Tubing (FisherScientific, PI88243). Dialysis buffer was changed every 24 hours. Before use, protein concentration was measured by BCA assay and protein concentration was normalized between lipid stripped and lipid replete FBS samples by addition of 9 g/L NaCl as needed.

### Measurement of cellular proliferation rates

Cellular proliferation rate was quantified using mPDAC cell lines expressing Nuclight-Red (Sartorius, 4717), a red fluorophore that localizes to the nucleus in live cells^133^. Nuclight-Red expressing cells were plated at a density of 1500 cells per well in clear bottom 96-well plates and allowed to attach overnight. Cells were scanned on the first and last day of treatment using a Sartorius Bioscience Incucyte S3. Live cells were identified by nuclear red fluorescence signal and counted. Cellular proliferation rate was calculated using the formular below:

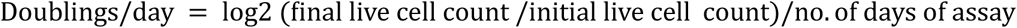

### Normalized growth rate inhibition analysis

GR values were determined from cells treated with various conditions as previously described^54^. In brief, GR value was determined using the following formula:

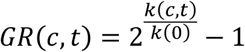

Where *k(c,t)* is the growth rate of cells at concentration = c and time = t, and *k(0)* is the average growth rate of cells in the control condition.

### Viability assays

PDAC cell lines were plated in 96-well tissue culture plates at a density of 1500 cells/well and allowed to attach overnight. The following day, the media was removed from each well and replaced with fresh media containing Cytotox Green dye (Sartorius, 4633) at 50nM and the indicated concentration of compounds or vehicle. Plates were scanned using a Sartorius/Essen BioScience IncuCyte S3 Instrument to monitor changes in cell viability and cell number over time. Unless otherwise indicated, all data shown reflects cell viability at 72 hours of the relevant treatment. Cellular segmentation and object quantification from the red and green fluorescence channels was performed using the IncuCyte analysis Cell viability was quantified based on the proportion of red and green objects identified across the entire well, using the following formula:

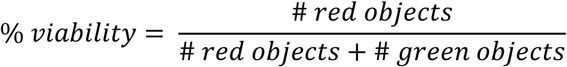

### Determining cell proliferation rate upon lipid starvation, fatty acid treatment, tung oil treatment, DGAT inhibition and ferroptosis inducing treatments

#### Lipid starvation

Cells were plated at 2000 cells per well in 12 well dishes in triplicate. After seeding, cells were washed once with 1x PBS and given indicated media supplemented with lipid replete FBS as a control or lipid stripped FBS. After 5 days, cellular growth rate was determined as described above.

#### Fatty acid treatment

Fatty acid free BSA (Sigma, A8066) was dissolved in deionized water to a final concentration of 0.17 mM solution. α-eleostearic acid (Cayman Chemical, 10008349), eicosapentaenoic acid (Cayman Chemical, 90110), linoleic acid (Cayman Chemical, 90150), linolenic acid (Cayman Chemical, 90210), and docosahexaenoic acid (Cayman Chemical, 90310) were dried under nitrogen gas. Fatty acid sufficient to yield 1 mM final solution was resuspended in the 0.17 mM fatty-acid free BSA solution. This solution was incubated on a rotator at 37 °C with intermittent sonication until the fatty acid fully dissolved. The FA-BSA solution was then filtered through a 0.22uM membrane and used immediately or stored at-20 °C. Tung oil lipids were conjugated to BSA by mixing tung oil (Sigma, 440337) with 4 mg/mL fatty acid free BSA in water 1:1 and rotating for 45 minutes. After rotating, tung oil-BSA conjugate was spun and the aqueous layer was removed and passed through a 0.22 um filter and used for analysis. 4 mg/mL fatty acid free BSA in water was prepared and used as a vehicle control.

To determine how fatty acids affected growth rates of PDAC cells, Nuclight-Red (Sartorius, 4717) expressing PDAC cells were plated at 1500 cells/well in clear-bottomed 96 well plate and allowed to adhere overnight. The following day, media was replaced with fresh media containing BSA:PUFA/tung oil or fatty acid free BSA vehicle control at the indicated concentrations. Media additionally contained, 2 µM Ferrostatin (MedChem Express, HY-100579) or DMSO (vehicle). Cellular proliferation rate was then measured as described above.

#### Ferroptosis induction

mPDAC cells were plated in 96 well dishes at 1500 cells per well. After seeding, cells were treated with indicated concentrations of RSL3 (Cayman Chemical, 19288), erastin (Cayman Chemical, 17754), or viFSP1 (Cayman Chemical, 39927), along with 2 µM α-ESA (Cayman Chemical, 22976) conjugated to fatty acid free BSA (Sigma, A8806) or an equivalent amount of fatty acid free BSA as vehicle control. For cystine deprivation experiments, RPMI and TIFM without cystine were formulated and exogenous cystine stocks (1000x, RPMI: 207 mM; TIFM: 50 mM) were made in sterile water and added to the cell culture media to provide the indicated range of cystine levels. 2 µM α-ESA or vehicle was additionally added to the cultures.

#### DGAT inhibition

mPDAC cells were plated in 96 well dishes at 1500 cells per well. After seeding, cells were treated with a range of α-ESA (Cayman Chemical, 22976) conjugated to fatty acid free BSA (Sigma, A8806) or equivalent volumes of fatty acid free BSA. PF-04620110 (Cayman Chemical, 16425, DGAT1 inhibitor) and PF-04624439 (Cayman Chemical, 17680, DGAT2 inhibitor) were then added at 2.5 uM or equivalent volume of DMSO as the vehicle control.

#### RNA-sequencing and analysis

Cells were plated at 50,000 cells per well in 6 cm dishes and grown in RPMI, TIFM, or TIFM + arg with or without exogenous lipids. After 24 hours, RNA was extracted from treated cells using the RNeasy Micro Kit (Qiagen, 74004), RNA quality and quantity were assessed using the 2100 Bioanalyzer System (Agilent).

#### Library preparation and sequencing

Strand-specific RNA-SEQ libraries were prepared using an TruSEQ mRNA RNA-SEQ library protocol (Illumina). Library quality and quantity were assessed using the Agilent bio-analyzer and libraries were sequenced using an Illumina NovaSEQ6000.

## Data analysis

Data processing and analysis were done using the R-based Galaxy platform (https://usegalaxy.org/)^134^. Quality control was performed prior and after concatenation of the raw data with the tools *MultiQC* and *FastQC*, respectively. All samples passed the quality check with most showing ∼20% sequence duplication, sequence alignment greater or equal to 80%, below and below 50% GC coverage, all of which is acceptable and/or indicative of good quality for RNASeq samples ^135,136^. Samples were aligned and counts were generated using the tools *HISAT2* (Galaxy Version 2.2.1+galaxy0, NCBI genome build GRCm38/mm10) and *featureCounts* (Galaxy Version 2.0.1+galaxy1), respectively. Differential expression analyses were performed with *limma* (Galaxy Version 3.48.0+galaxy1)^137^. Gene Set Enrichment Analysis (GSEA) was performed with *fgsea* (Galaxy Version 1.8.0+galaxy1)^138^ or GSEAPreranked (v6.0.12, https://gsea-msigdb.github.io/gseapreranked-gpmodule/v6/index.html; RRID: SCR_003199^139^. The ranking metric for all GSEA analyses was the t-statistic calculated in differential expression in *limma*. GSEA plots were generated as previously described^140^.

### Analysis of human SREBP1 scores

Analysis of human SREBF target gene expression was performed using the Metabolic gEne RApid Visualizer (MERAV) database^138^. Gene expression from the mSigDB geneset “Horton SREBF Target genes” were analyzed from PDAC cancer cell lines and PDAC patient tumors. Replicates from the downloaded data were averaged for both primary tumors and cell lines. SREBF scores were calculated by first median centering samples across each gene in both groups. log2 transformed values for each median centered gene in the dataset was then summed for each sample to obtain the SREBF gene score. Cell lines that were duplicated in the downloaded data were removed manually.

### Immunoblotting analysis

For immunoblotting analysis, cells were plated during the log-phase of growth into 6 cm dishes at a density of 500,000 cells per plate and allowed to grow overnight. Cells were then washed with PBS and then scraped off plates and sonicated 3 x 10 second intervals using a probe sonicator. Protein concentration of the lysate was determined by BCA assay (ThermoFisher Scientific, 23225). Protein lysates (20–50 μg) were resolved on sodium dodecyl sulfate–poly-acrylamide gel electrophoresis, 4–12% Bis-Tris Gels (Invitrogen, NP0321) and transferred to a polyvinylidene difluoride membrane using the iBlot 2 Dry Blotting System (Invitrogen, IB21001). Membrane was blocked with Intercept Blocking Buffer (LI-COR, 927-70001) at room temperature for one hour, stained with primary and secondary antibodies and then scanned using a LI-COR imager with Image Studio software version 2.1.10. Quantification of western blots was performed using the Image Studio software version 2.1.10. In brief, rectangles containing band signal were measured for total signal. For measuring the ratio of cleaved/full length SREBP1 expression, intensity of cleaved bands was divided by the intensity normalized of the full length band.

For immunoblotting of lipid-deprived cells, cells were plated during the log-phase of growth into 6 cm dishes at a density of 500,000 cells per plate and allowed to grow overnight. Cells were then treated with media with lipid stripped or lipid replete FBS. 24 hours later, cells were washed once with PBS and lysed in 200 µL RIPA buffer [25 mM Tris–Cl, 150 mM NaCl, 0.5% sodium deoxycholate, 1% Triton X-100, 1× complete protease inhibitor (Roche, 11836170001)]. To detect SREBP1 protein, cells were treated with 10 µM MG132 (Selleck Chemical, S2619) 6 hours prior to harvesting where indicated.

The following antibodies were used for immunoblot analysis: SREBP1 (Santa Cruz Biotechnology, sc-13551; 1:100 dilution), Beta-actin (Proteintech 660009-lg; 1:5000 dilution), p-S6 serine 235 (Cell Signaling Technology 2211S, 1:1000 dilution), ATF4 (Cell Signaling Technology 11815S, 1:1000 dilution), GCN2 (Cell Signaling Technology 3302S, 1:1000 dilution), TSC2 (Cell Signaling Technology 3612, 1:1000 dilution). The following secondary antibodies were used: IRDye 680LT Goat Anti-Mouse Ig (LI-COR, G926-68020; 1:10,000 dilution), IRDye 800CW Goat anti-Rabbit IgG (LI-COR, 926-32211; 1:10,000 dilution), and IRDye 800CW Goat anti-Mouse IgG (LI-COR, 926-32210; 1:10,000 dilution).

### Reverse-phase protein array (RPPA) analysis

Cells were plated in triplicate in 6-well dishes. To control or confluency 150,000 mPDAC-TIFM cells and 350,000 mPDAC-RPMI cells were plated per well. After seeding, cells were washed with PBS and then scraped off plates and sonicated 3 x 10 second intervals using a probe sonicator. Protein concentration of the lysate was determined by BCA assay (ThermoFisher Scientific, 23225).

Cell lysate samples were serially diluted two-fold for 5 dilutions (undiluted, 1:2, 1:4, 1:8; 1:16) and arrayed on nitrocellulose-coated slides to produce sample spots. Sample spots were then probed with antibodies by a tyramide-based signal amplification approach and visualized by DAB colorimetric reaction to produce stained slides. Stained slides were scanned on a Huron TissueScope scanner. Sample spots were identified and their densities quantified by Array-Pro Analyzer. Relative protein levels for each sample were determined by interpolating each dilution curve produced from the densities of the 5-dilution sample spots using a “standard curve” (SuperCurve) for each slide (antibody). Relative protein levels were designated as log2 values. Relative protein level data points were normalized for protein loading and transformed to linear values.

### Deuterium tracing of fatty acid synthesis

250,000 cells were plated in 2 mL culture medium in a 6-well plate with three technical replicates per condition. The following day, cells were washed with 1mL PBS and given 2 mL of media as indicated containing 50% D_2_O. Cells were allowed to grow for 6 hours media with 50% D_2_O. Lipids were then extracted by adding 300 µL 0.88% (w/v) KCl in LC/MS grade water and 400 µL LC/MS grade methanol containing butylated hydroxytoluene (BHT) and heptadecanoic acid to measure recovery. Cells were scraped and moved into a glass tube where dichloromethane was added and samples were vortexed at 4C for 15 minutes followed by centrifugation for 10 minutes. The organic fraction was separated and dried down with nitrogen gas. The insoluble and aqueous fractions were also dried and stored in-80 °C. These were later resolubilized in RIPA and protein amount was determined by BCA assay to normalize lipid measurements by protein content for each sample.

### FAME derivatization and gas chromatography-mass spectrometry (GC-MS) analysis

Dried lipid samples were resuspended in 100 µL toluene. 200 µL 2% sulfuric acid in methanol was then added to each sample. Samples were then incubated overnight at 50 °C. The next day, 500 µL 5% NaCl was added to each sample to quench the reaction. Samples were then extracted twice with 500 µL hexanes and the samples were dried under liquid nitrogen. Dried lipid samples were resuspended in 100 µL hexanes. Samples from animal tissues or biofluids were additionally run through Bond Elut LRC-Florisil columns (Agilent Technologies, 12113049) to remove contaminants. After loading lipids onto the LRC-Florisil column, FAMEs were eluted twice with 1 ml of hexane:diethyl ether (95:5 v/v), dried under nitrogen gas and resuspended in hexane for analysis.

All lipid samples were analyzed by GC-MS using a 8890 gas chromatograph system (Agilent Technologies) with a DB-FastFAME column (Agilent Technologies, G3903-63011) coupled with an 5997B Mass Selective Detector (MSD. Columns were pre-conditioned with 3 ml of hexane prior to analysis of FAME samples.Helium was used as the carrier gas at a constant pressure of 14 p.s.i. One microliter of sample was injected in splitless mode at 250 °C. After injection, the GC oven was held at 50 °C for 0.5 min, increased to 194 °C at 25 °C per min, held at 194 °C for 1 min, increased to 245 °C at 5 °C per min and held at 245 °C for 3 min. The MS system operated under electron impact ionization at 70 eV, and the MS source and quadrupole were held at 230 °C and 150 °C, respectively. The detector was used in scanning mode with an ion range of 104–412 *m*/*z.* Metabolites were analyzed using Masshunter software. Library files for FAME analysis were routinely updated using retention times from the Supelco 37 Component FAME Mix (Millipore Sigma, CRM47885). Mass isotopomer distributions were determined by integrating metabolite ion fragments and corrected for natural abundance using Isocor^141^. To calculate oil compositions, the intensity for the most abundant fragment for each FAME species were summed. The fraction of each of the summed intensity for each fatty acid species was then computed. FAMEs contributing less than 1% were not considered.

To calculate the PUFA/MUFA ratio from FAME data, peak areas of the most abundant fragment for each PUFAs detected by FAME (18:2, 18:3, 20:2, 20:3, 20:4, 20:5, 22:4, 22:5) were summed and divided by the peak areas of the most abundant fragment for each MUFA spcies detected by FAME (16:1, 17:1, 18:1, 20:1, 24:1).

### Animal experiments

Animal experiments were approved by the University of Chicago Institutional Animal Care and Use Committee (IACUC, Protocol #72587) and performed in strict accordance with the Guide for the Care and Use of Laboratory Animals of the National Institutes of Health. Mice were housed in a pathogen-free animal facility at the University of Chicago with a 12 hr light/12 hr dark cycle. Humidity is maintained between 30–70% humidity and temperature is maintained at 68–74 °F in the facility.

### Generation of orthotopic allograft PDAC tumor

C57BL6J mice 8–12 weeks of age were obtained from Jackson Laboratories (Strain #000664). 250,000 mPDAC cells were resuspended in 20 µl of 5.6 mg/ml Reduced Growth Factor Basement Membrane Extract (R&D Biosystems #3433-010-01) suspended in serum free RPMI. The BME:cellular mixture was injected into the splenic lobe of the pancreas to generate orthotopic tumors.

### Lyz2-Cre^+/-^; Arg1^fl/fl^ animal husbandry

C57BL6J *Lyz2-Cre^+/-^* and *Arg1^fl/fl^* mice were bred to generate *Lyz2-Cre^+/+^; Arg1^fl/fl^* and litter mate control *Arg1^fl/fl^*mice. All mice were genotyped using primers described previously^20–22^. Animal husbandry was carried out in strict accordance with the University of Chicago Animal Resource Center guidelines. Tumor implantation as described above was performed in mice at 8–20 weeks of age.

### Tumor interstitial fluid isolation from murine PDAC tumors

TIF from murine PDAC tumors was isolated as previously described^22^. Briefly, tumors from PDAC bearing mice were rapidly extracted after animals were euthanized by cervical dislocation. Tumors were then rinsed in blood bank saline (Thermo Scientific, 23-293-184) and blotted dry using Whatman paper. Tumors were then weighed, cut in half, and placed on a 20 µm filter (Spectrum labs, 145811) resting in a 50 mL conical tube. Tubes were loosely covered with 50 mL conical caps and tumors were centrifuged 400 x g for 10 minutes at 4 °C. The fluid that passed through the filter during centrifugation was then snap frozen and stored at-80 °C for future analysis.

### Tung oil administration to PDAC bearing animals

CBL57/6 mice 8-20 weeks in age were implanted with orthotopic allograft PDAC tumors as described above. After 1 week of tumor establishment, mice were orally gavaged once daily with 1.67 mL/kg tung oil or safflower oil for 4 weeks. After four weeks, mice were euthanized. Plasma, tumors and tissues were collected for analysis.

### Deuterium incorporation into fatty acids in PDAC bearing animals

Orthotopic tumors were implanted into 8-20 week old mice. After 3 weeks of tumor onset, mice were given 8% deuterated water (Cambridge isotopes, DLM-4-1L) which they consumed for 8 days. After 8 days, plasma was collected from mice and they were euthanized. Tumors were rapidly extracted from mice. Half of each tumor was clamped and snap frozen in liquid nitrogen and the other half was dissociated to a single cell slurry as previously described^22^. The PDAC cells were then sorted from the single cell slurry. To sort the PDAC cells, the single cell slurry was first blocked using 1% BSA in PBS. Cells were then stained for the PDAC cell marker mesothelin (MBL, D233-3, 1:100 dilution), washed, and stained with goat anti-Rat IgG APC secondary antibody (ThermoFisher, A10540, 1:200 dilution). PDAC cells were then sorted on a FACSAria Fusion 5-18 flow cytometer. A minimum of 50,000 PDAC cells were sorted per sample as we found samples with fewer cells did not provide sufficient material for GC-MS analysis. Tumor samples for which fewer cells were isolated were excluded from analysis. Cells were extracted and derivatized for FAME analysis as described above.

### Arginine extraction and analysis from interstitial fluid and plasma samples by Liquid chromatography-mass spectrometry (LC-MS)

5 µL of TIF or plasma sample was mixed with 45 µL of ice-cold acetonitrile containing stable isotope labeled amino acid mix from Cambridge Isotope Labs (MSK-A1-1.2) as internal standards. Samples were vortexed, incubated on ice for 20 minutes, centrifuged at 20,000 g for 20 minutes at 4°C. The supernatant was transferred to an autosampler vial for LC-MS analysis. Arginine calibration curves was prepared from 0.1µM - 500µM. Metabolite separation was performed on XBridge BEH amide column (2.1×150mm, 1.7 mm, Waters Corporation, MA). Mobile phase A was 90/5/5 water/acetonitrile/methanol, 20mM ammonium acetate, 0.2% acetic acid and mobile phase B was 90/10 acetonitrile/water, 10mM ammonium acetate, 0.2% acetic acid. The column temperature was 40 °C and flow rate was 0.3 mL/min. The chromatographic gradient was: 0min: 95% B, 9min: 70% B, 9.75min: 40% B, 12min: 40% B, 13min: 30% B, 14min: 30%B, 14.1min: 10% B,17min: 10% B, 17.5min: 95% B, 22min: 95% B. IQX orbitrap high resolution MS parameters were: sheath gas flow = 40, aux gas flow = 7, sweep gas flow = 1, spray voltage = 2800 for negative, 3600 for positive, ion transfer tube = 250 °C, vaporizer temp=350 °C, REF lens=60%. Data acquisition was done using Xcalibur 4.1 (ThermoFisher Scientific) and performed in switch polarity mode with a range of 70-1000 m/z, a resolving power of 60,000, an AGC target=100%, and a maximum injection time of 118ms. Tracefinder 4.1 was used for data analysis and quantification.

### LC-MS analysis of lipids

#### Analysis of lipids in PDAC cells starved of exogenous lipids

620,000 PDAC cells were plated in 15 cm dishes in indicated media and allowed to attach overnight. The following day, cells were treated with lipid depleted or lipid replete media and allowed to grow for 24 hours. Lipids were then extracted from cells as follows. Cells were kept ice cold throughout extraction. Samples were washed once with a 4°C saline solution (150 mM NaCl). 800 µL ice cold methanol containing 25 mg/L beta-hydroxytoluene and 600 µL ice cold 0.88% KCl in water were then added to each dish. Cells were scraped into glass tubes. 10 µL Mouse Splash Lipidomix (Avanti Polar Lipids, 330710) and 1600 µL ice cold dichloromethane was added to each sample. Cells were then shaken at 4°C for 15 minutes and spun down at 4500xg. The dichloromethane layer was then removed and dried down under nitrogen for analysis. The insoluble protein layer was dried down and kept at-80°C for quantification of protein amount.

#### Analysis of lipids in PDAC cells treated with α-ESA or tung oil

620,000 PDAC cells were plated in 15 cm dishes in indicated media and allowed to attach overnight. The following day, cells were treated with BSA-conjugated α-ESA (50 µM), or BSA-conjugated tung oil (200 µg/mL) or fatty acid free BSA. Lipids were extracted as described above. In addition to collection of protein for normalization of resulting LC-MS data, cell number for each plate were measured in triplicate using a ViCell (ThermoFisher Scientific) cell counting instrument. The average cell count for each sample was used to normalize LC-MS data.

#### Analysis of lipids in PDAC cells isolated from orthotopic tumor bearing mice fed either tung or safflower oil

YFP+ mPDAC cells were orthotopically implanted into C57BL/6 *Lyz2-Cre^-/-^; Arg^fl/fl^* or *Lyz2-Cre^+/+^; Arg1^fl/fl^* mice. After treatment with tung or safflower oil, tumors were removed and dissociated to a single cell slurry as previously described^139^. Cells were then passed through 70 µM filter and YFP positive cancer cells were sorted on a FACSAria Fusion 5-18 flow cytometer. We collected at least 500,000 cells per sample as we previously found this number of cells is needed for reliable LC-MS analysis. Samples with fewer sorted cells were excluded from analysis. Sorted cells were immediately pelleted and frozen in liquid nitrogen and kept at-80 °C prior to LC-MS analysis.

#### LC-MS data acquisition

All samples, blanks, and QC sample pools were resuspended in 50 µL of LC-MS grade acetonitrile (A955, Fisher) and LC/MS grade isopropanol (A461, Fisher) 50:50 (v/v), pulse vortexed, and sonicated for 5 minutes in a room temperature water bath prior to LC-MS/MS analysis. Samples were analyzed with a Vanquish dual pump liquid chromatography system coupled to an Orbitrap ID-X (Thermo Fisher Scientific) using a H-ESI source in both positive and negative mode. All samples were injected at 2 μL and lipids were separated with a reversed-phase chromatography using an Accucore C30 column (2.6 μm, 2.1mm × 150mm, ThermoFisher Scientific, 27826-152130) combined with an Accucore guard cartridge (2.6 μm, 2.1 mm × 10 mm, ThermoFisher Scientific, 27826-012105). Mobile phase A consisted of 60% LC-MS grade acetonitrile (ThermoFisher Scientific, A955). Mobile phase B consisted of 90% LC-MS grade isopropanol (ThermoFisher Scientific, A461) and 8% LC-MS grade acetonitrile. Both mobile phases contained 10mM ammonium formate (Sigma, 70221) and 0.1% LC-MS grade formic acid (ThermoFisher Scientific, A11710X1-AMP). Column temperature was kept at 50 °C, flow rate was held at 0.4 mL/min, and the chromatography gradient was as follows: 0-1 min held at 25% B,1-3 min from 25% B to 40% B, 3-19 min from 40% B to 75% B, 19-20.5 min from 75% B to 90% B, 20.5-28 min from 90% B to 95% B, 28-28.1 min from 95% B to 100% B, and 28.1-30 min held at 100% B. A 30 minute re-equilibration went as follows: 0-10 min held at 100% B and 0.2 mL/min, 10-15 min from 100% B to 50% B and held at 0.2 mL/min, 15-20 min held at 50% B and 0.2 mL/min, 20-25 min from 50% B to 25% B and held at 0.2 mL/min, 25-26 min held at 25% B and ramped from 0.2 mL/min to 0.4 mL/min, and 26-30 min held at 25% B and 0.4 mL/min. Each sample was injected twice, once for ESI-positive and once for ESI-negative mode. Mass spectrometer parameters were: source voltage 3250V for positive mode and-2800V for negative mode, sheath gas 40, aux gas 10, sweep gas 1, ion transfer tube temperature 300°C, and vaporizer temperature 275°C. Full scan data was collected using the orbitrap with scan range of 200-1700 m/z at a resolution of 240,000 for profiling samples. Primary fragmentation (MS2) was induced in the orbitrap with assisted HCD collision energies at 15, 30, 45, 75, 110%, CID collision energy was fixed at 35%, and resolution was at 15,000. Secondary fragmentation (MS3) was induced in the ion trap with rapid scan rate and CID collision energy fixed at 35% for 3 scans. Lipid identifications were assigned from ddMS3 files using LipidSearch (v5.0, Thermo), which was then used to generate a transition list for peak picking and integration in Skyline (v23.1).

## Data analysis

Raw peak areas for lipid species were normalized to total LC-MS signal intensity for analysis of lipid starved cells in Fig. 3. Raw peak areas for lipid species were normalized by the average cell number multiplied by cell volume for analysis of tung and α-ESA treated samples (Fig. 5,6). Lipids with a cumulative variance (%CV) greater than 30% in pooled QC samples were excluded from analysis. Principal component analysis was performed on mean-centered normalized lipid species peak areas using Metaboanalyst 6.0^142^. Volcano plots were generated by performing a Welch’s t-test on normalized peak areas and converting the resulting p-value into –Log_10_(p) value. Log_2_ fold change (Log_2_FC) was calculated by calculating the log_2_(treatment condition/control condition). Species not present in all samples were excluded from fold change analysis. The percent distribution of 18:3 containing lipids was calculated by summing the peaks of all 18:3 containing complex lipids for each species and dividing that by the total signal of 18:3 containing lipids.

### PUFA/MUFA ratio analysis

To calculate PUFA/MUFA ratios from LC-MS samples, only lipids with full identity were considered (all acyl chains identified in each lipid). Next, the quantity of monounsaturated and polyunsaturated acyl chains was calculated for each phospholipid. That number was then multiplied by the normalized peak area of the phospholipid to generate a monounsaturated or polyunsaturated score for each phospholipid. These values were then summed for all acyl chains for each lipid class (e.g. TGs, PEs etc) or for all lipid species. The sum PUFA score was then divided by the sum MUFA score of each individual sample to generate PUFA/MUFA ratios. An example of the PUFA and MUFA score calculation for a lipid is shown below:

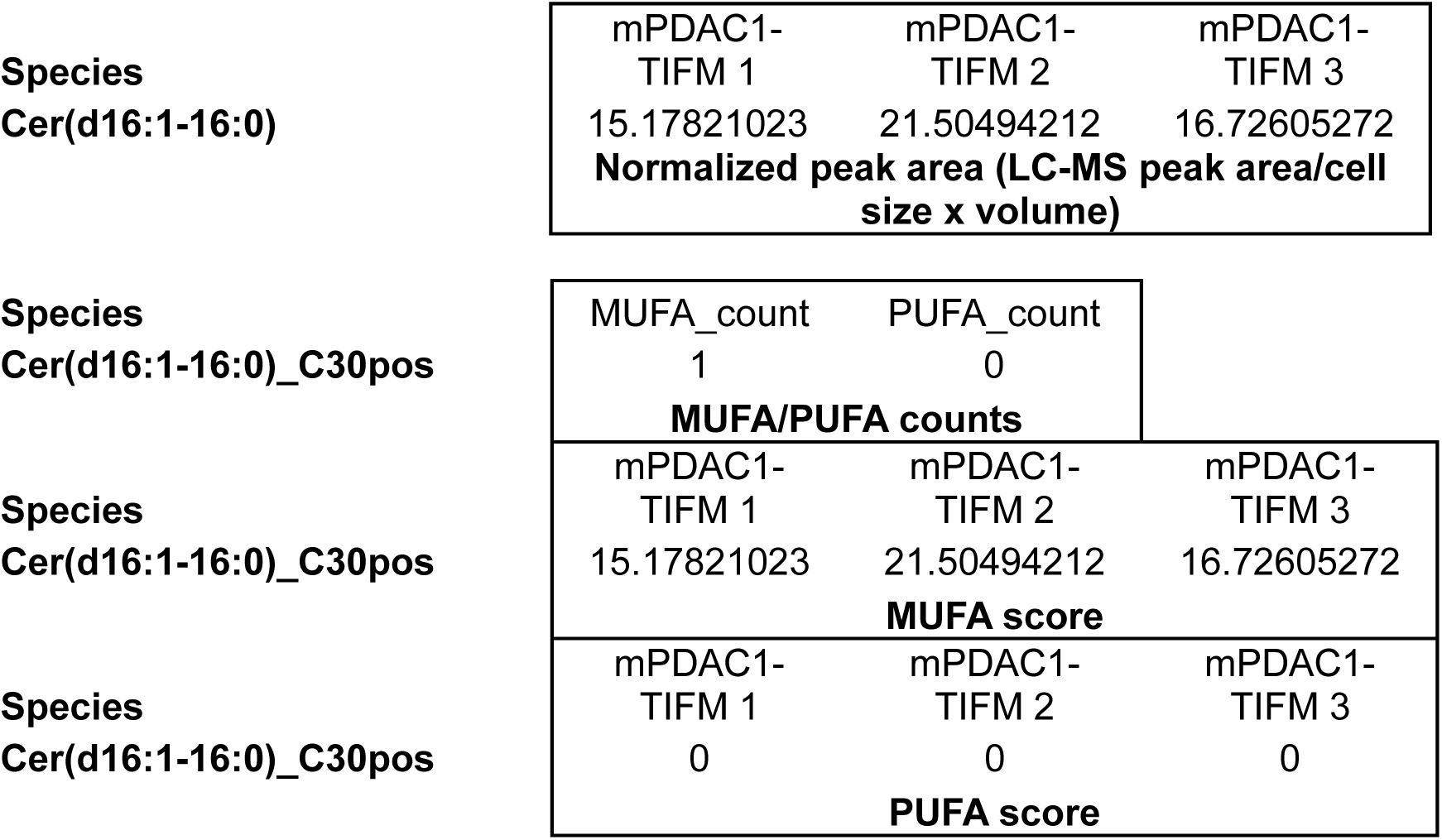

### Calculation of the Cellular Peroxidation Index (CPI)

Calculation of CPI was performed as previously described^62^ using only data from lipids for which we had complete identity. In brief, for each lipid class, or for all cellular lipids, we first calculated a score (as described above for calculating PUFA/MUFA ratios in lipids) for acyl chains containing 0-6 double bonds. We then summed these scores for fatty acyl chains with each double bond count to determine a total intensity score. We then determined the percentage of the total intensity score for acyl chains from 0-6 double bonds. The percentage of acyl chains with a given number of double bonds was calculated by taking the sum of acyl chain scores containing that number of double bonds in all lipid species divided by the sum of scores for all acyl chains in all lipid species. This percentage was then multiplied by its corresponding relative H atom transfer propagation rate constant^62^. These values were then summed for acyl chains with 0-6 double bonds to generate the CPI.

### Immunofluorescence analysis of SREBP1 in PDAC tumors

Whole tumors were placed in 15% sucrose dissolved in PBS at 4 °C for 6 hours. After 6 hours, tumors were moved into a solution of 30% sucrose dissolved PBS at 4 °C for 24 hours. Tumors were removed and embedded in histology cassettes with optimal cutting temperature (OCT) compound (Fisher 23-730-571). Tumors were cut into 5 µm thick sections and mounted onto glass microscopy slides. Slides were thawed to room temperature for 15 minutes in the dark, after which slides were washed once with PBS for 5 minutes. Cells were permeabilized for 15 minutes at room temperature in 0.1% Tween-20 dissolved in PBS (PBST). Slides were then blocked for 1 hour in 10% normal goat serum (Invitrogen, 50062Z). Slides were then washed once with PBST and SREBP1 antibody (Abcam ab28481) at 1:150 in 1:1 PBST with 10% goat serum as previously described^50,51^. Slides were mounted into a humidified chamber at 4 °C and allowed to stain in SREBP1 antibody overnight. The following day, slides were washed 3 times for 5 minutes with PBST in the dark. After this, Goat anti-Rabbit IgG (H+L) Cross-adsorbed secondary antibody (Invitrogen A21244) was added at 1:250 dilution in 1:1 PBST with 10% normal goat serum for 1 hour in the dark at room temperature. Slides were washed 3 times with PBST and mounted using mounting medium with DAPI (Invitrogen, P36962). Slides were scanned using the Olympus VS200 whole slide scanner using the YFP channel to analyze cancer cells, Cy5 channel for SREBP1, and DAPI channel to scan cell nuclei.

Analysis of SREBP1 in tumors was performed using quPath^143^. Individual cells were classified by DAPI using quPath cell detection. Cancer cells were identified based on YFP expression using the Object Classifier function. QuPath object classifier then used data from this training to identify other YFP expressing cells in each image. Images were spot checked for accuracy before analysis. For analysis of SREBP1 staining intensity, three sections were selected from each tumor slice based on DAPI intensity as a metric of cellularity. SREBP1 staining intensity (Cy5 channel) was calculated for YFP-expressing cancer cells in that section. Sample identity was blinded prior to selection of sections for analysis. An 361 YFP positive cells were selected from each tumor. 9 tumors were analyzed from both *Lyz2-Cre^-/-^; Arg^fl/fl^* and *Arg^fl/fl^*animals. The mean cell intensity of SREBP1 signal was calculated using quPath cell measurement function. The mean intensity for each cell was then plotted from *Lyz2-Cre^-/-^; Arg^fl/fl^* and *Arg^fl/fl^* animals.

### Immunohistochemical analysis of ATF4 and p-GCN2^T899^ in PDAC tumors

Immunohistochemistry on mouse tumor sections was performed as previously described^144^ using head denaturation in citrate buffer (pH 6.0). Stained slides were scanned using an Olympus VS200 Whole Slide Scanner. Antibodies and dilutions used for immunohistochemistry were the following: ATF4 (Cell Signaling Technology 11815, 1:50); p-GCN2^T899^ (Biorbyt orb6078, 1:200). Qupath was used to quantify mean DAB intensity per cell for each slide. Briefly, 3-8 representative regions were selected from each slide and traced in Qupath. Automated cell counting and DAB intensity quantification was accomplished using the ‘Cell detection’ and ‘Positive Cell detection’ tools with standard settings and a cell expansion of 15 um, which correctly identified the area of each cell. After detecting the mean cell intensity of each cell in a given slide, a random, equal number of cells (n=388 cells for p-GCN2, n=762 cells for ATF4) was selected for each slide and the mean DAB intensity per cell quantified.

### Immunofluorescence analysis of lipid droplets

YFP-expressing mPDAC cells were plated at 300 cells/well in Ibidi 8-well chamber slides (Ibidi, 80806) in RPMI or TIFM media. Cells were allowed to attach overnight. Cells were washed once with PBS and fixed with 4.0% formaldehyde for 15 minutes at room temperature. After fixation, cells were treated with 1:200 dilution of LipidTox neutral lipid stain (Invitrogen, H34477) in PBS for 30 minutes. Cells were washed once and then imaged on a Zeiss LSM 980 Airyscan 2 microscope at 40x magnification. Cells were identified using the YFP marker and lipid droplets were analyzed. For analysis, cells outlines were defined using the YFP signal and used to measure cellular areas. For lipid droplet analysis, background was subtracted from images and droplets were detected using auto thresholding by the Renyi Entropy method. Images were then smoothed, despeckled, and droplets were quantified using the Analyze Particles function in FIJI^145^ based on LipidTox positivity.

### qRT-PCR analysis for SREBP1 target genes in human PDAC cells

RNA was extracted from cells using the RNeasy Mini Kit and optional on-the-column DNAse digestion (QIAGEN, 74104). Extracted RNA was converted to cDNA by reverse transcription using the High-Capacity cDNA Reverse Transcription Kit (Applied Biosystems, 4374967). Expression levels of Elovl6 transcript were amplified using PowerUp SYBR Green Master Mix (Invitrogen, A25741) and primers targeting human Elovl6 from GeneCopeia (MQP037222). Quantification was performed using a QuantStudio 3 RealTime PCR System (Applied Biosystems). The average change in threshold cycle (ΔCt) values was determined for each of the samples relative to actin levels (Genecopeia, HQP016381) and compared with vehicle control (ΔΔCt). Finally, relative gene expression was calculated as (2−ΔΔCt). Experiments were performed in triplicate cultures.

## Data availability

Sequencing data from Fig. 1A-G was previously published^142^ and is deposited in GEO with the accession code GSE199163. Sequencing from Fig. 1 – Figure supplement 1B and Fig. 3 is deposited in GEO with the accession code GSE288530. Raw mass spectra data for LC-MS based measurements of arginine (Fig. 2) and lipids (Fig. 3-5) have been deposited in the NIH sponsored Metabolomics Workbench repository under project IDs ST003714, ST003715, ST003716, ST003717, and ST003718.

## Acknowledgements

We thank all members of the Muir lab, especially James Martin III and Lindsey Dzierozynski, for feedback and discussions on this project. We thank Costas Lyssiotis and Marina Pasca di Magliano at the University of Michigan for the generous gift of *Lyz2-Cre* and *Arg1^fl/fl^* mice. We thank Evan Lien and Ryan Sheldon at the Van Andel Institute, and Hardik Shah from the University of Chicago, for providing helpful insight into design, execution, and analysis of lipidomics data. We would also like to thank Jessalyn Ubellacker and Isaac Harris for providing feedback on this project. We thank the University of Chicago Animal Resources Center (RRID:SCR_021806), especially Ani Solanki, for their assistance with animal models of pancreatic cancer. We thank the University of Chicago Genomics Facility (RRID:SCR_019196), Human Tissue Resource Center (RRID:SCR_019199), Metabolomics Platform (RRID:SCR_022932), Cellular Screening Center (RRID:SCR_017914) and Organoid and Primary Culture Research Core (RRID:SCR_022935) and Integrated Light Microscopy Core (RRID: SCR_019197) for experimental assistance in running assays. We would also like to thank the Gardel lab and Steven Redford for training us and allowing us to use their Zeiss LSM 980 Airyscan 2 microscope. This work was supported by funding from the Phi Beta Psi Sorority, the Ludwig Center for Metastasis Research, the Cancer Research Foundation (2025 Team Science Award) and the University of Chicago Comprehensive Cancer Center (P30 CA014599) provided to A.M. P.B.J. was supported by the National Cancer Institute (T32 CA009594) and the University of Chicago Mary Ellen Connellan Award.

## Ethics

Animal experiments were approved by the University of Chicago Institutional Animal Care and Use Committee (IACUC, Protocol #72587) and performed in strict accordance with the Guide for the Care and Use of Laboratory Animals of the National Institutes of Health.

**Figure 1 – source data 1** Table of significantly differentially expressed metabolism gene sets between mPDAC cells cultured in TIFM and RPMI as shown in Fig. 1B. Gene set names, enrichment scores, normalized enrichment scores, p-values, adjusted p-values, gene set sizes, and leading edges for each geneset are shown in this table.

**Figure 1 – source data 2** Mass isotopomer distribution of palmitate analyzed by GC-MS from mPDAC cells after treatment with deuterated water as shown in Fig. 1L. Peak areas are reported in this table for palmitate methyl ester isotopomers M+0 through m+32.

**Figure 1 – source data 3** Mass isotopomer distribution of palmitate analyzed by GC-MS from mPDAC cells after treatment with deuterated water as shown in Fig. 1 - Figure supplement 1E. Peak areas are reported in this table for palmitate methyl ester isotopomers M+0 through m+32.

**Figure 1 – source data 4** Full unedited immunoblots (left) and unedited immunoblots with sample and band identification (right) for immunoblots shown in Fig. 1J and Fig. 1 – Figure supplement 1C.

**Fig. 1 – Figure supplement 1 – source data 1** Tables of linear and log transformed levels of proteins from RPPA analysis of mPDAC1-3-RPMI and mPDAC1-3-TIFM cultures.

**Figure 2 – source data 1** Mass isotopomer distribution of palmitate analyzed by GC-MS from mPDAC cells after treatment with deuterated water as shown in Fig. 2B. Peak areas are reported in this table for palmitate methyl ester isotopomers M+0 through m+32.

**Figure 2 – source data 2** Mass isotopomer distribution of palmitate analyzed by GC-MS from mPDAC3 cells sorted from tumors from *Lyz2-Cre^+/+^; Arg1^fl/fl^* or *Lyz2-Cre^-/-^; Arg1^fl/fl^* mice as shown in Fig. 2G. Peak areas are reported in this table for palmitate methyl ester isotopomers M+0 through m+32.

**Figure 2 – source data 3** Mass isotopomer distribution of palmitate analyzed by GC-MS from plasma of *Lyz2-Cre^+/+^; Arg1^fl/fl^* or *Lyz2-Cre^-/-^; Arg1^fl/fl^* mice bearing mPDAC3 tumors as shown in Fig. 2H. Peak areas are reported in this table for palmitate methyl ester isotopomers M+0 through m+32.

**Figure 2 – source data 4** Full unedited immunoblots (left) and unedited immunoblots with sample and band identification (right) for immunoblots shown in Fig. 2A,C.

**Figure 3 – source data 1** Table of peak areas from LC-MS analysis of mPDAC1 cancer cell lipids cultured in TIFM, RPMI, or TIFM + arg, with or without exogenous arginine supplementation. As shown in Figure 3C,I-M and Fig. 3 - Figure supplement 3. NA indicates the metabolite not detected.

**Figure 3 – source data 2** Mass isotopomer distribution of palmitate analyzed by GC-MS from mPDAC cells after treatment with deuterated water as shown in Fig. 3H. Peak areas are reported in this table for palmitate methyl ester isotopomers M+0 through m+32.

**Figure 3 – source data 3** Full unedited immunoblots (left) and unedited immunoblots with sample and band identification (right) for immunoblots shown in Fig. 3D,F,O.

**Figure 3 – source data 4** Mass isotopomer distribution of palmitate analyzed by GC-MS from mPDAC cells after treatment with deuterated water as shown in Fig 3 – Figure supplement 3C-F. Peak areas are reported in this table for palmitate methyl ester isotopomers M+0 through m+32.

**Figure 3 – source data 5**Full unedited immunoblots (left) and unedited immunoblots with sample and band identification (right) for immunoblots shown Fig. 3 – Figure supplement 2A,B.

**Figure 3 – source data 6** Full unedited immunoblots (left) and unedited immunoblots with sample and band identification (right) for immunoblots shown in Fig. 3 – Figure supplement 4.

**Figure 4 – source data 1** Full unedited immunoblots (left) and unedited immunoblots with sample and band identification (right) for immunoblots shown in Fig. 4B.

**Figure 4 – source data 2** Full unedited immunoblots (left) and unedited immunoblots with sample and band identification (right) for immunoblots shown in Fig. 4C.

**Figure 4 – source data 3** Full unedited immunoblots (left) and unedited immunoblots with sample and band identification (right) for immunoblots shown in Fig. 4E.

**Figure 4 – source data 4** Full unedited immunoblots (left) and unedited immunoblots with sample and band identification (right) for immunoblots shown in Fig. 4F.

**Figure 4 – source data 5** Mass isotopomer distribution of palmitate analyzed by GC-MS from mPDAC cells after treatment with deuterated water as shown in Fig. 4G. Peak areas are reported in this table for palmitate methyl ester isotopomers M+0 through m+32.

**Figure 5 – source data 1** Table of lipid peak areas from LC-MS analysis of mPDAC1 cancer cells cultured in TIFM, RPMI, or TIFM + arg with or without exposure to α-ESA as shown in Fig. 5A,B,H-P and Fig. 5 – Figure supplement 3. NA indicates the metabolite was not detected.

**Figure 6 – source data 1** Table of lipid peak areas from LC-MS analysis of mPDAC1 cancer cells cultured in TIFM, RPMI, or TIFM + arg with or without exposure to tung oil-BSA conjugates as shown in Fig. 6 – Figure supplement 1A-D. NA indicates the metabolite was not detected.

**Figure 6 – source data 2** Table of peak areas of GC-MS fatty acid methyl ester (FAME) analysis of plasma of mice treated with tung oil or safflower oil as shown in Fig. 6D.

**Figure 6 – source data 3** Table of peak areas of GC-MS fatty acid methyl ester (FAME) analysis of tumors and tissues of mice treated with tung oil or safflower oil as shown in Fig. 6G.

**Figure 6 – source data 4** Table of peak areas from LC-MS analysis of mPDAC3 cells sorted from orthotopic tumors from *Lyz2-Cre^+/+^; Arg1^fl/fl^* or *Lyz2-Cre^-/-^; Arg1^fl/fl^* mice treated with tung or safflower oil as shown in Fig. 6J.

**Supplementary file 1** Table with complete formulation of Tumor Interstitial Fluid Medium (TIFM).

**Supplementary file 2** Table of sgRNA sequences used in this study.

---

**Figure 1 – figure supplement 1:**
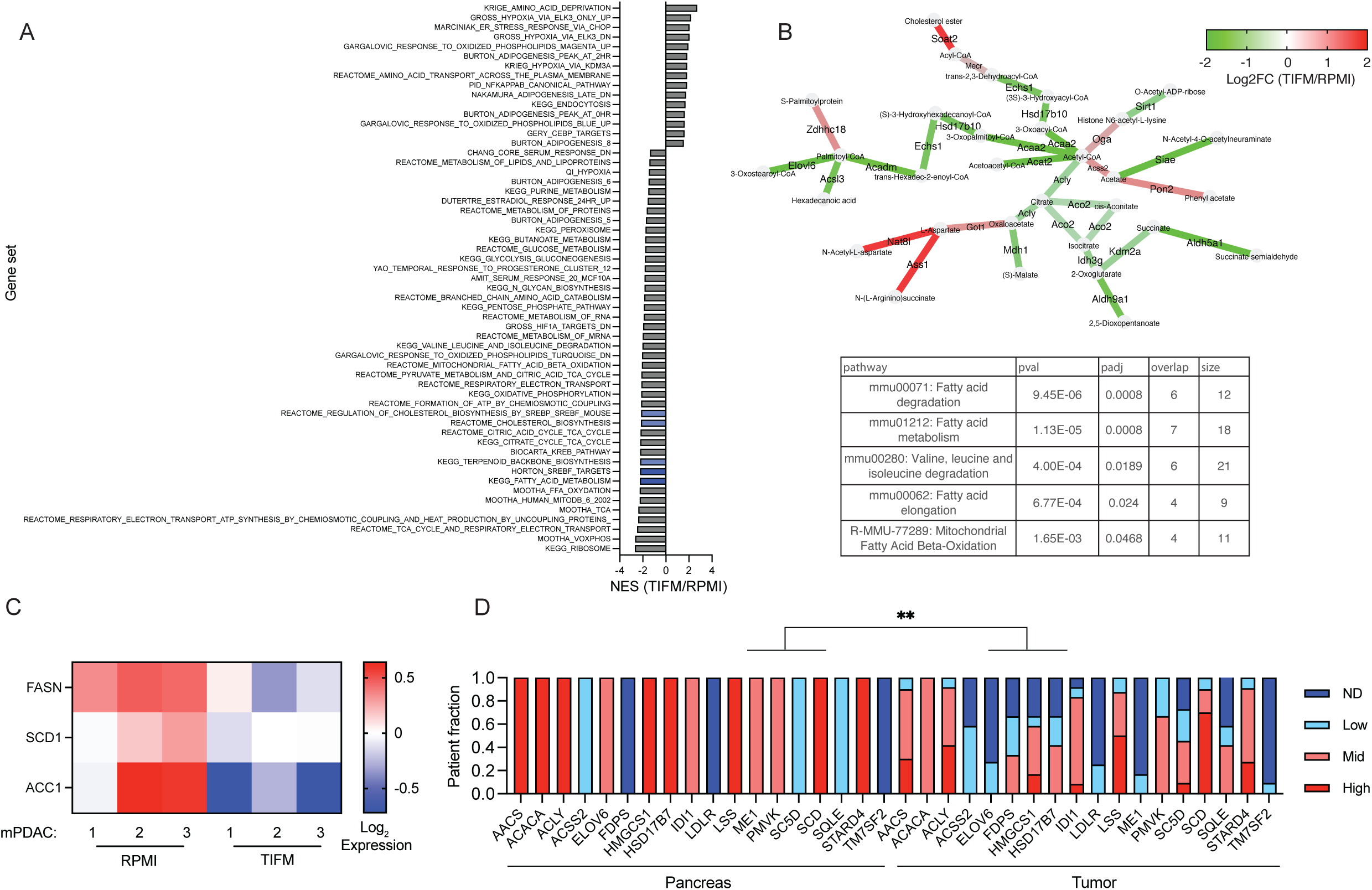
Transcriptomic analysis reveals exogenous nutrient availability regulates expression of SREBP-regulated fatty acid synthesis genes. **(A)** Complete list of significantly altered metabolism gene sets in mPDAC cells grown in TIFM and in RPMI. **(B)** GATOM analysis of transcriptomic data from mPDAC cells cultured in TIFM or RPMI. **(C)** Heatmap of log_2_ transformed protein levels of SREBP1 regulated lipogenic genes in mPDAC1-3-RPMI and mPDAC1-3-TIFM cells from RPPA analysis. **(D)** Proportion human PDAC and normal pancreas specimens from the Human Protein Atlas^146^ classified with low, mid, high, or non-detectable (ND) expression of SREBP1 target gene expression. Significance was tested in D using Fisher’s exact test.

**Figure 1 – figure supplement 2:**
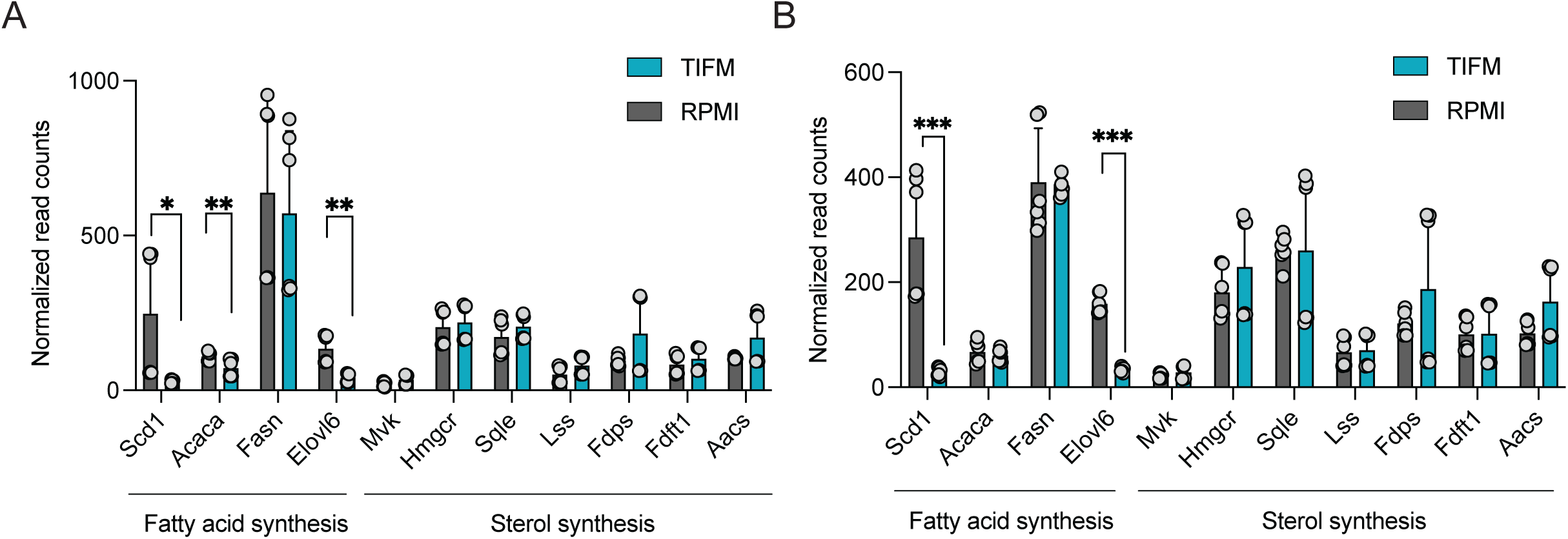
Transcriptomic analysis reveals exogenous nutrient availability specifically regulates SREBP1. (A,B) Read counts of fatty acid (SREBP1) or sterol (SREBP2) synthesis genes in TIFM and RPMI cultured mPDAC cells (two experiments plotted, n=3 each, n=6 total). Significance was determined using a two-tailed Welch’s t-test.

**Figure 1 – figure supplement 3:**
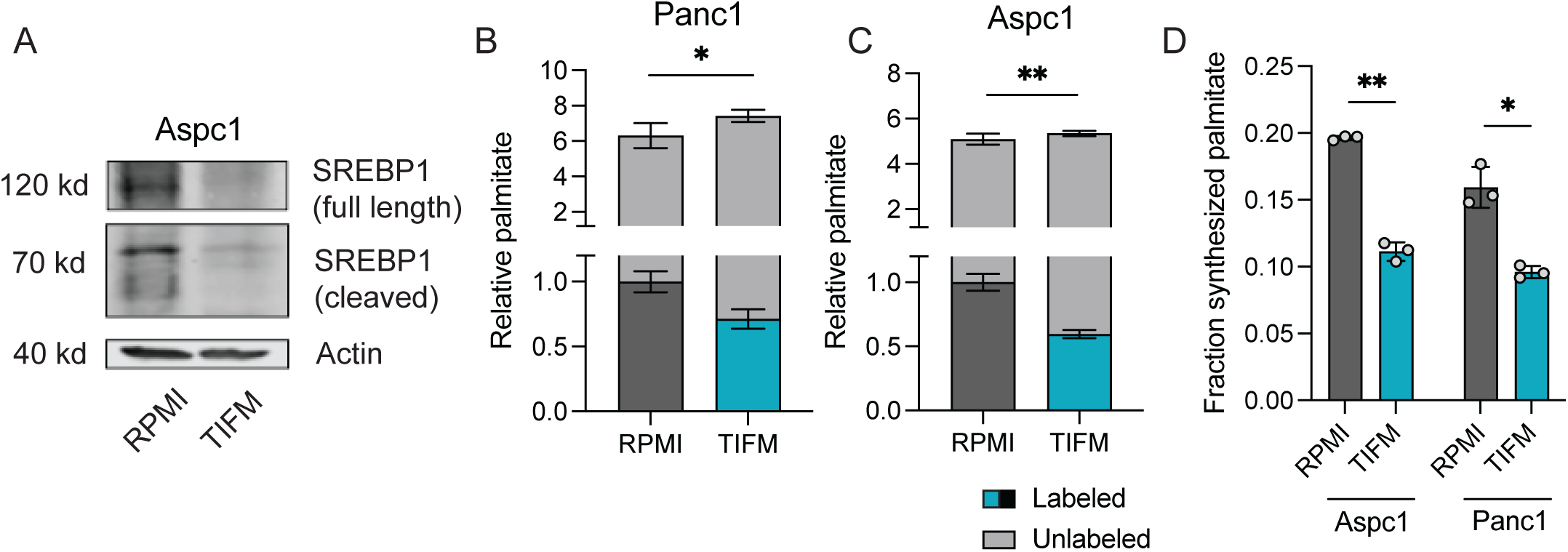
TME nutrients downregulated SREBP1 and fatty acid synthesis in human PDAC. **(A)** Immunoblot analysis of full length and cleaved SREBP1 expression in Aspc1 human cell lines grown in TIFM or RPMI. **(B,C)** Levels of unlabeled (grey) and deuterium labeled (blue, dark grey) palmitate (16:0) in human PDAC cell lines grown in TIFM or RPMI. Labeled palmitate indicates de novo synthesized lipid (n=3). **(D)** The fraction of cellular palmitate (16:0) that is deuterium labeled indicated human PDAC cell lines in TIFM or RPMI (n=3). Significance was determined using a two-tailed Welch’s t-test for B-D.

**Figure 2 – figure supplement 1:**
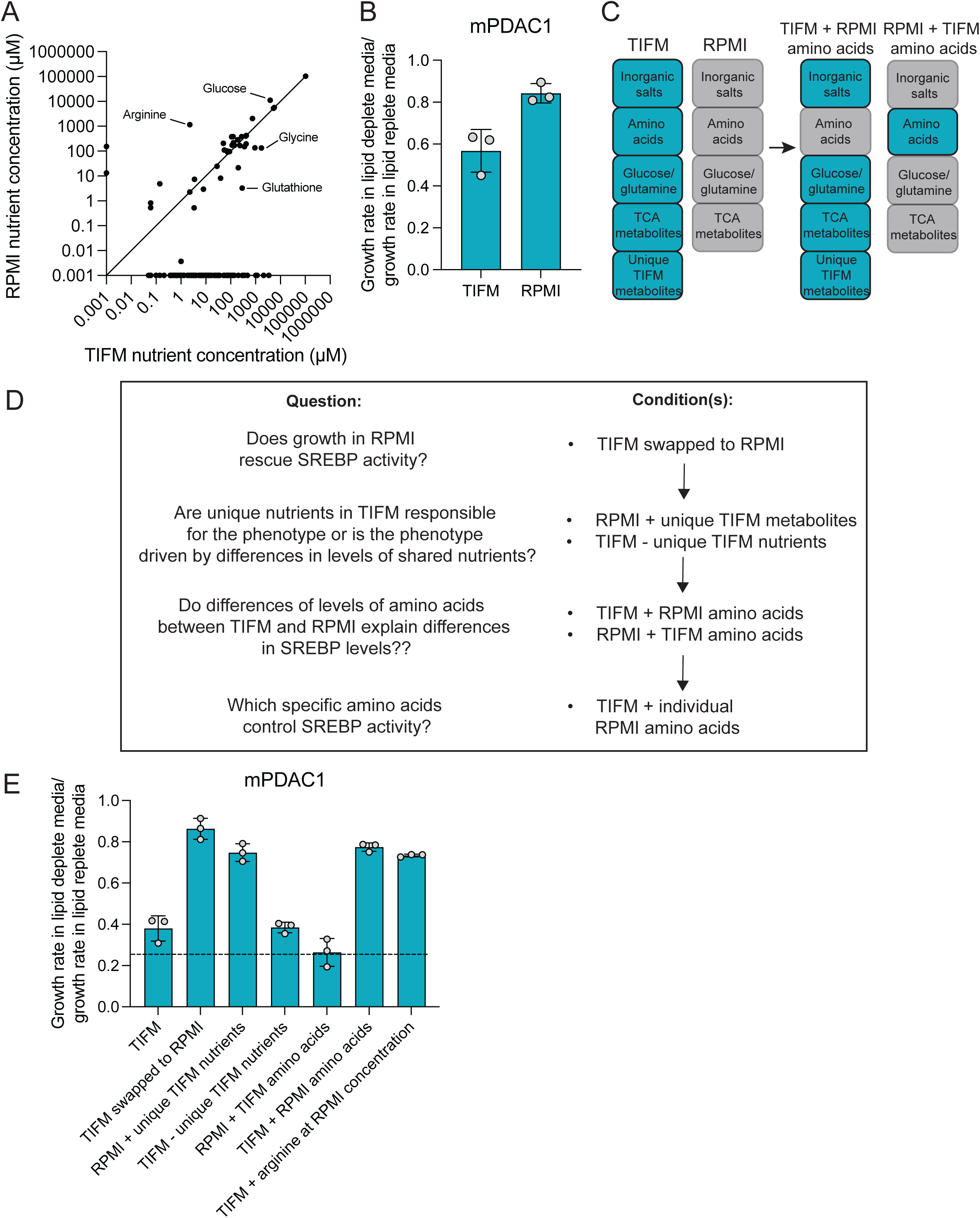
Systematic analysis of TIFM content identifies arginine restriction in TIFM as a regulator of PDAC lipid metabolism. **(A)** Concentrations of nutrients in TIFM or in RPMI. Each point represents an individual nutrient in RPMI and in TIFM. **(B)** Cell proliferation of mPDAC1 cell lines TIFM or RPMI with or without exogenous lipids (n=3). **(C)** Diagram of nutrient swapping strategy in TIFM. Nutrient pools (e.g. amino acids) are swapped between TIFM and RPMI to determine the nutrients that drive cellular phenotypes such as growth without exogenous lipids. **(D)** Logical flow chart and **(E)** data from nutrient swap experiments to identify TIFM nutrients that affect the ability of mPDAC1 cells to grow in the absence of exogenous lipids. Briefly, chemicals unique to TIFM when added to RPMI did not prevent mPDAC1 cells from growing in lipid depleted conditions, nor did removing these chemicals from TIFM improve mPDAC1 growth without lipids. This led us to conclude that alterations of levels of a shared nutrient between TIFM and RPMI was responsible for differences in lipid metabolism. We then observed that mPDAC1 cells grown in TIFM containing RPMI amino acid levels were able to grow without lipids, indicating that differences in amino acid availability altered lipid metabolism. We then found that supplementation of arginine in TIFM to RPMI levels alone was sufficient to improve mPDAC1 growth without lipids to levels nearly seen in RPMI (n=3).

**Figure 2 – figure supplement 2:**
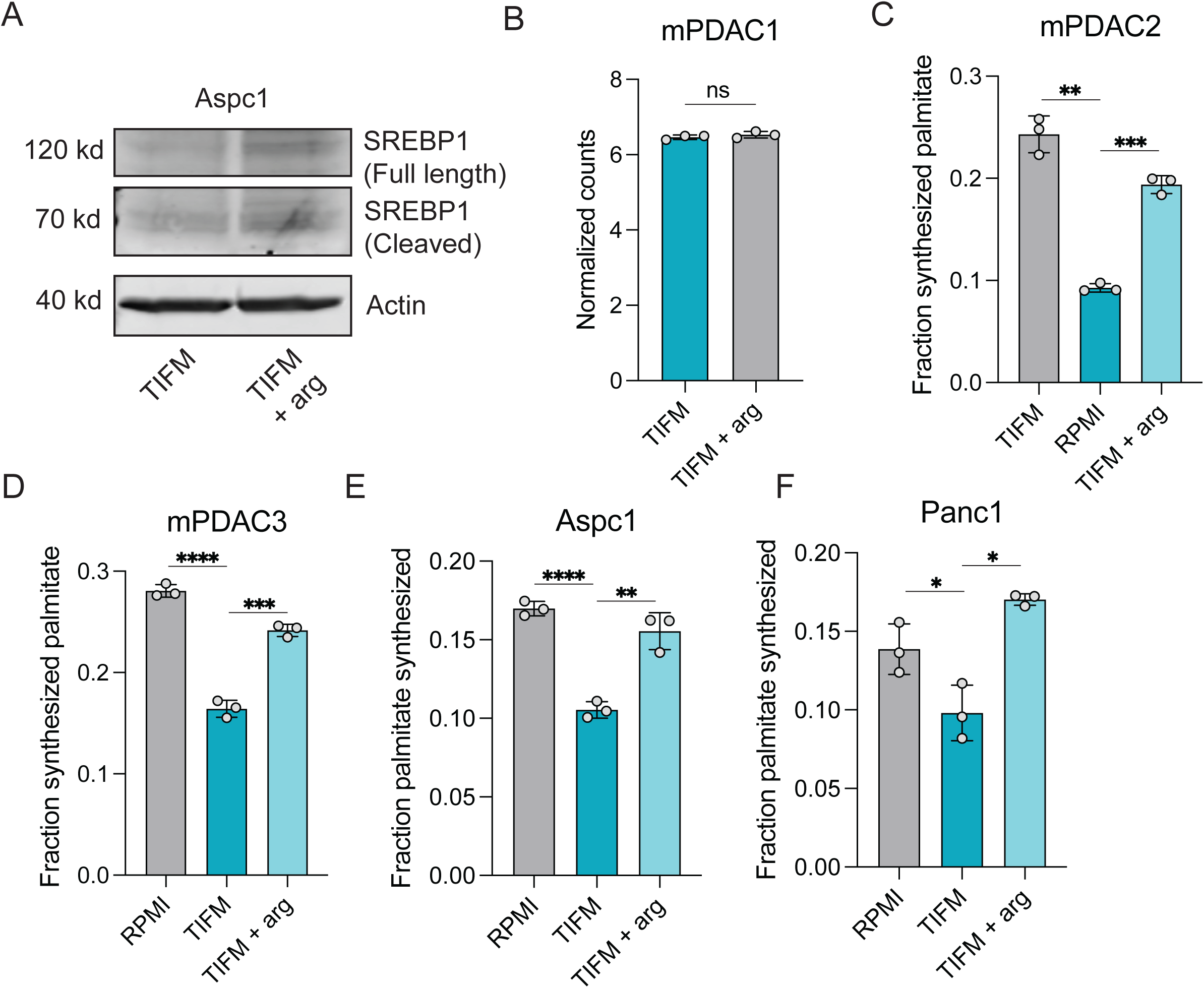
SREBP1 activity is rescued by arginine supplementation. **(A)** Immunoblot for SREBP1 in Aspc1 cells grown in TIFM or TIFM with supplemented arginine. **(B)** TMM normalized counts from data analyzing transcriptomic changes between cells grown in TIFM or TIFM with supplemented arginine (n=3). **(C-F)** Fraction synthesized palmitate in mPDAC2, mPDAC3, Aspc1, and Panc1 cells grown in TIFM or TIFM with supplemented arginine (n=3). Significance was determined using a two-tailed Welch’s t-test.

**Figure 2 – figure supplement 3:**
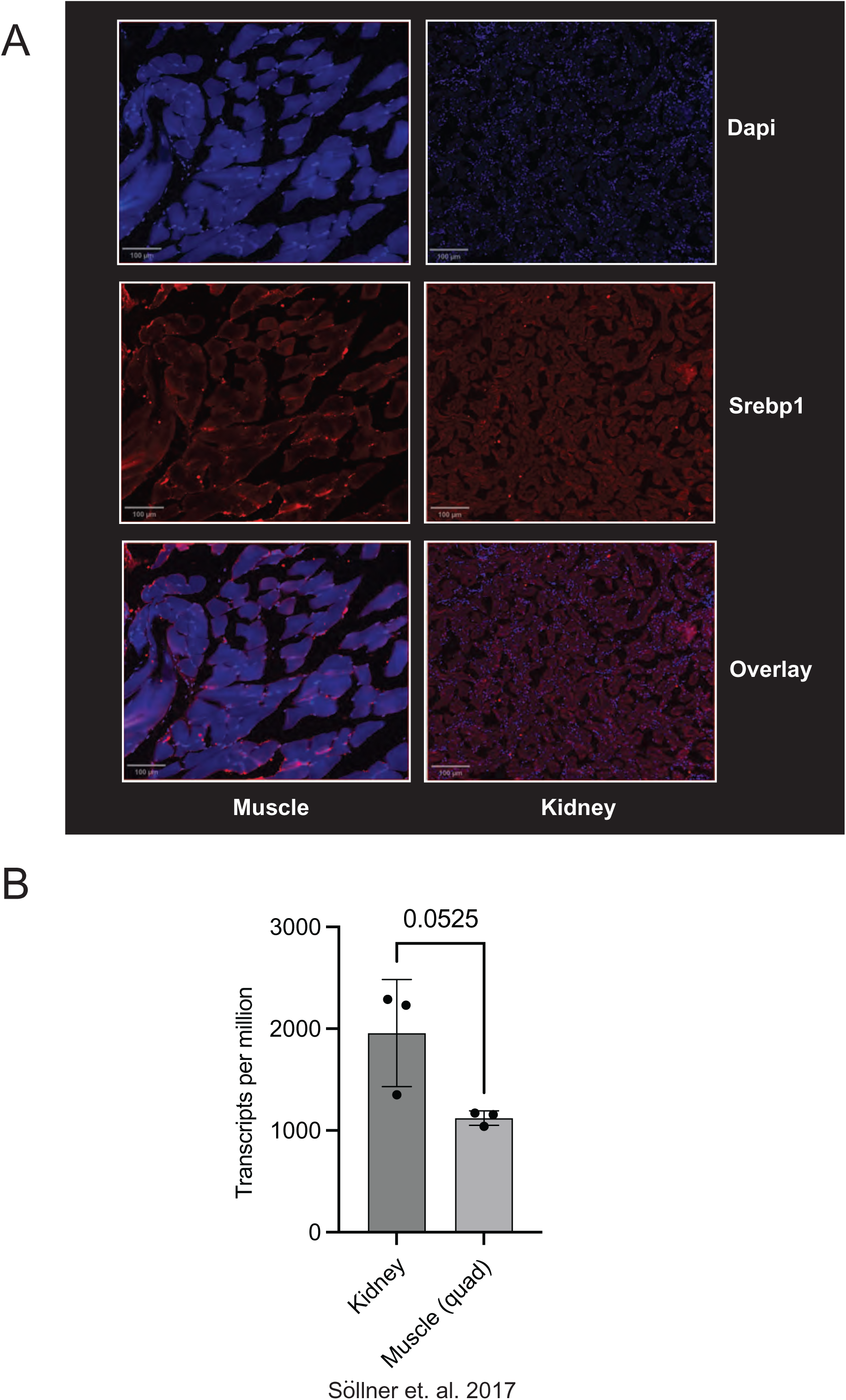
SREBP1 immunofluorescent signal reflects the expression of SREBP1 in tissues. **(A)** Murine kidney and muscle sections were stained with SREBP1 antibody as performed with murine PDAC samples in Figure 2. **(B)** Expression levels of SREBP1 mRNA in murine muscle vs kidney from reference ^147^.

**Figure 3 – figure supplement 1:**
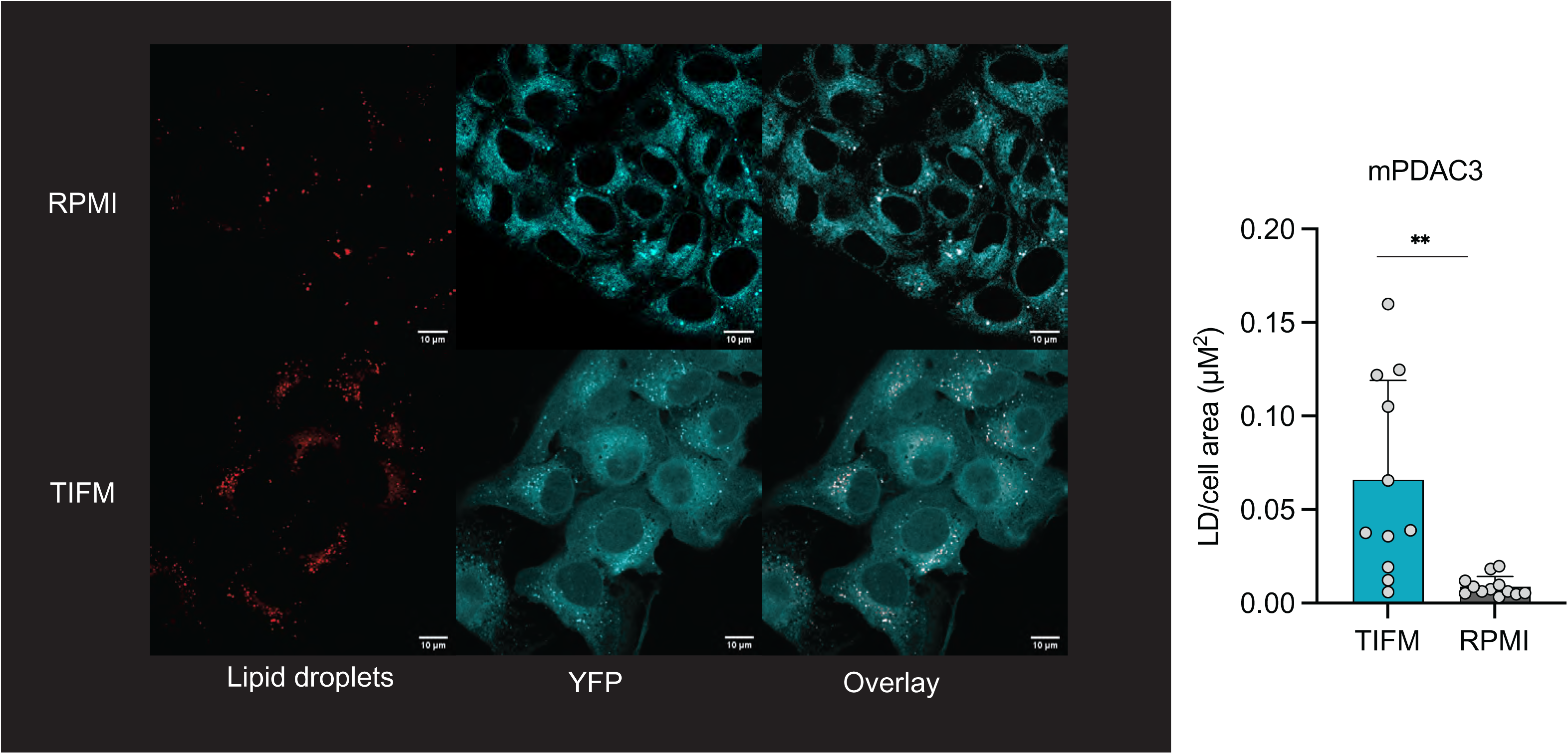
TIFM cultured PDAC cells have more lipid droplets than PDAC cells in standard culture. **(A)** Immunofluorescent analysis of YFP-expressing mPDAC3 stained with LipidTox to label lipid droplets. **(B)** Quantification of lipid droplets per cell area from **(A)**. Significance was assessed in B using a two-tailed Welch’s t-test.

**Figure 3 – figure supplement 2:**
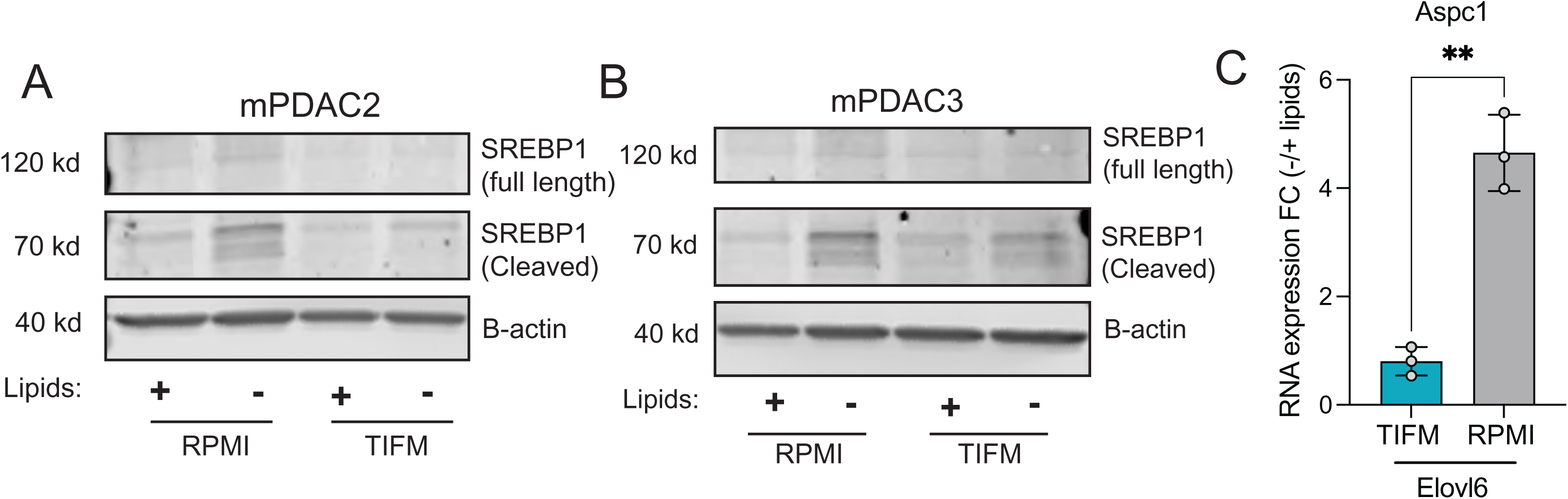
SREBP1 activation is impaired in TIFM-cultured PDAC cells. Immunoblot analysis of full length and cleaved SREBP1 expression in **(A)** mPDAC2 and **(B)** mPDAC3 cells grown in TIFM or RPMI with and without exogenous lipids. **(C)** Fold change in expression of ELOVL6 mRNA, an SREBP1 target, in mPDAC1-TIFM and mPDAC1-RPMI cells after deprivation of exogenous lipids (n=3).

**Figure 3 – figure supplement 3:**
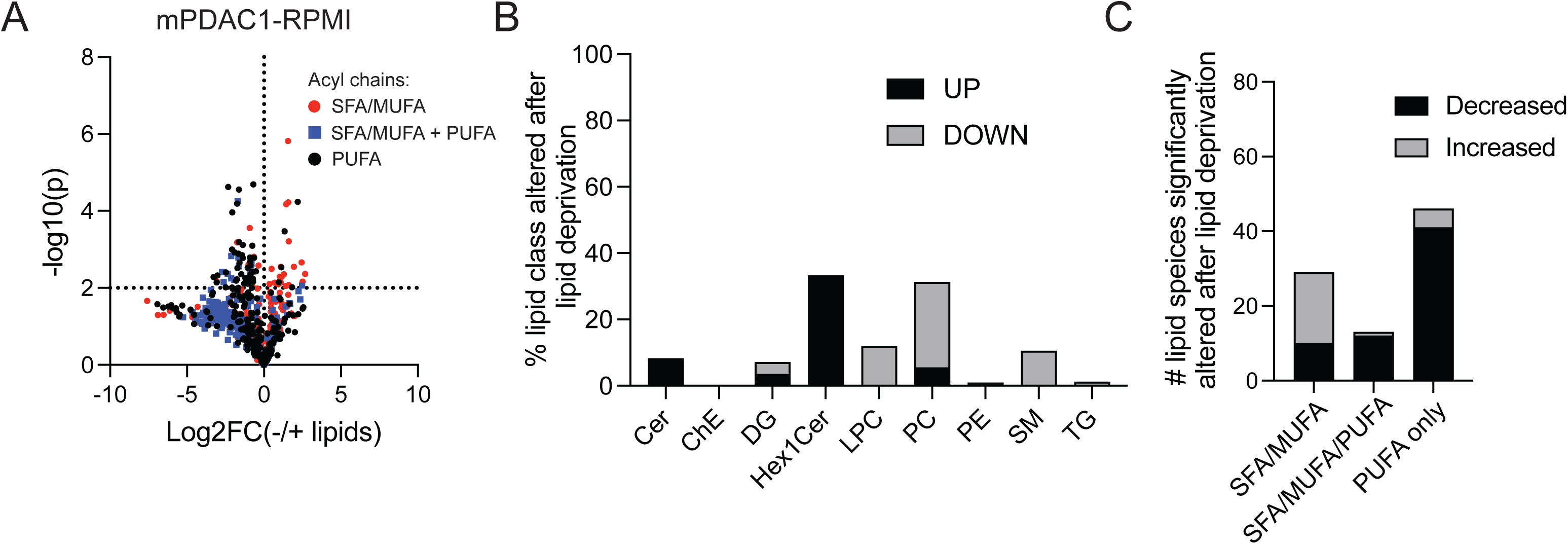
PDAC cells in RPMI can maintain lipid homeostasis upon starvation of exogenous lipids. **(A)** Volcano plots depicting the log2 fold change in levels of lipids in mPDAC1 cells upon starvation of exogenous lipids when cultured in RPMI. Lipids containing only SFA/MUFA acyl groups are highlighted in red. Lipids containing both an SFA or MUFA and PUFA acyl group are highlighted in blue. Lipids containing only PUFA acyl chains are shown in black. The cut off for statistical significance in this analysis is p<0.01. **(B)** Quantification of the percent of an indicate lipid class that is altered upon lipid deprivation in RPMI cultured mPDAC1 cells. **(C)** Number of lipid species with indicated composition of acyl groups whose cellular abundance is significantly altered upon lipid withdrawal in RPMI cultured mPDAC1 cells.

**Figure 3 – figure supplement 4:**
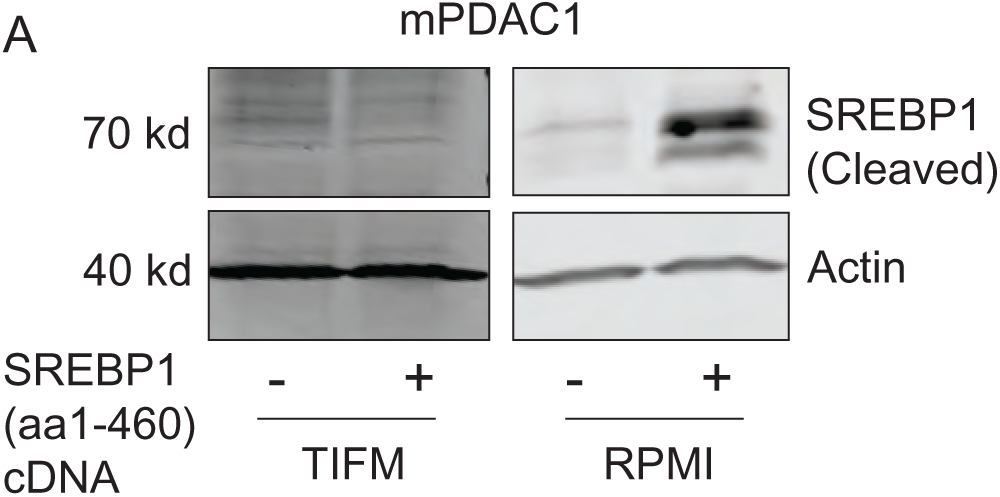
TIFM-cultured cells are unable to overexpress mature SREBP1. **(A)** Immunoblot analysis of cleaved SREBP1 expression in mPDAC1 cells cultured in TIFM or RPMI after transduction with cleaved SREBP1 cDNA.

**Figure 5 – figure supplement 1:**
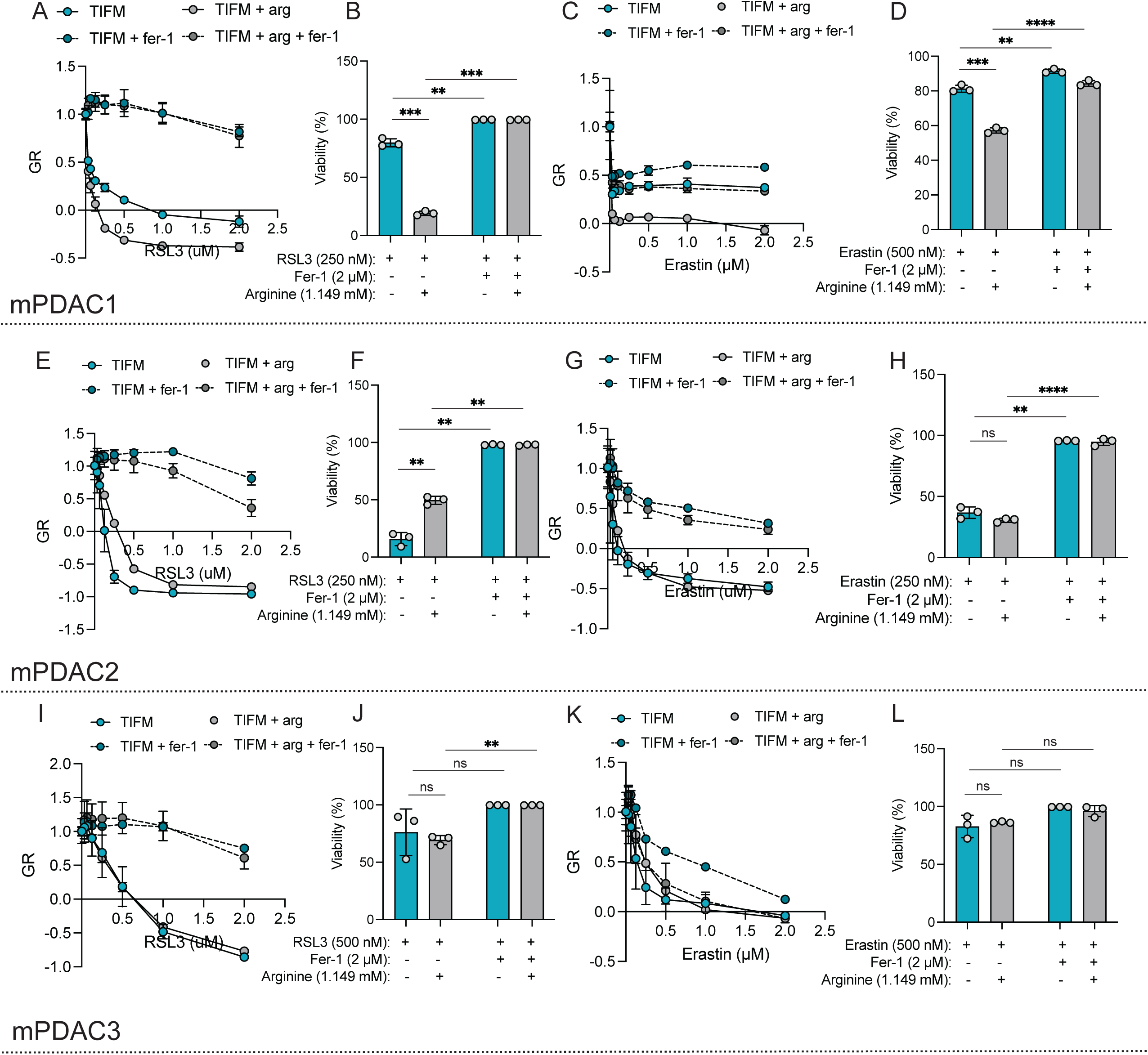
Arginine starvation does not sensitize mPDAC cells to GPX4 or xCT inhibitors. (A, C, E, G, I, **K)** GR values and **(B, D, F, H, J, L)** viability of mPDAC1, mPDAC2, and mPDAC3 cells cultured in TIFM or TIFM + arg upon treatment with RSL3 and erastin at indicated concentrations GPX4 or erastin. Ferrostatin-1 (fer-1, 2 µM) or vehicle (DMSO) was additionally added. Statistical significance was assessed in all panels by two-tailed Welch’s t-tests.

**Figure 5 – figure supplement 2:**
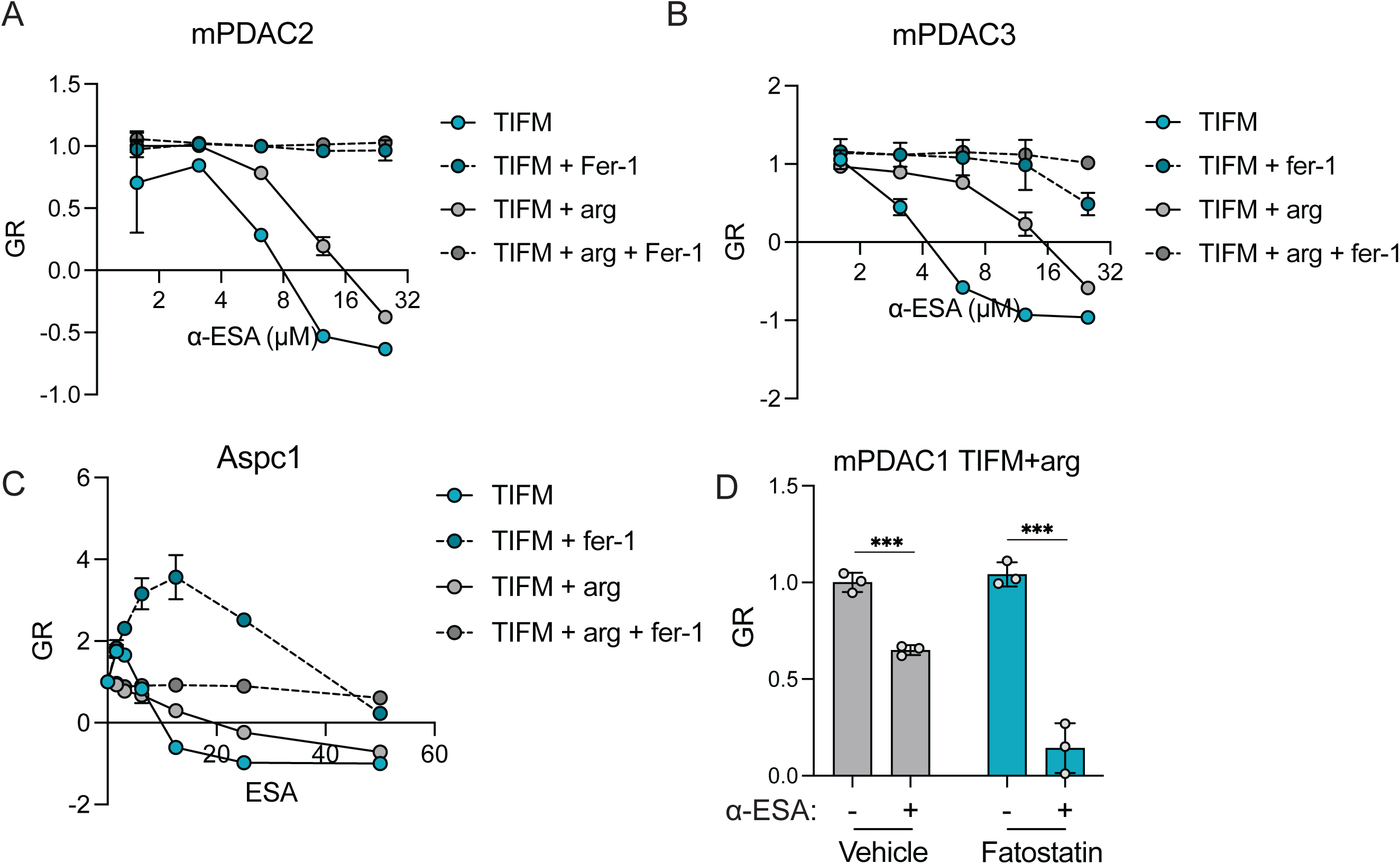
Arginine starvation sensitizes murine and human PDAC cells to exogenous α-ESA. GR values of **(A)** mPDAC2, **(B)** mPDAC3 and **(C)** Aspc1 cells cultured in TIFM or TIFM + arg with indicated concentrations of α-ESA. Ferrostatin-1 (fer-1, 2 µM) or vehicle (DMSO) was additionally added. **(D)** GR values of mPDAC1 cells grown in TIFM with supplemented arginine, with or without fatostatin treatment (2.5 uM) and α-ESA treatment (6.25 uM).

**Figure 5 – figure supplement 3:**
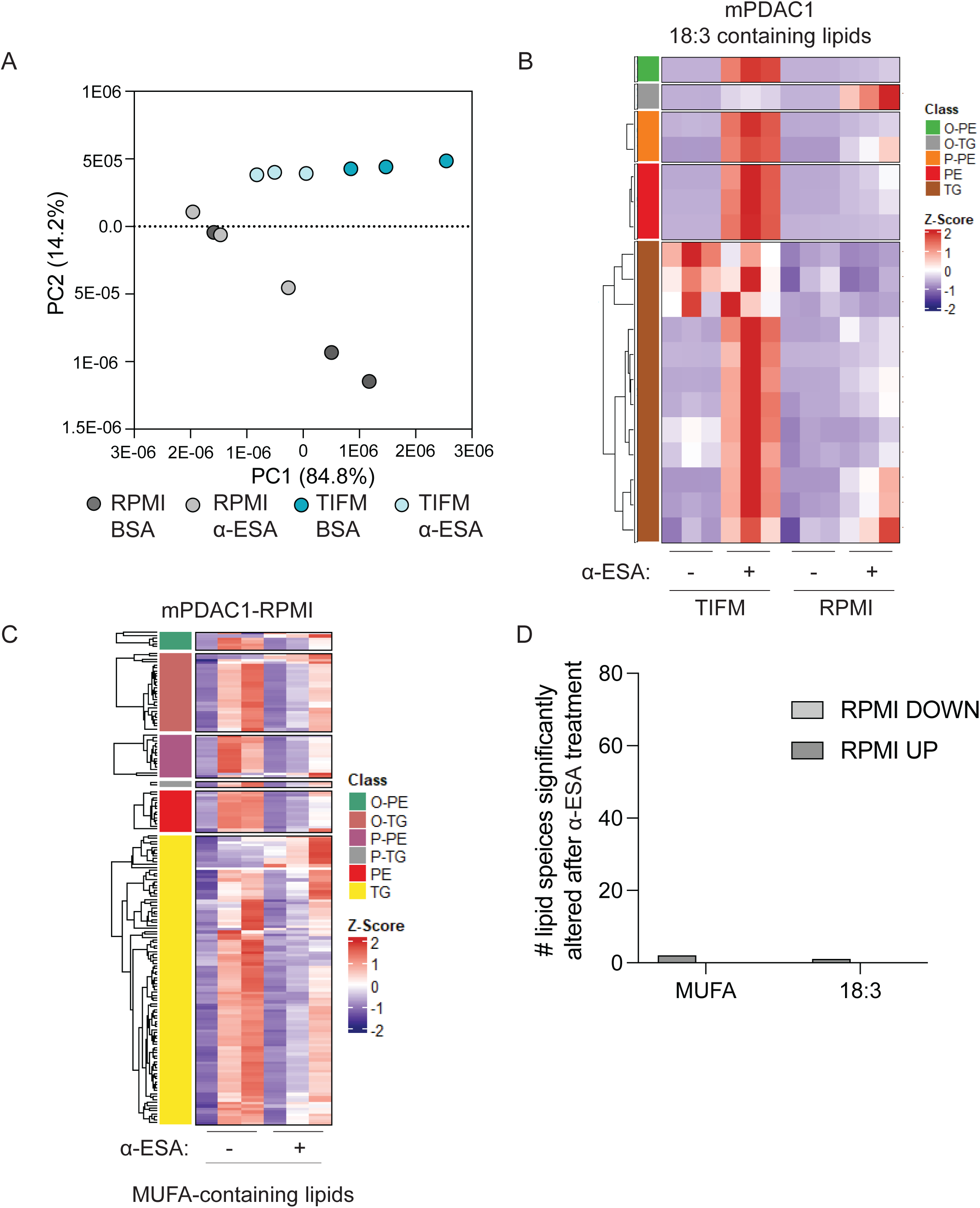
α-ESA supplementation of RPMI-cultured PDAC cells has minimal effects on the cellular lipidome. **(A)** Principal component analysis of LC-MS measurements of lipid levels of mPDAC1 cells cultured in RPMI with α-ESA (50 µM) or vehicle (fatty acid free BSA) (n=3). **(B)** Heatmap of TG and PE species with 18:3 acyl chains in mPDAC1 cells cultured as indicated upon treatment with α-ESA (50 µM). Plotted values are Z-scores for each lipid species (n=3). **(C)** Heatmaps of MUFA-containing PE and TG species in mPDAC1 cells cultured as indicated upon treatment with α-ESA (50 µM). Plotted values are Z-scores for each lipid species (n=3). **(D)** Analysis of the number of significantly altered lipid species containing MUFA or 18:3 acyl groups in mPDAC1 cells cultured as indicated upon treatment with α-ESA (50 µM). Significance cutoff p<0.05.

**Figure 6 – figure supplement 1:**
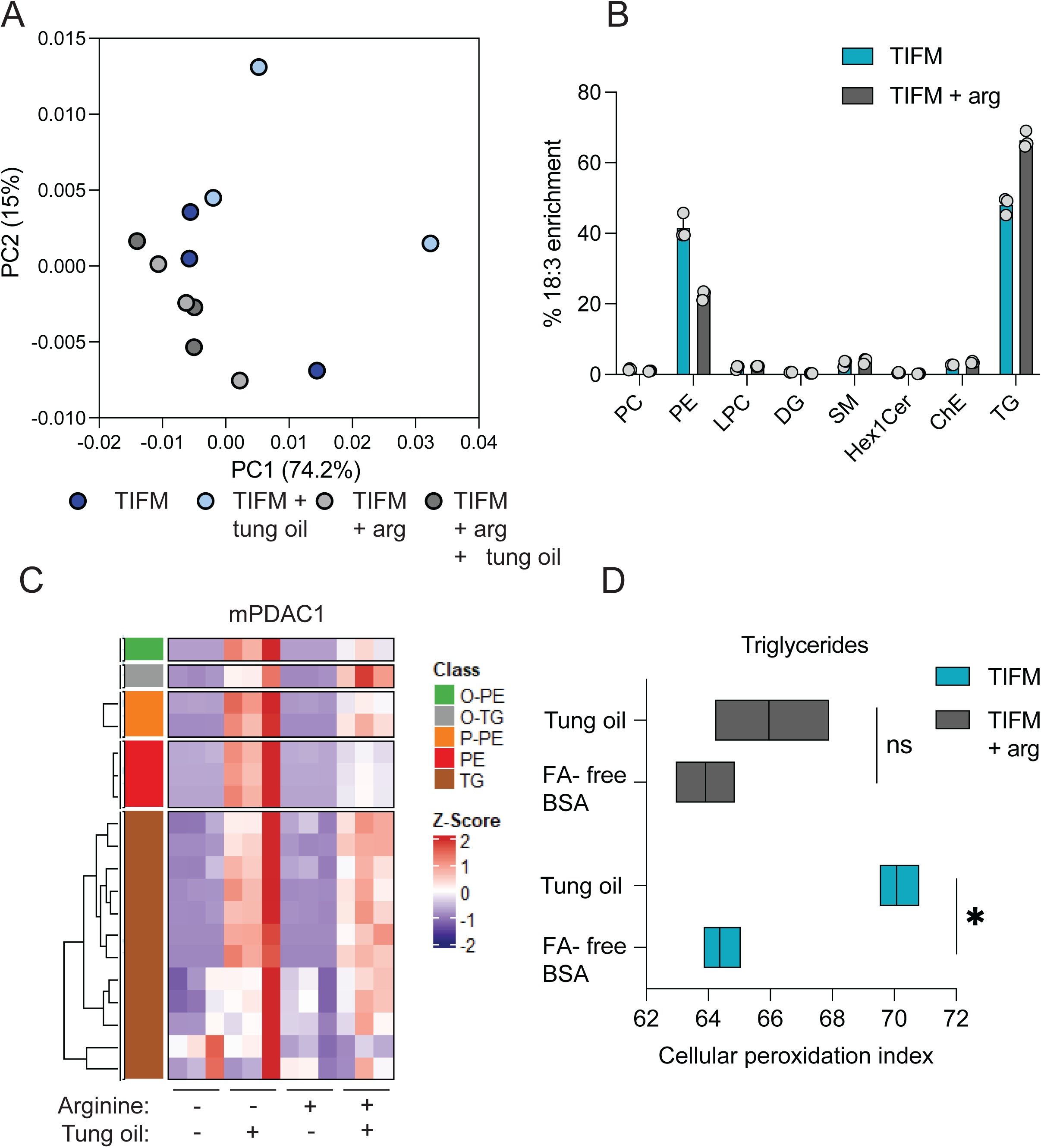
PUFA-rich oils alter the lipidomic profile of mPDAC1-TIFM cells in an arginine dependent manner. **(A)** Principal component analysis of LC-MS measurements of lipid levels of mPDAC1 cells cultured in TIFM or TIFM + arg with BSA-tung oil (200 µg/mL) or vehicle (fatty acid free BSA) (n=3). **(B)** Distribution of 18:3 fatty acyl groups in different complex lipid species in BSA-tung oil (200 µg/mL) or vehicle (fatty acid free BSA) treated mPDAC1 cells cultured in TIFM or TIFM + arg. **(C)** Heatmap of TG and PE species with 18:3 acyl chains in mPDAC1 cells cultured as indicated upon treatment with BSA-tung oil (200 µg/mL) or vehicle (fatty acid free BSA). Plotted values are Z-scores for each lipid species (n=3). **(D)** Calculated cellular phospholipid peroxidation index (CPI) for TGs in TIFM and TIFM + arg cultured mPDAC1 cells treated with BSA-tung oil (200 µg/mL) or vehicle (fatty acid free BSA) (n=3). Significance assessed using two-tailed Welch’s t-test.

**Figure 7 – figure supplement 1:**
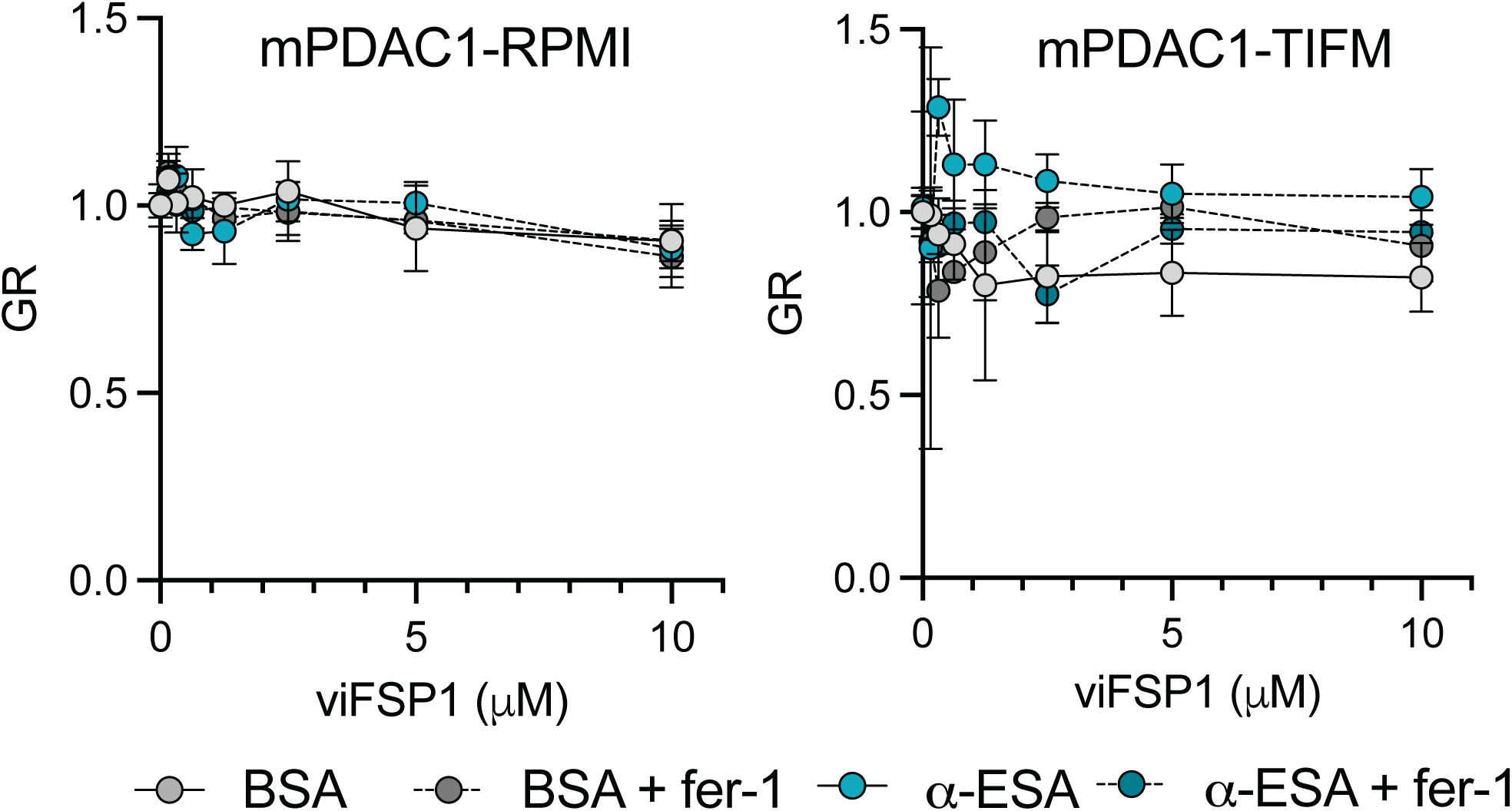
mPDAC1 cells are not sensitive to FSP1 inhibition. **(A)** GR values of mPDAC1 cells treated with α-ESA (2 µM) of vehicle (fatty acid free BSA) in tandem with viFSP1 at indicated concentrations. Ferrostatin-1 (fer-1, 2 µM) or vehicle (DMSO) was additionally added.

