## Supplementary figures and images for "Microenvironmental arginine restriction sensitizes pancreatic cancers to polyunsaturated fatty acids by suppression of lipid synthesis"

### Figure 1 - source data 4.pdf

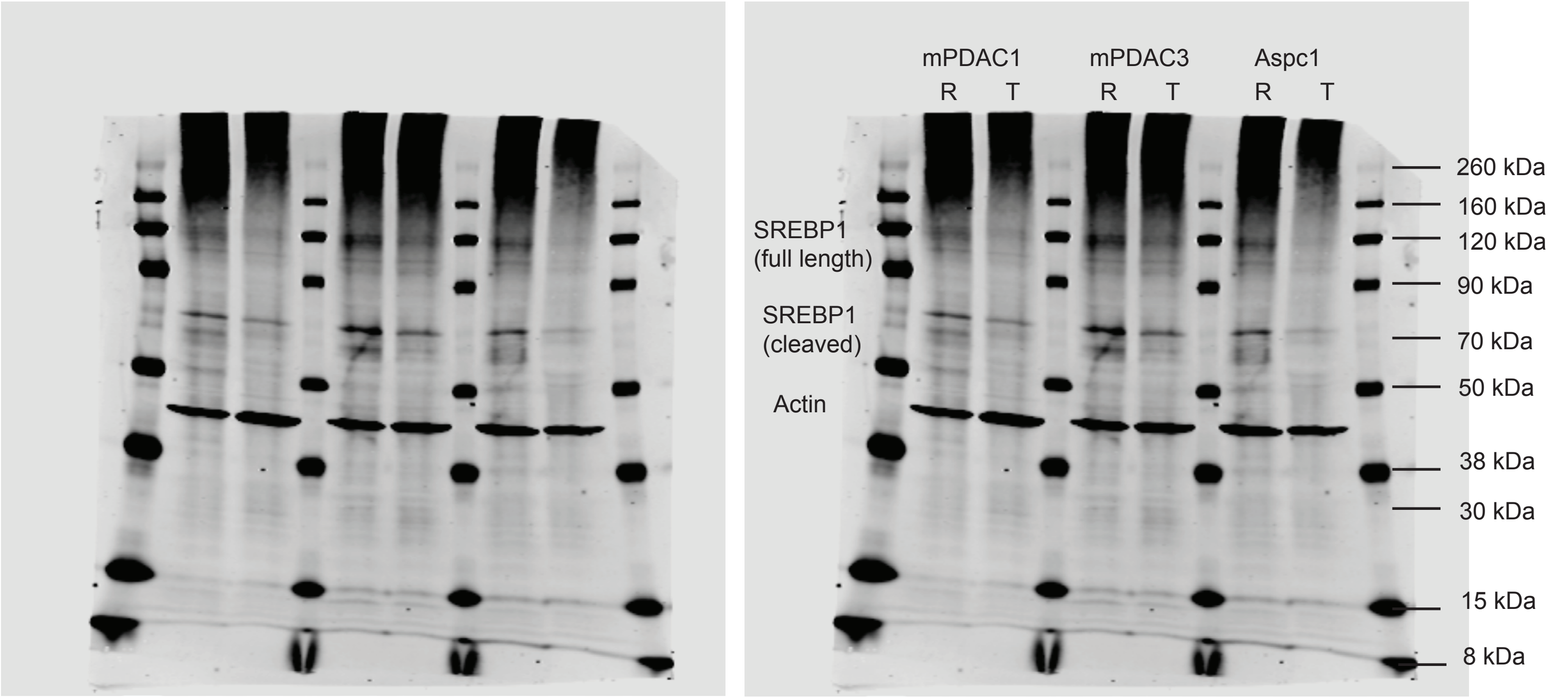

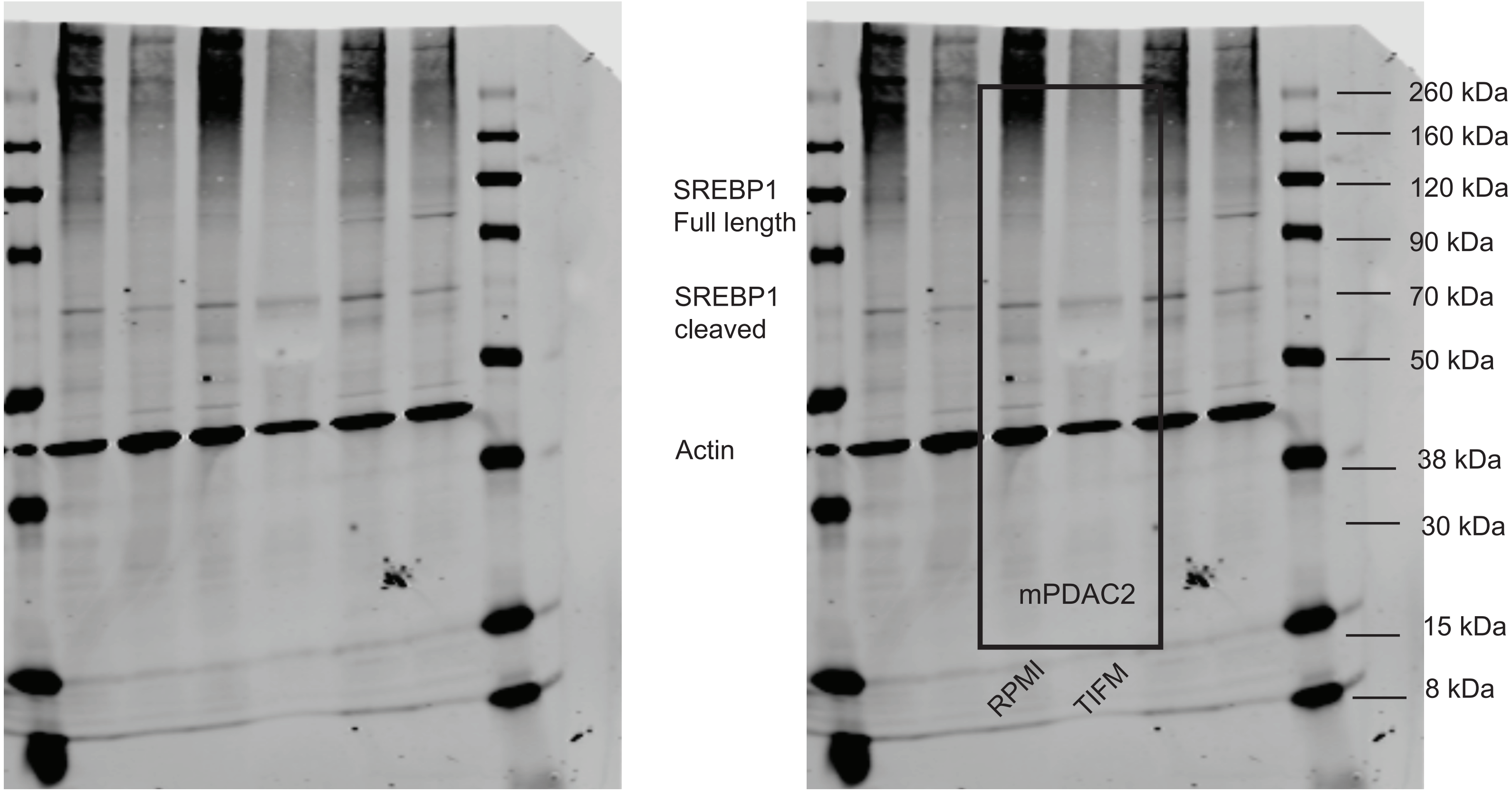

### Figure 2 - Source data 4.pdf

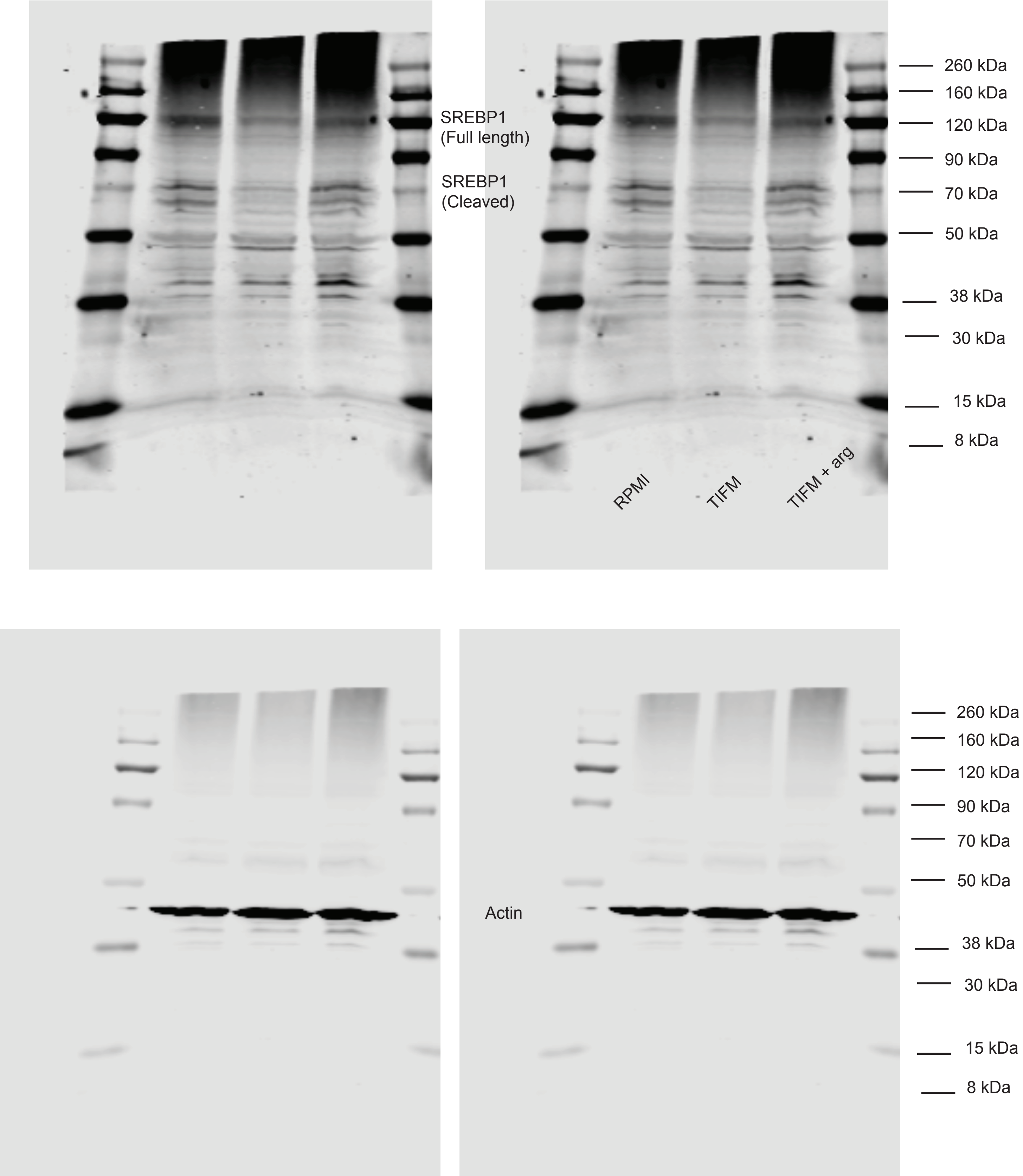

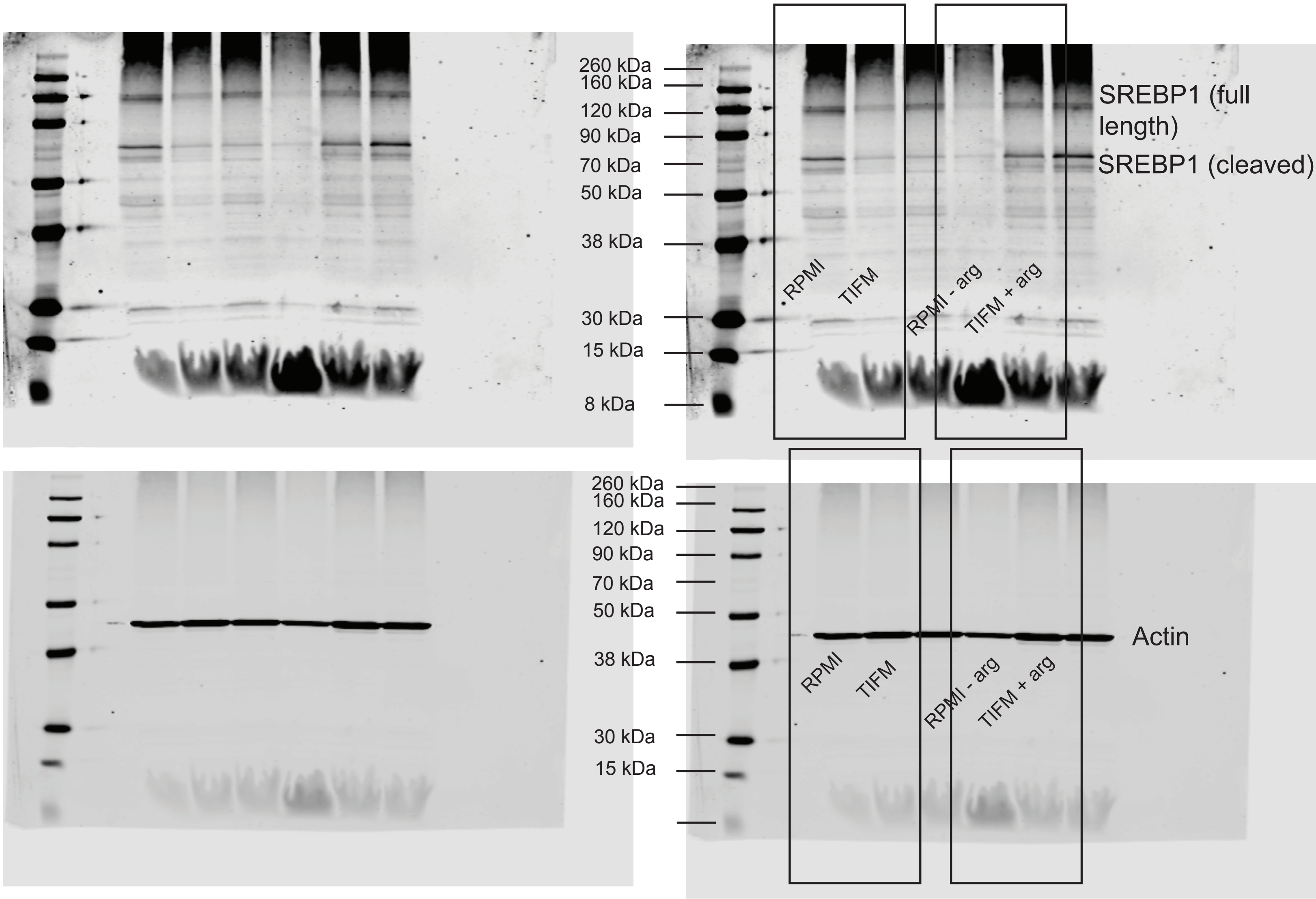

### Figure 3 - source data 3.pdf

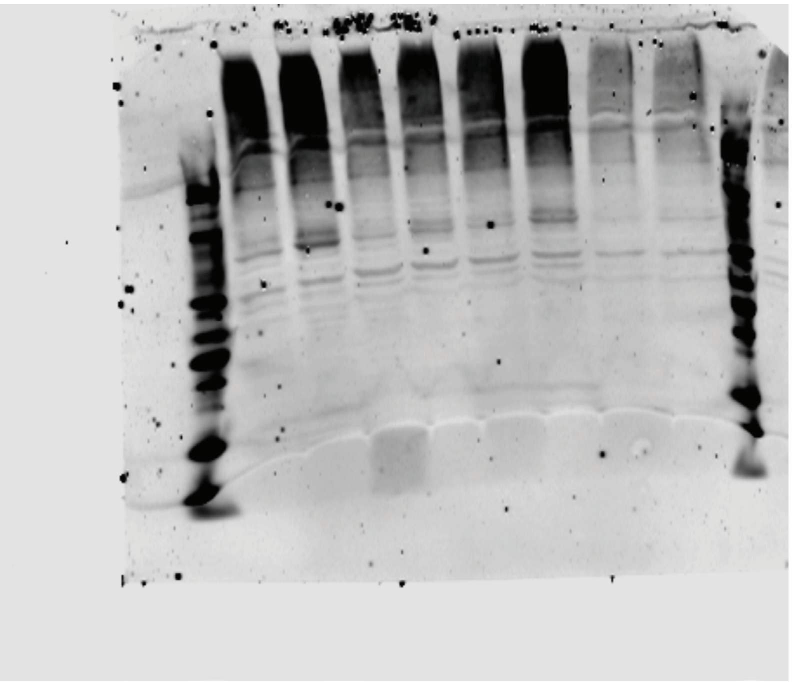

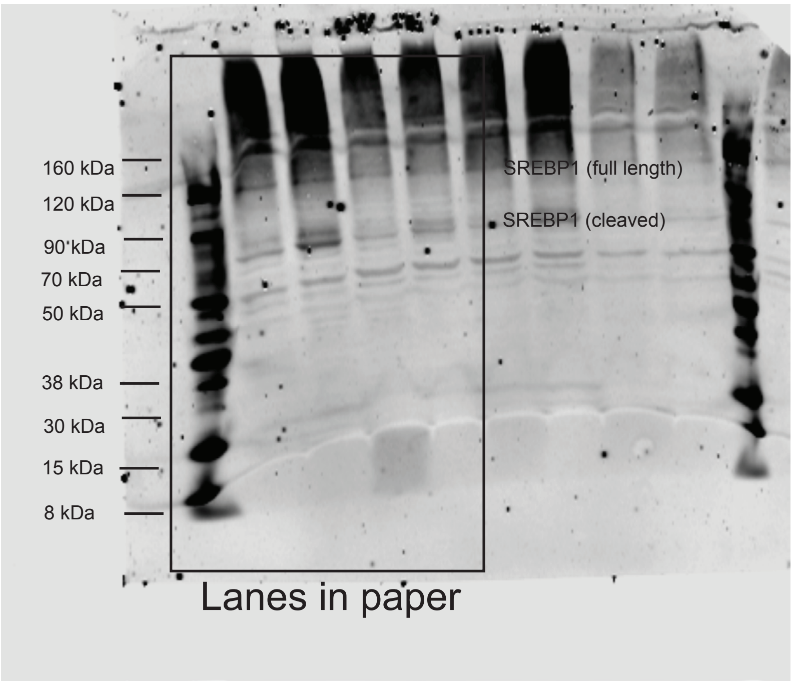

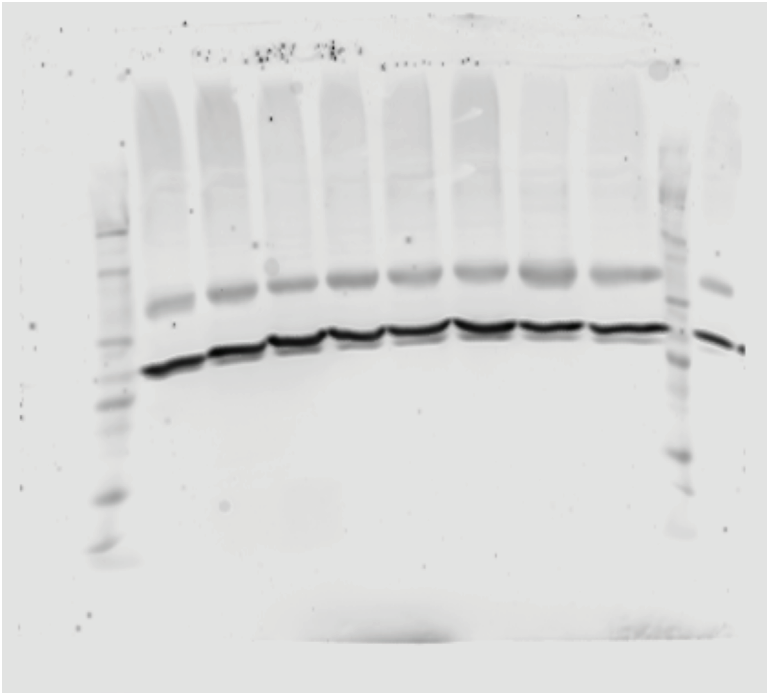

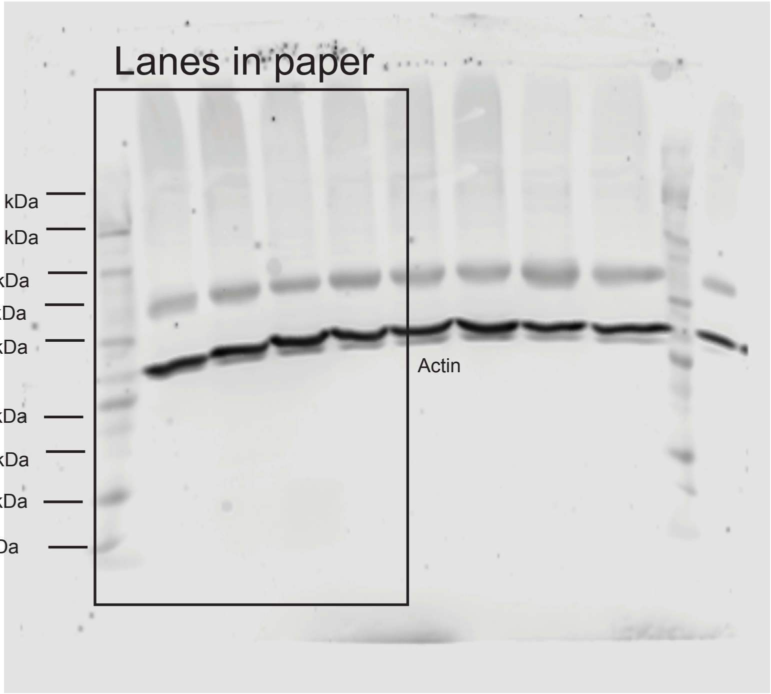

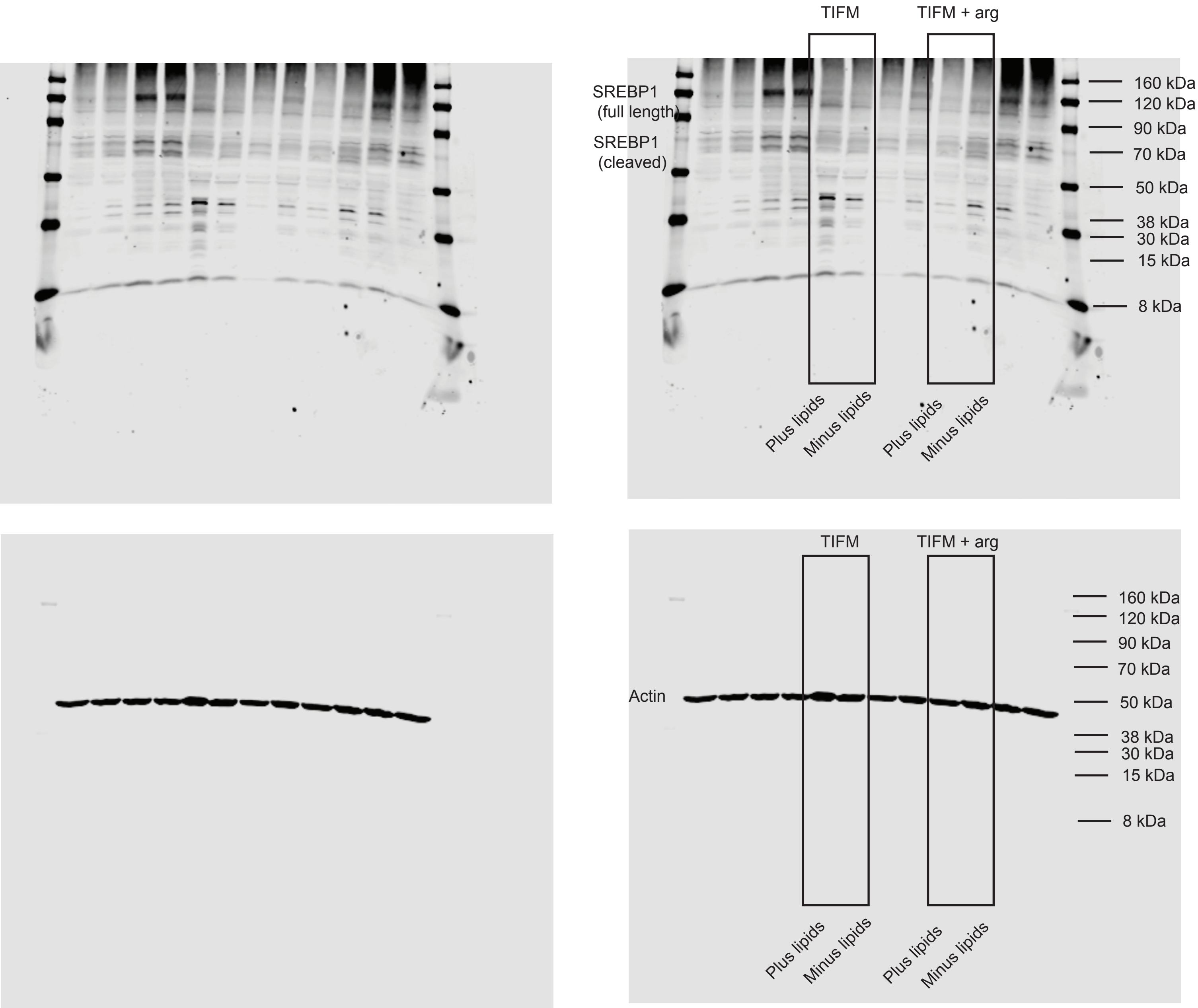

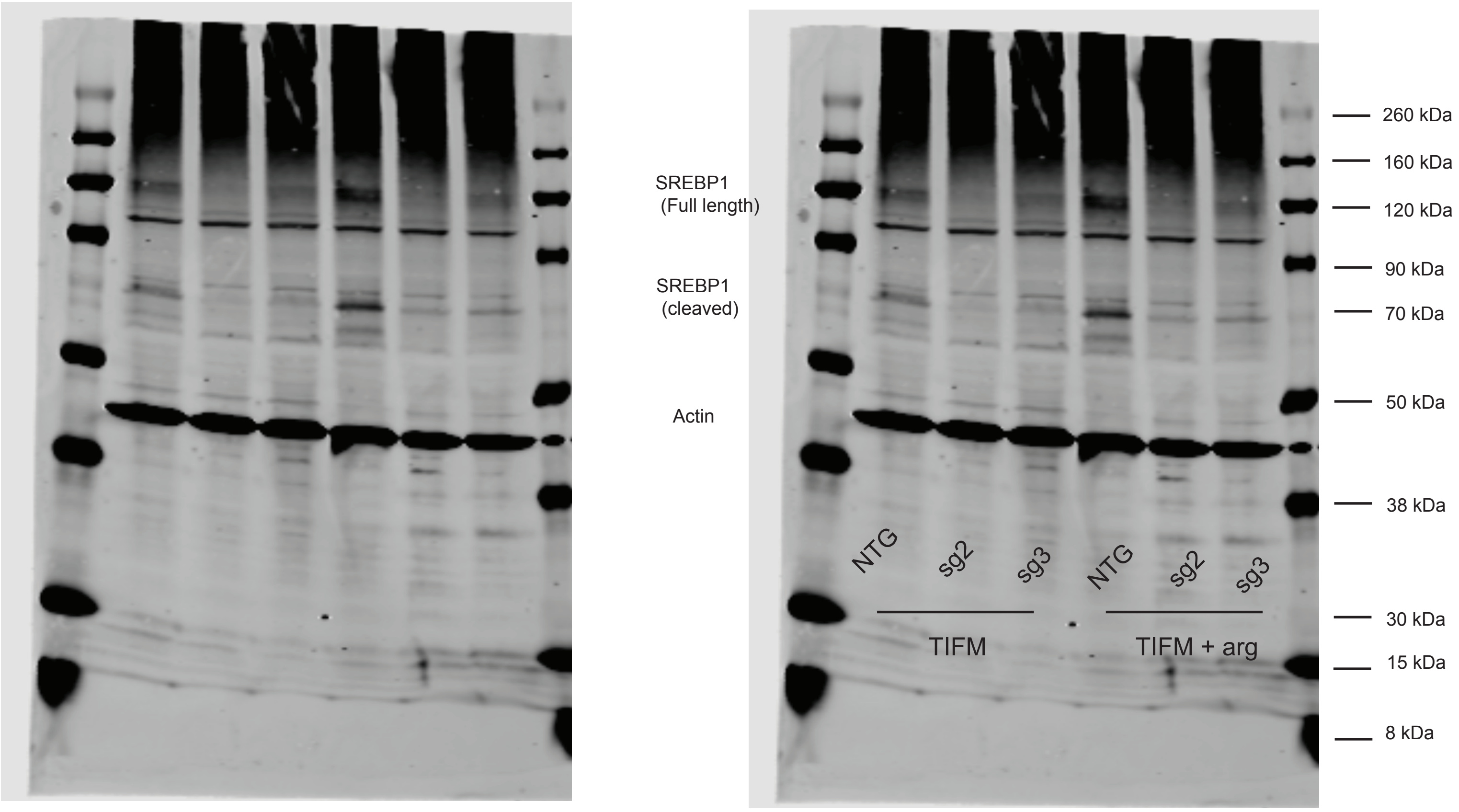


Sg2/3 targeting SREBP1

### Figure 3 - source data 4.pdf

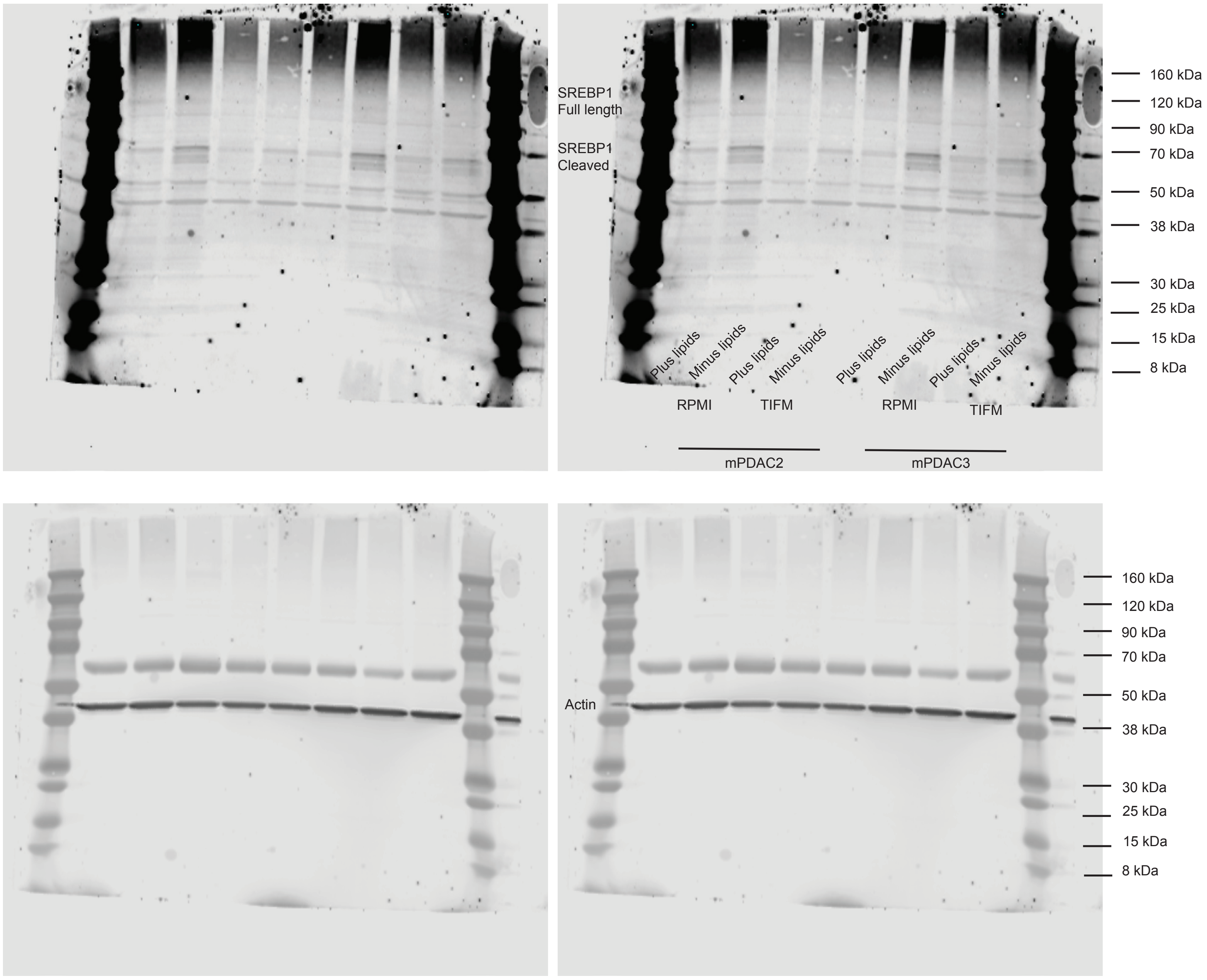

### Figure 3 - source data 5 real.pdf

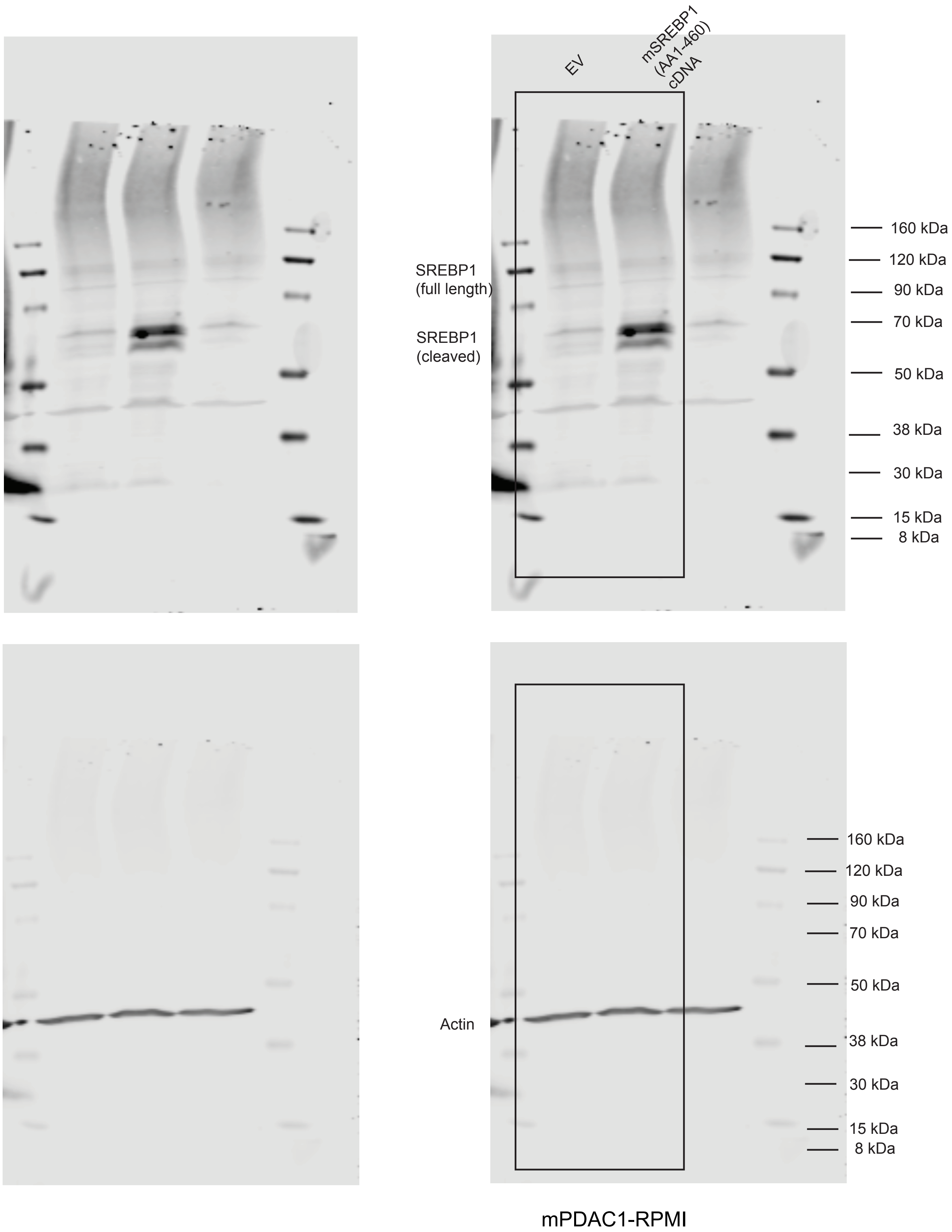

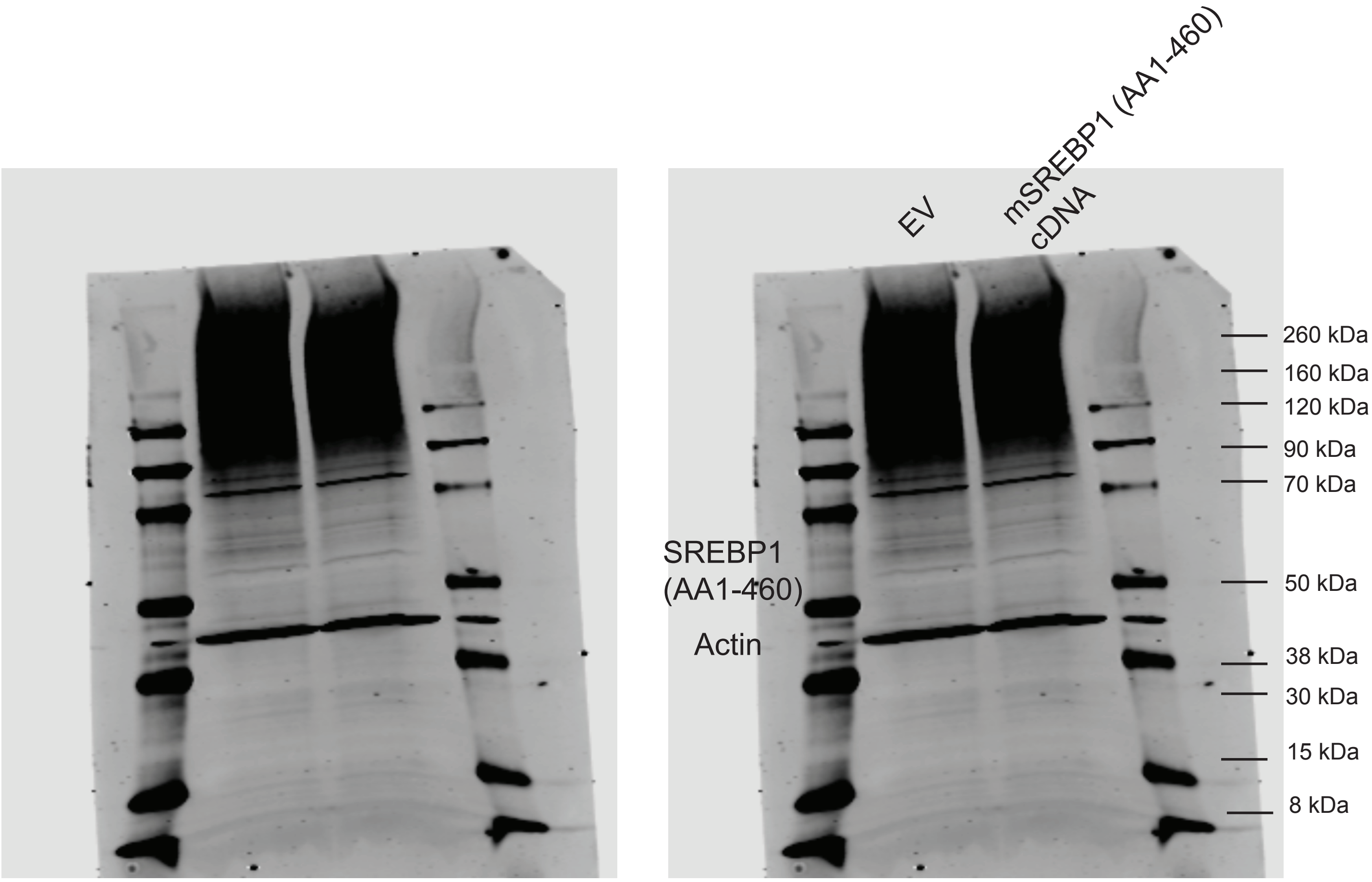

### Figure 4 Source data 1.pdf

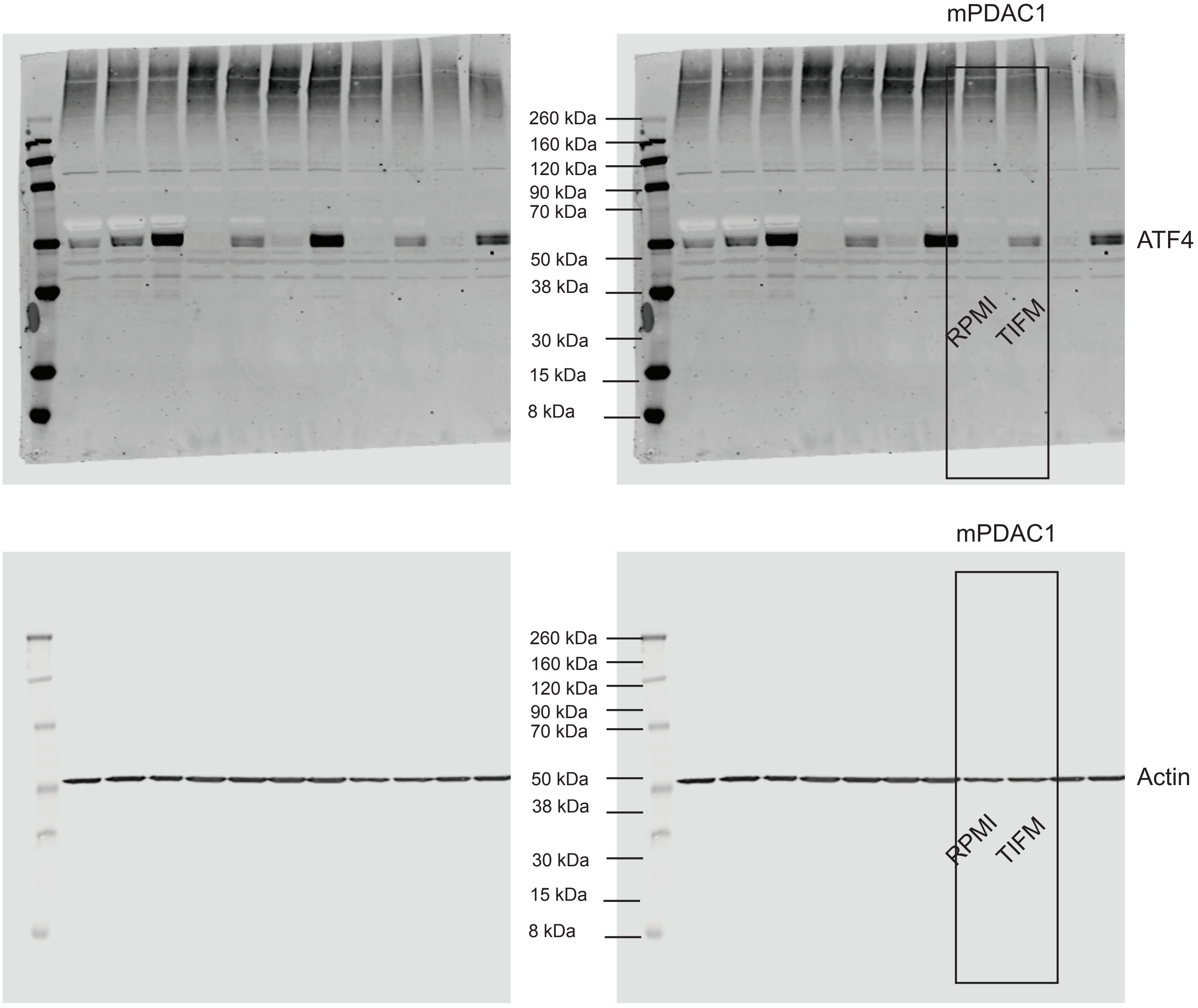

### Figure 4 Source Data 2.pdf

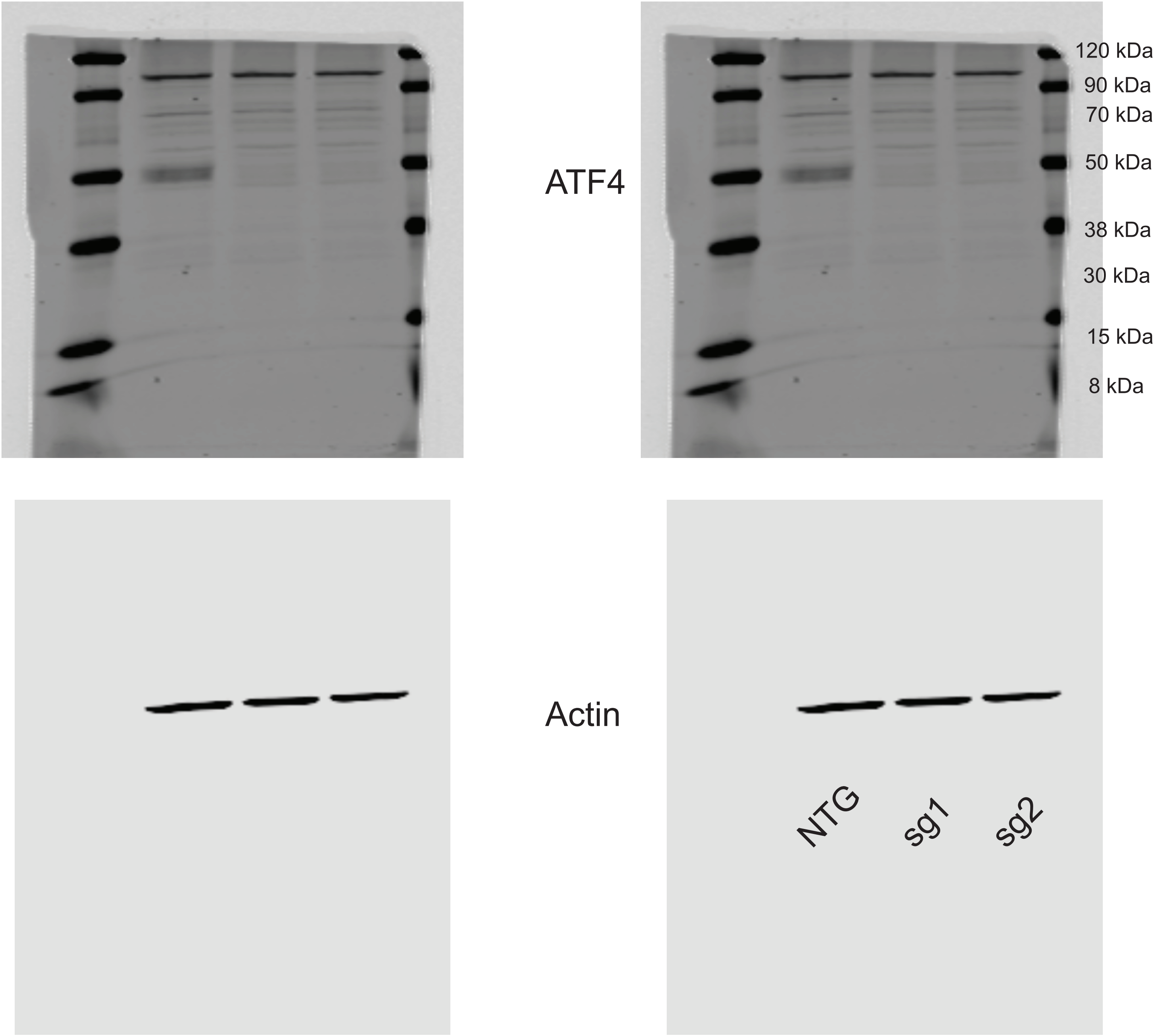

### Figure 4 Source data 3.pdf

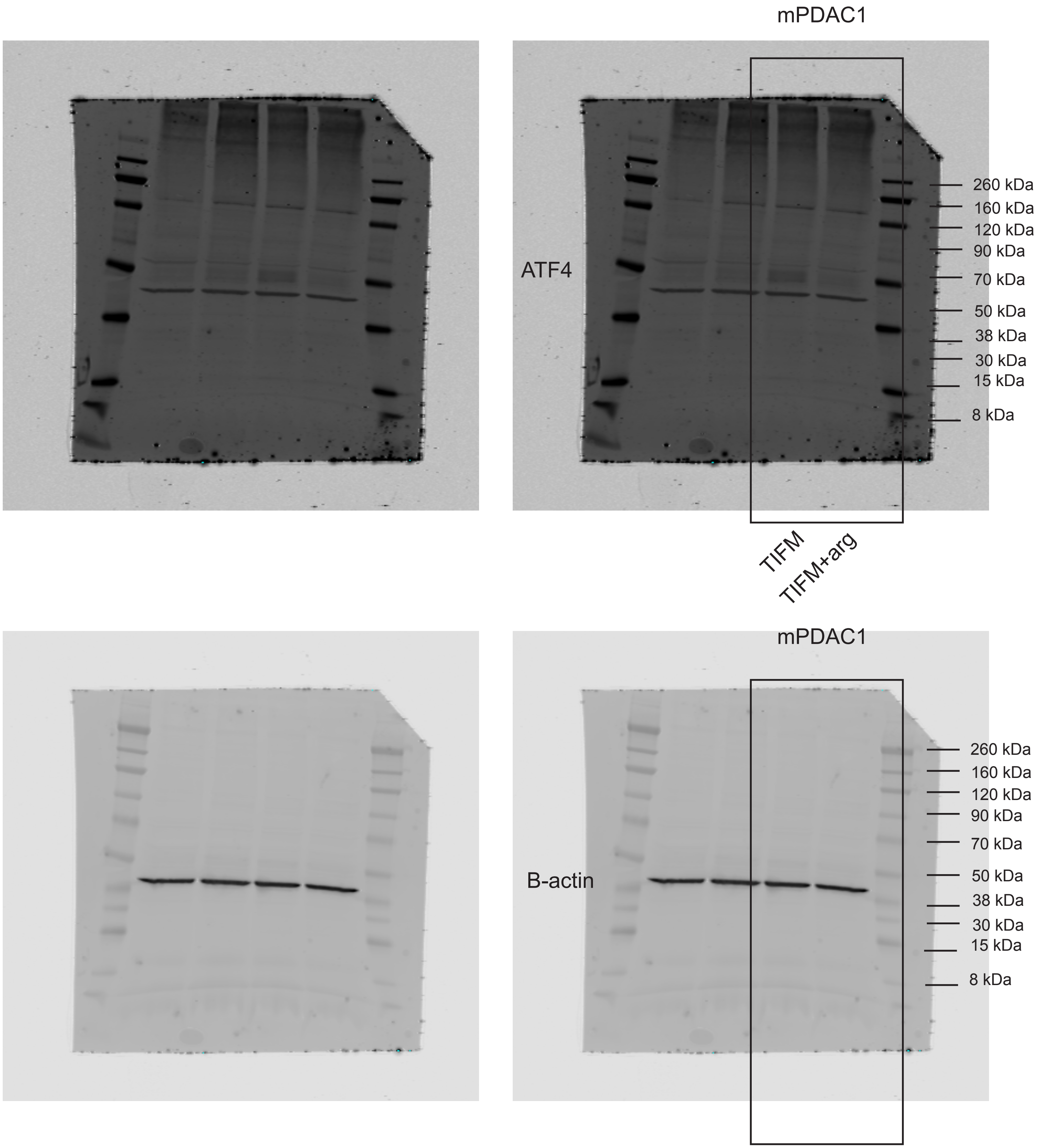

### Figure 4 Source Data 4.pdf

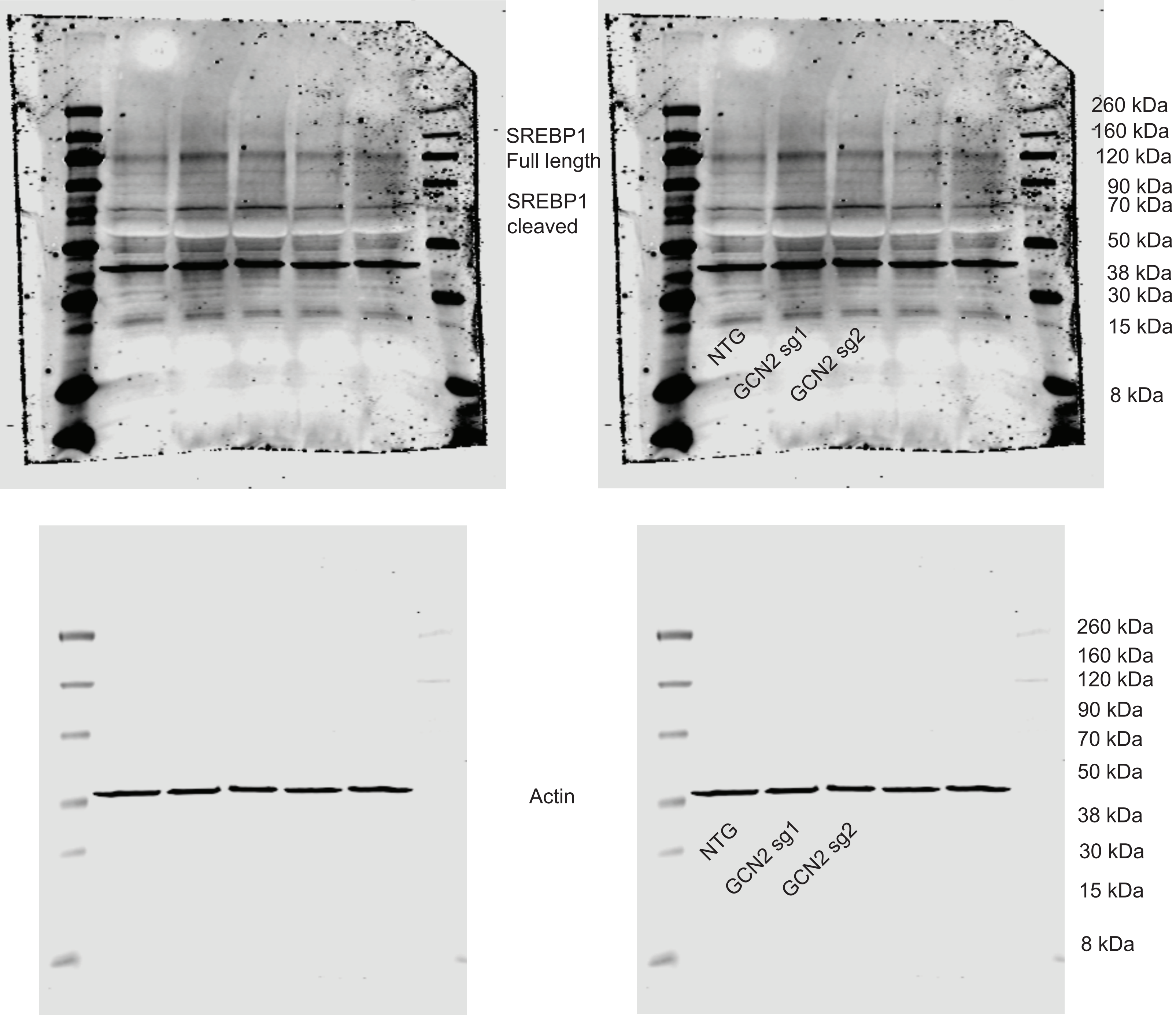
